# Integration of aged brain multi-omics reveals cross-system mechanisms underlying Alzheimer’s disease heterogeneity

**DOI:** 10.1101/2025.09.23.678110

**Authors:** Lucas P Scheidemantel, Katia de Paiva Lopes, Chris Gaiteri, Vilas Menon, Philip L De Jager, Julie A Schneider, Aron S Buchman, Yanling Wang, Shinya Tasaki, Roberto T Raittz, David A Bennett, Ricardo A Vialle

## Abstract

The molecular correlates of Alzheimer’s disease (AD) are increasingly being defined by omics. Yet, the findings from different data types or cohorts are often difficult to reconcile. Collecting multiple omics from the same individuals allows a comprehensive view of disease-related molecular mechanisms, while addressing conflicting findings derived from single omics. Such same-sample multi-omics can reveal, for instance, when changes observed in the transcriptome share distinct but coordinated signals in epigenetics and proteomics, relationships otherwise unclear. Here, we apply a data-driven multi-omic framework to integrate epigenomic, transcriptomic, proteomic, metabolomic, and cell-type-specific population data from up to 1,358 aged human brain samples from the Religious Orders Study (ROS) and Rush Memory and Aging Project (MAP). We demonstrate the existence of sprawling cross-omics cross-system biological factors that also relate to AD phenotypes. The strongest AD-associated factor (factor 8) involved elevated immune activity at the epigenetic level, decreased expression of heat shock genes in the transcriptome, and disrupted energy metabolism and cytoskeletal dynamics in the proteome. We also showed immune-related factors (factors 2 and 3) with discordant enrichments, reflecting reactive-like glial subpopulations and protective contributions from surveillance microglia. Both were negatively associated with AD pathology, suggesting potential immune resilience mechanisms. Finally, unsupervised clustering of participants revealed eleven molecular subtypes of the aging brain, including three clusters strongly associated with AD but displaying distinct molecular signatures and phenotypic characteristics. Our findings provide a comprehensive map of molecular mechanisms underlying AD heterogeneity, highlighting the complex role of neuroinflammatory processes, and yielding potential novel biomarkers and therapeutic targets for precision medicine approaches to AD treatment.

## INTRODUCTION

Alzheimer’s disease (AD) is a complex neurodegenerative disorder characterized by progressive cognitive decline, accumulation of β-amyloid (Aβ) plaques and neurofibrillary tangles (NFTs), neuroinflammation, and synaptic loss, significantly impacting older populations. Multifactorial conditions are known to influence the risk of sporadic late-onset form (LOAD) of the disease, including age, sex, genetic predisposition (mainly driven by *APOE* ε4 haplotype), and various lifestyle and health determinants. Despite extensive research efforts, effective treatments remain limited, in part due to the therapeutic focus predominantly on β-amyloid pathology ^1^. Numerous alternative molecular hypotheses for AD have been suggested, extending beyond β-amyloid and NFTs. Cholinergic neuron damage, oxidative stress, immune dysfunction, and neuroinflammation are some examples that emerge as potential new targets for treatment ^2–4^. Pathophysiological changes in lipid and glucose metabolism, among others, also play key roles in the onset and development of the disease ^5–8^. Still, these mechanisms are not fully elucidated, and a unified theory remains elusive. Generating comprehensive omics data from brain tissues can improve our understanding of these biological processes and may lead to early diagnosis and the development of targeted therapies.

In recent years, a vast amount of omics data has been generated to study the molecular changes in AD biology, usually focused to specific molecular levels, such as transcriptomics or proteomics ^9–12^. These efforts were pivotal in revealing and validating potential risk mechanisms not only in AD but also in other neurodegenerative diseases and led to the nomination of hundreds candidate gene targets ^13–22^. For example, analysis of coexpression networks from RNA-Seq data led to the prioritization of the module 109, a set of genes associated with AD traits and strongly linked to cognitive decline ^23^. Similarly, coexpression networks derived from proteomics data revealed the matrisome module (M42) and the MAPK signaling and metabolism module (M7), both associated with AD but not observed in RNA networks ^24^. While these studies have provided valuable insights, they are inherently limited in scope. Single-omics methods focus on isolated molecular layers and may fail to capture the interconnected complexity of the molecular framework. Studies integrating multi-omics have been proposed to overcome this limitation and have been applied in multiple contexts, including identifying molecular signatures associated with AD and suggesting better pre-symptomatic stage treatments ^25^, identifying relevant genes ^26^, suggesting molecular biomarkers ^27^, providing potential drug and therapy targets ^28^, identifying possible disease subtypes ^29^, detecting distinct molecularly differentiable disease trajectories ^30^, and translating these molecular trajectory signatures from *in vivo* biomarkers ^31^. However, most of these studies rely on feature preselection or explicit modeling of known phenotypes or disease states, which can bias interpretation and limit discovery of unexpected mechanisms.

Here, we performed an unbiased, data-driven computational approach to model multi-layered omics data from postmortem aged human brain tissues. Specifically, we applied Multi-Omics Factor Analysis (MOFA), an unsupervised latent factor model, designed to capture shared and modality-specific sources of variation across heterogeneous multi-omics datasets ^32,33^. MOFA differs fundamentally from traditional dimensionality reduction techniques like PCA in that it is generative, probabilistic, and explicitly models missing data, which is a common challenge in real-world multi-omics studies. Moreover, it supports interpretation of learned factors in terms of both contributing molecular features and sample-level variation, making it particularly well-suited to uncover biologically meaningful patterns that span across molecular modalities. In previous studies, MOFA has been used to discover novel molecular mechanisms and altered pathways associated with AD pathology ^8^, detect sex-specific metabolic alterations in the AD mouse models ^34^, and provide potential drug and therapy targets ^29^. Building on this, we applied MOFA to integrate epigenomics, transcriptomics, proteomics, metabolomics, and cell-type subpopulation data from two cohort studies of aging and dementia: The Religious Orders Study (ROS) and the Rush Memory and Aging Project (MAP). We then performed integrative pathway analyses to characterize the molecular mechanisms involved in these derived factors. Finally, we performed sample clustering based on factor scores to identify molecular subtypes of AD.

## RESULTS

### Characteristics of the participants

We utilized data from 1,358 participants in the ROS/MAP cohorts. After enrollment, participants from both cohorts were annually evaluated for physiological and cognitive function followed by brain donation at death ^35^. Of the 1,358 participants, 436 (32.1%) were male, the mean age at death was 89.4 years and the average years of education was 16.2. A total of 600 (44.2%) participants had Alzheimer’s dementia, while 318 (23.4%) had mild cognitive impairment (MCI). TDP-43 pathology extending beyond the amygdala was observed in 32.1% of participants, while Lewy bodies in the nigra, amygdala, and/or cortex regions were present in 23.2% of cases. Cerebrovascular diseases, including macroscopic infarcts and microinfarcts, were identified in 35.9% and 29.3% of cases, respectively. Additionally, moderate to severe amyloid angiopathy, atherosclerosis, and arteriolosclerosis were observed in about a third of the brains. More than a quarter of participants carried at least one *APOE* ε4 allele, and similarly, at least one long allele of TOMM40’523 (**Supplementary Table 1**).

### Multi-omics factor analysis of the aging human brain

We implemented MOFA integrating 7 different omics datasets (hereafter referred to as *views*) sourced from a partially overlapping set of brain samples, totaling 1,358 ROS/MAP participants (**Supplementary Figure S1**). These included: H3K9ac histone acetylation markers measuring 26,384 peaks (n = 632); mRNA expression of 17,309 transcripts from 3 brain regions: anterior cingulate (AC; n = 721), dorsolateral prefrontal cortex (DLPFC; n = 1,210) and posterior cingulate gyrus (PCG; n = 656); proteomics comprising 8,425 proteins (n = 596); metabolomics covering 667 metabolites (n = 500) and proportions of 96 cell-type specific subpopulations from the cortex (n = 424) (**Figure 1a**). MOFA was set to decompose the data into 50 largely orthogonal latent factors (**Supplementary Figure S2**) representing the underlying sources of variation across samples. To provide an unbiased characterization of brain molecular signatures, no pre-selection step was applied to the input features (i.e. genes, proteins, epigenetic markers, metabolites, or cell proportions). The most significant portion of total data variance was captured by mRNA (over 40% of variance explained) followed by proteomics (over 35%) (**Figure 1b**). In addition, different contribution patterns where observed from each view, for example Factors 3, 5, and 6 were mostly driven by transcriptomics from DLPFC, AC, and PCG respectively, while others views had their variance explained by at least 2 or more omics views, indicating possible more complex interplay between different molecular levels or tissues (**Figure 1c**, **Supplementary Figure S3**).

**Figure 1.**
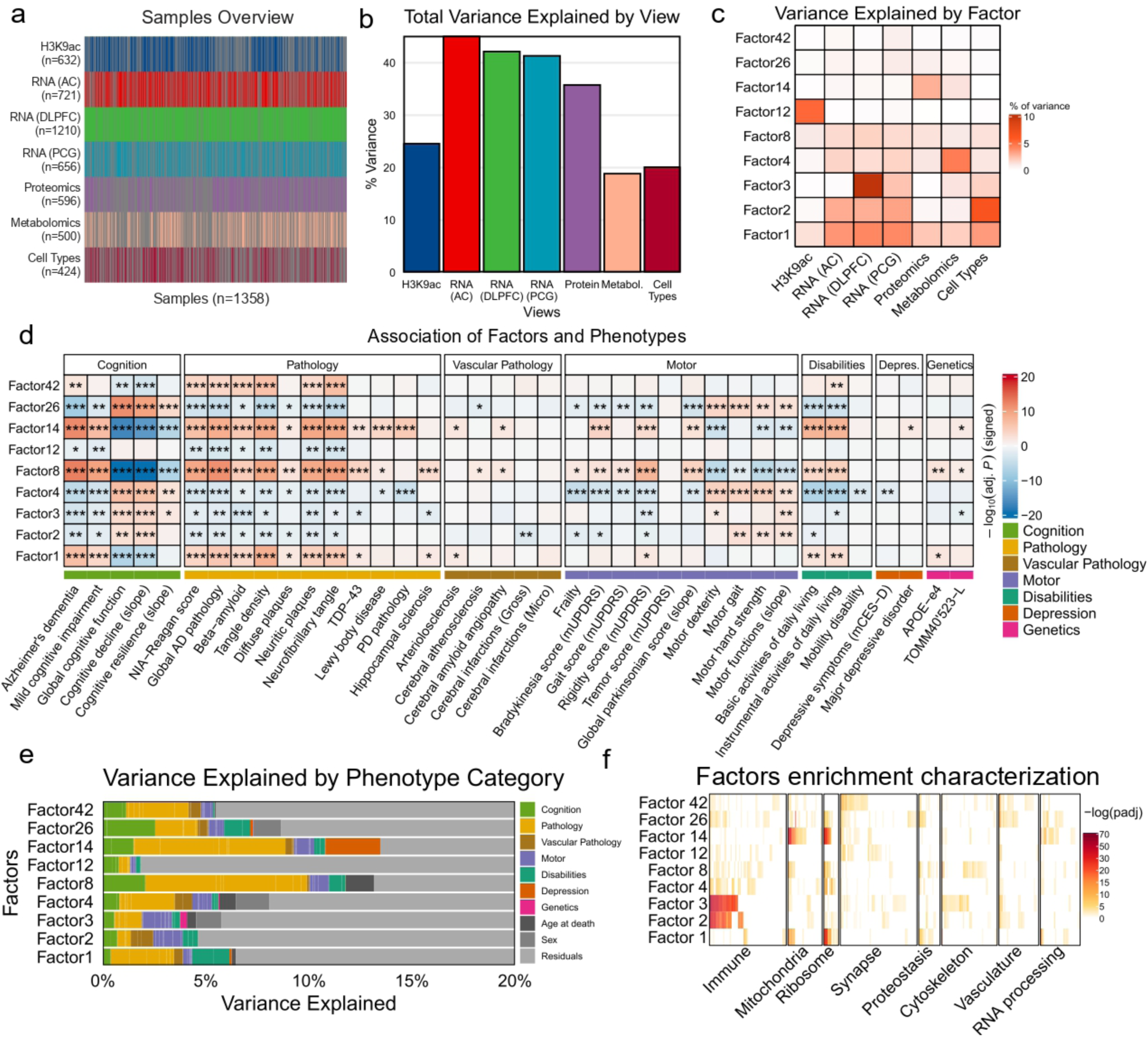
Multi-omics factor analysis of the aging human brain. **(a)** Summary of the number of samples for each *view*. Numbers in parentheses represent the total number of samples and gray lines represent the missing samples. **(b)** The sum of variance explained from all factors for each *view*. **(c)** Variance explained by factors across *views*. The color scale shows the percentage of variance explained, ranging from zero (white) to 10 (dark orange). **(d)** Relationship of factors with covariates via regression analysis. Results were controlled for sex, age, and years of education, with *P*-values corrected using the False Discovery Rate (FDR) method to address multiple comparisons. (*) = -log_10_ (adj. *P*) > 1.30103; (**) = -log_10_ (adj. *P*) > 2; (***) = -log_10_ (adj. *P*) > 3. The color scale shows the association effect size indicating direction and strength from negative (blue) to positive (red). **(e)** Variance explained for each factor by phenotype category. **(f)** Pathway enrichment analysis. Gene Set Enrichment Analysis (GSEA) was performed for each factor and the resulting terms were assigned into eight pathways. The selection of terms was made through the use of pathway related keywords. Approximately 31% of the total number of terms were allocated into one of the categories. The color scale shows the *P*-values corrected by the *Benjamini-Hochberg* (BH) method.

### Identification of AD-associated factors

From the 50 decomposed factors, we selected nine that were associated (*adj. P* < 0.05) either with pathological diagnosis of AD (NIA-Reagan score) or clinical diagnosis of Alzheimer’s dementia for downstream analysis (**Supplementary Figure S4**). These factors accounted for a range of total variance, from 20.5% (Factor 1) to 1.7% (Factor 42) (**Supplementary Figure S5**). To better characterize their phenotypic impact, we further performed association tests against 38 common age-related clinical and neuropathologic traits, including indices for cognition, pathology, motor function, disabilities, depression, and genetic risk variants for AD (**Figure 1d**, **Supplementary Tables 2 and 3**). These analyses revealed multiple associations (*adj. P* < 0.05) between factors and phenotypes, primarily with cognitive and AD pathological traits but also with motor function and disabilities. Most of these persisted after adjusting for *APOE* ε4 (**Supplementary Figure S6**). Factor 8 showed the strongest associations with cognitive decline, and was linked to 30 of the 38 tested phenotypes, followed by factors 4 and 14, associated with 27 traits each. Although many factors shared correlations across the same phenotypes (as expected due to the intrinsic correlation between phenotypes), some specific factor-trait associations emerged, suggesting that certain phenotypes might be linked to specific molecular mechanisms. For example, gross cerebral infarction was associated exclusively with factor 2, while major depressive disorder diagnosis was uniquely linked to factor 14. We also conducted a variance partition analysis to jointly assess the contribution of multiple phenotypes to each factor. This analysis revealed distinct patterns of phenotypic influence. For instance, while variance in factors 8 and 14 were mostly affected by pathologies, factor 8 had an additional influence of age at death, whereas Factor 14 showed substantial effects from depression-related phenotypes (**Figure 1e**). By analyzing the biological features driving each factor, we were able to describe the molecular functions associated with each view. All nine AD-related factors were enriched with multiple functional terms, involving key cellular mechanisms (**Figure 1f**, **Supplementary Tables 4-12**). These encompassed terms related to immune response (factors 2, 3, 4, 8 and 26); mitochondria (factors 1, 8 and 14), ribosome (factors 1, 4, 8, 14 and 26); synapse (factor 12 and 42); proteostasis (factor 1, 8 and 26); cytoskeleton (factors 3 and 8); vasculature (factors 26 and 42); and RNA processing (factors 12 and 26).

### MOFA robustness and reproducibility

To assess the robustness of the MOFA model, we compared feature weight correlations across different subsampling strategies (**Supplementary Figure S7**). First, we evaluated whether MOFA could handle missing data samples by restricting the analysis to samples with increasing numbers of omics layers. We trained MOFA models on subsets including only individuals with at least two, three, or four omics layers and compared them to the complete model (i.e. including all individuals with at least one *view*). Next, we assessed the effect of sample size by training MOFA models on random subsets of decreasing size, while keeping proportions of each omics stable, and comparing them to the complete model. And finally, we tested reproducibility by randomly splitting the dataset into two non-overlapping sets of donors, training independent MOFA models on each, and correlating the feature weights between matching factors. Across all analyses, feature weight correlations remained high (Pearson’s *r* > 0.7), indicating that the MOFA factors are stable and reproducible even when the dataset is restricted or split. In addition, to assess the impact of a single *view* on the MOFA factors, we performed another set of experiments by dividing the samples into two groups: one consisting of samples with proteomics data and another consisting of samples lacking proteomics data. For this test, we focused on Factor 14, in which variance was mostly driven by proteomics. We then performed regression analyses on all phenotypes to evaluate how the presence or absence of proteomics measurements influences the results (**Supplementary Figure S8)**. Although both groups exhibited meaningful associations with several covariates, the group with proteomics data showed considerable stronger signals and more biologically interpretable results. This suggests that the missingness of proteomics affects the final factor scores negatively, and that, possibly, the other *views* are insufficient to infer the missing information. Therefore, it seems important to include all omic layers measurements as well as increase their sample sizes, as they contribute significantly to the MOFA factors and can enhance the overall interpretability and robustness of the multi-omics analysis.

### Factor 8 emerges as a key disease multi-omic unifier

To further understand the molecular functions that could impact the disease, we investigated the omics-specific feature weights contributing to each of the AD-related factors. Here we highlight results from Factor 8, but a detailed description of other factors can be found in the **Supplementary Material - Supplementary Notes** and **Supplementary Figures S9-S16**. Factor 8 was positively associated with higher disease risk, displaying the strongest positive association with Alzheimer’s dementia (*adj. P* = 2.01 x 10^-13^) and NIA-Reagan score (*adj. P* = 4.34 x 10^-10^), and the strongest negative association with global cognitive function (*adj. P* = 2.53 x 10^-20^) and cognitive decline (*adj. P* = 2.53 x 10^-20^). Factor 8 was also positively associated with all pathological traits except for Parkinson’s disease (PD) pathology, as well as with *APOE ε4* (*adj. P* = 4.93 x 10^-3^) and TOMM40’523-L (*adj. P* = 4.10 x 10^-2^) haplotypes, and was, among the AD-related factors, the only factor positively associated with cerebral atherosclerosis (*adj. P* = 3.74 x 10^-2^), frailty (*adj. P* = 1.51 x 10^-2^), and gait score (mUPDRS) (*adj. P* = 8.40 x 10^-3^) (**Figure 1d**).

Factor 8 explained 9.17% of the total variance of the input data, with overall similar contribution of multiple *views*, ranging from 0.94% (H3K9ac) to 1.76% (DLPFC mRNA) (**Figure 1c**). Still, many biologically relevant terms were identified within each *view* (**Figure 2a**). Histone acetylation peaks (H3K9ac) were primarily enriched for immune-related processes, including immune response (*adj. P* = 3.86 x 10^-7^) and interferon signaling (*adj. P* = 1.17 x 10^-3^). H3K9ac peaks also showed positive enrichment for gene markers of *Mic.14* (*adj. P* = 3.61 x 10^-7^), a microglial subtype characterized by interferon response ^36^. At the RNA level, AC and PCG transcriptomes shared enrichment for terms related to heat shock factor 1 (*HSF1*) activation and cellular response to heat stress. AC also showed enrichment for protein folding processes, including response to topologically incorrect proteins (*adj. P* = 2.27 x 10^-6^), while PCG was enriched for immune system and transport-related terms, such as immune receptor activity (*adj. P* = 2.52 x 10^-3^) and organic acid sodium symporter activity (*adj. P* = 1.27 x 10^-2^). Proteomics was enriched for cytoskeletal organization (*adj. P* = 9.91 x 10^-8^), mitochondrial matrix (*adj. P* = 2.35 x 10^-14^) and co-expression protein modules, such as Prot-M24 (ubiquitination-related, *adj. P* = 4.99 x 10^-19^) and Prot-M7 (MAPK metabolism-related, *adj. P* = 3.96 x 10^-9^) (**Figure 2a**). Finally, at metabolomics level, Factor 8 was enriched for pathways involving the urea cycle, arginine, and proline metabolism (*adj. P* = 3.10 x 10^-2^) (**Supplementary Table S1**).

**Figure 2.**
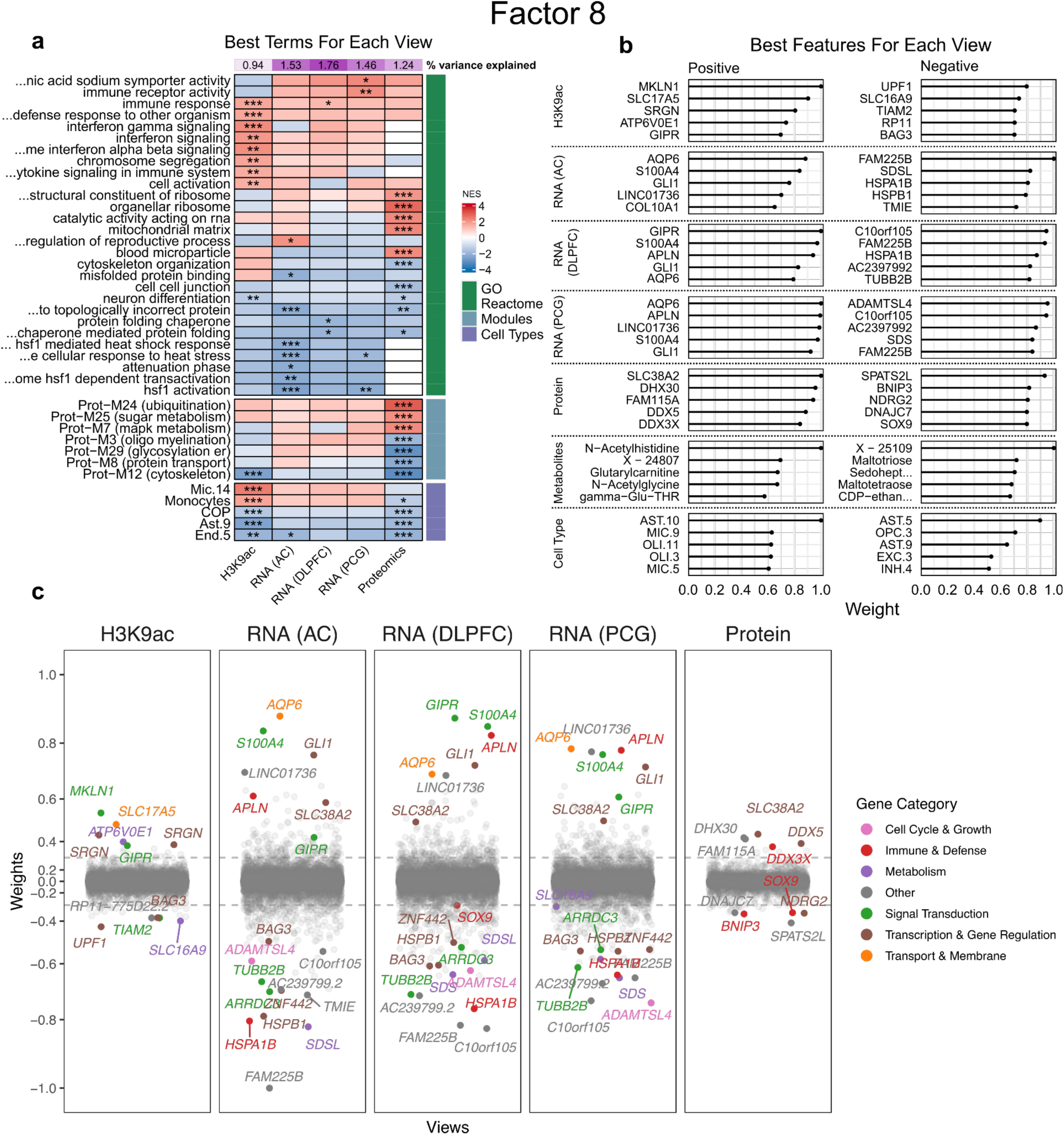
Characterization of factor 8 functional enrichment and feature contributions. **(a)** Gene Set Enrichment Analysis of factor 8, showing the most significant terms for each *view*. *P*-values were corrected using *Benjamini-Hochberg* (BH) method to address multiple comparisons. Terms were selected based on the lowest adjusted *P* values for each *view*. ‘GO’ and ‘Reactome’ comprise terms from Gene Ontology and Reactome databases, respectively. ‘Modules’ include relevant co-expression protein modules from ^46^. ‘Cell Types’ refers to subpopulations of cell lineages as determined by ^47^. The color scale shows the normalized enrichment score (NES) direction and strength from negative (blue) to positive (red). (*) = -log_10_ (adj. *P* < 0.05); (**) = -log_10_ (adj. *P <* 0.01); (***) = -log_10_ (adj. *P <* 0.001). **(b)** Top five positive and negative feature weights in each *view*. Weights are scaled to each *view*. Positive weights indicate that the feature has higher levels in the cells with positive factor values, and vice versa. Metabolites shown as ‘X –’ have not been characterized. **(c)** Jitter plot showing genes with weights in each *view* scaled across all *views* (cell types and metabolomics are not shown). Genes from selected pathways and with an absolute weight greater than 0.3 are highlighted (colored by category).

Further analysis of the top features contributing to Factor 8 (**Figure 2b–c**) highlights the multifaceted biological signature linked to AD pathophysiology. Genes such as *AQP6*, *GIPR*, *GLI1*, *S100A4*, and most members of the SLC family have previously been shown to be upregulated in AD brains ^37^. In addition, also contributing to Factor 8, the stress response cells of AST.10 (*SLC38A2*), previously associated with increased tau pathology and rapid cognitive decline ^38^, contributed significantly to this factor. Metabolically, glutarylcarnitine was associated with mitochondrial fatty acid β-oxidation, which was prominent in our analysis ^39,40^. Overall, the integration across multiple omics captured by Factor 8 reveals key mechanisms such as immune response (epigenetic), reduced cytoskeletal organization (mainly proteomics-driven), decreased levels of heat shock chaperones, and increased transmembrane transporter gene expression, highlighting interconnected pathological mechanisms in Alzheimer’s disease ^41–45^.

### Discordant immune-related functions modulate the protective effects of factors 2 and 3

Factors 2 and 3 exhibited an overlap of several functional terms across multiple *views*, representing a convergence of the same genes and pathways at distinct molecular levels (**Figure 1f**). Both factors were strongly enriched for processes involving immune response and displayed overlapping functional terms across their *views*, with the key difference that these enrichments were mostly in opposite directions (**Supplementary Figures S10a and S11a**). Among the top feature contributors to Factors 2 and 3, we identified several genes related to neuroinflammatory processes and β-amyloid metabolism, such as *SERPINA3* and *CD14* (**Supplementary Figures S10b and S11b**). Additionally, Factor 2’s highest positive weights for cell types included AST.5, OPC.3, and MIC.7, reactive-like cell subpopulations linked to brain aging trajectory independent of AD progression ^48^. In contrast, Factor 3 included the same cell types among the highest negative weights, alongside positive contributions from a microglial subpopulation involved in cell surveillance (MIC.4).

Although these factors shared overall functional enrichments in opposite directions, both were negatively associated with AD and neuropathologies (**Figure 1d**). We then investigated the feature weights contributing to all 7 omics *views*. We found that for most omics there was a significant negative correlation, ranging from Pearson’s r = -0.55 in H3K9ac to -0.12 in AC RNA-Seq, explaining the observed rank-based enrichment results (**Figure 3a**). However, specifically in the DLPFC transcriptomics, we identified 2 clusters of seemingly contradictory gene contributions observed between Factors 2 and 3 (**Figure 3b**). Cluster 1 shows a strong negative association (r = -0.32) and cluster 2 shows a strong positive correlation (r = 0.80). Further functional enrichment analysis of these two clusters revealed a concordant gene signature of increased synaptic function and decreased vasculature-related gene expression (cluster 2), and the previously described discordant immune-related function in the discordant set of genes (cluster 1), suggesting potential protective immune mechanisms operating through different cellular pathways but converging on similar neuronal outcomes (**Figure 3c**). These results observed in Factors 2 and 3 suggest that the timing and context of immune activation may be more critical than the specific immune pathways involved in determining whether neuroinflammatory responses are protective or detrimental in AD pathogenesis. Further studies are needed to clarify the temporal dynamics and cellular specificity of these immune-related processes in AD progression.

**Figure 3.**
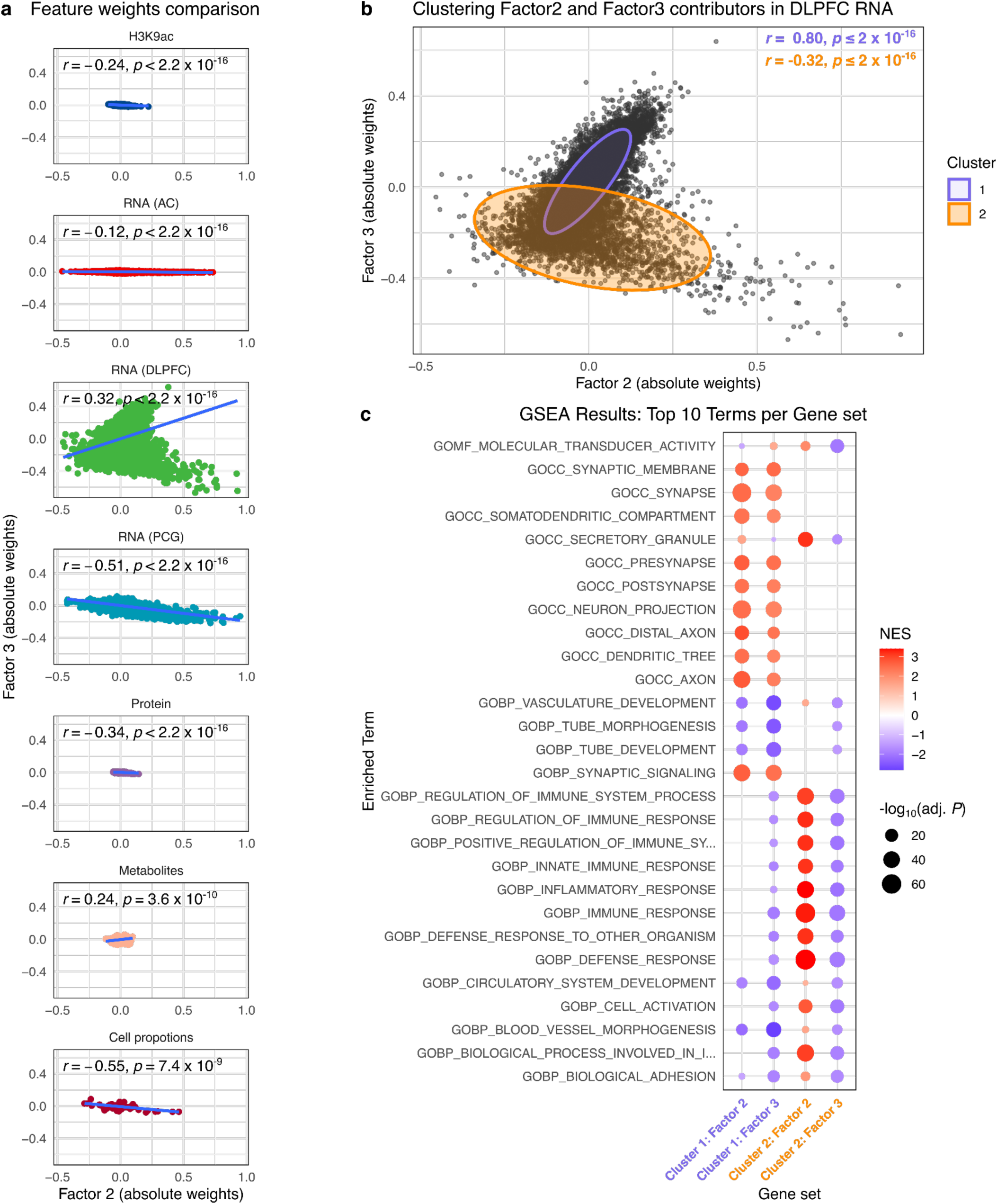
Comparison of features contributing to Factors 2 and 3. **(a)** Feature weights for each *view*, x-axis represents Factor 2, while y-axis refers to Factor 3. The blue line represents the regression line. Pearson R and respective P-values are shown in each *view*. **(b)** Shows the comparison of features from Factor 2 and 3 for the RNA (DLPFC) *view*. A Gaussian Mixture Model (GMM) was applied to cluster the data into 2 clusters: cluster 1 (in blue) and cluster 2 (in red). **(c)** Gene ontology enrichments using GSEA for features in contributing to Factors 2 and 3 in each GMM cluster. The color scale shows the normalized enrichment score (NES) direction and strength from negative (blue) to positive (red). The size of the circles represents -log_10_ (adj. *P*).

### Additional factors reveal complex molecular mechanisms involved in AD

The remaining AD-related factors also provide important aspects of aging and AD biology. Factor 1 (**Supplementary Figure S9**), which explains the largest proportion of total data variation, was negatively enriched in mitochondrial and ribosome-related terms, and positively enriched with MAPK and sugar metabolism, RNA splicing, and ubiquitination terms. Most of these enrichments were primarily driven by transcriptomics (AC) and proteomics, with additional contributions from anterior cingulate transcriptomics driven by heat shock stress response genes. Factor 1 was also positively associated with AD in our tests, aligning with previous findings linking these functions to high AD pathology and worse cognitive outcomes at the proteomics level ^49^. Factor 14 (**Supplementary Figure S14**) demonstrated the second strongest association with AD pathology after Factor 8, uniquely correlating with both PD pathology (*adj. P* = 4.87 x 10^-6^) and major depressive disorder (*adj. P* = 2.78 x 10^-2^). This factor was primarily driven by proteomics (2.83% variance) and metabolomics (1.19% variance), with enrichments pointing to the downregulation of respirasome and oxidoreductase related terms, increased protein co-expression modules linked to Cell-Extracellular Matrix interactions and MAPK metabolism. Factor 14 was also linked to metabolomic alterations in energy metabolism pathways, including glycolysis, gluconeogenesis, and pyruvate metabolism, having uridine diphosphate (UDP)-glucose and FG dipeptide (F and G amino acids) as top metabolite markers. Factor 42 (**Supplementary Figure S16**), despite explaining only 1.7% of total variance, captured some aspects of vascular biology relevant to AD pathogenesis. Top contributors to Factor 42 were PCG and AC transcriptomes, showing significant enrichments for marker genes of subpopulations of pericytes (Peri.2), smooth muscle cells (SMC.2), and arteriole, along with *APOLD1* as top positive weights in PCG, a gene involved in endothelial cell function and hemostasis.

Our analysis also identified significant protective signatures against AD pathology. Factor 26 (**Supplementary Figure S15**) exhibited the strongest protective effect, with enrichment for RNA processing mechanisms, vascular-related pathways, and cytoskeletal organization that may preserve cognitive function despite pathological burden. Factor 4 (**Supplementary Figure S12**) emerged as the second strongest protective factor, negatively associated with AD dementia (*adj. P* = 2.34 x 10^-6^) and NIA-Reagan score (*adj. P* = 5.78 x 10^-4^), while positively correlating with global cognitive function (*adj. P* = 1.74 x 10^-7^). Notably, Factor 4 was uniquely negatively associated with Lewy body disease (*adj. P* = 1.71 x 10^-2^), PD pathology (*adj. P* = 8.27 x 10^-5^), and depressive symptoms (*adj. P* = 9.09 x 10^-3^). Its protective mechanisms involve high expression of genes like *VGF* and *CX3CR1*, enrichment for microglial subtype MIC.14 characterized by interferon response, and metabolomic signatures involving lysophospholipids and long-chain polyunsaturated fatty acids. The features with the highest positive weights included several genes considered protective against AD, such as *CX3CR1*, which modulates microglial activation, and metabolites like trigonelline, which has shown protective functions in mice by ameliorating axonal and dendritic atrophy in cortical neurons treated with β-amyloid ^50^. Finally, Factor 12 (**Supplementary Figure S13**) showed protective effects through increased cell activation, cell signaling, and extracellular matrix organization, strongly driven by H3K9ac epigenetic markers. These markers are also enriched for RNA co-expression modules M7, involved in transcription regulation and tau tangles, and M114 involved with immunity and associated with β-amyloid.

### Identification of multi-omic molecular subtypes of AD

In order to represent the molecular heterogeneity of the aging brain and its relation with Alzheimer’s dementia, we applied the SpeakEasy ^51^ consensus clustering algorithm using all 50 MOFA latent factors to group correlated participants into distinct clusters (**Figure 4a**). These clusters are expected to provide an unbiased classification of participants based on molecular signatures from multi-omics in the postmortem brain, which is important because patient stratification may enable more precise understanding of disease mechanisms and tailored therapeutic approaches. We found three major clusters broadly differentiating individuals from a non disease state, with lower Alzheimer’s Disease and Related Dementias (ADRD) pathologies (major cluster 1) to individuals in a disease-like state with elevated pathology and lower cognition scores (major cluster 3) (**Figure 4b**). Further refinement of each major level cluster resulted in 11 subclusters with a more detailed stratification of protective and pathological subtypes of AD.

**Figure 4.**
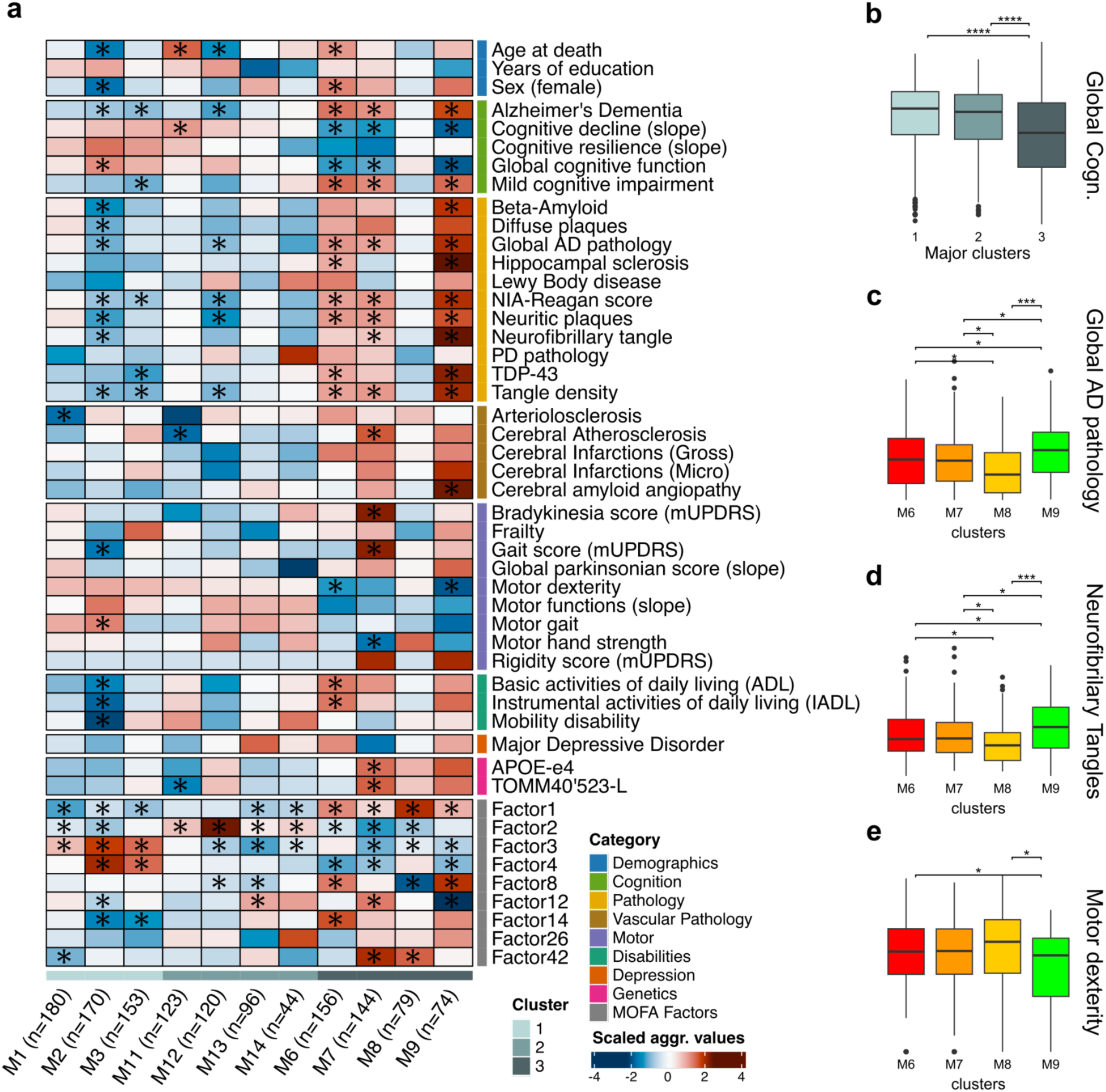
Molecular subtyping of ROS/MAP participants. **(a)** Association of clusters with Alzheimer’s covariates and factors. Color scale shows the scaled aggregated values strength and direction from negative (blue) to positive (red). ‘Cluster’ represents the first level of clustering, with 3 distinguished clusters. The X-axis shows clusters from the second level of clustering, along with the number of samples for each cluster in parenthesis. The Y-axis shows factors and covariates subdivided in nine categories. The traits with asterisks diverge significantly from the mean or median (for traits with gaussian distribution) of all clusters after 1000 permutations. **(b)** t-test comparing global cognitive function across the three groups at major cluster level. **(c-e)** t-test comparing clusters M6, M7, M8 and M9 with the traits: global pathology AD burden **(c)**, neurofibrillary tangle burden **(d)** and motor dexterity **(e)**. (*) = adj. *P* < 0.05; (**) = adj. *P* < 0.01; (***) = adj. *P* < 0.001; (****) = adj. *P* < 0.0001.

Among the clusters related to a non/low disease individuals (major cluster 1), three subclusters were identified (M1, M2, and M3). Participants in cluster M1 displayed phenotypic traits that were not significantly different from the average. Participants in cluster M2 presented a negative correlation with Alzheimer’s clinical diagnosis and NIA-Reagan scores, as well as with most pathological traits and all disabilities; however, it was also associated with a younger age of death, possibly confounding observation in the other traits. Cluster M3, on the other hand, exhibited a negative correlation with Alzheimer’s clinical diagnosis and NIA-Reagan scores, and did not show a significant correlation with age of death (**Figure 4b**). Clusters M2 and M3 also presented higher than average levels of Factors 3 and 4, two factors negatively associated with Alzheimer’s clinical diagnosis and most of AD-related traits (**Figure 1d**).

Among clusters positively associated with AD (major cluster 3), four subclusters were identified (M6, M7, M8, and M9). While participants in the cluster M8 did not show differences compared to the average, clusters M6, M7 and M9 shared similar clinical and pathological characteristics. These clusters were positively associated with Alzheimer’s dementia, mild cognitive impairment, and NIA-Reagan scores, as well as with most pathological traits, and negatively correlated with cognitive function and cognitive decline (**Figure 4c-e**). Despite these similarities, each cluster showed distinct phenotypic differences. Participants in M6, for example, had higher numbers of females and higher age of deaths compared to the average. It was also the only cluster with higher than average indices of instrumental activities of daily living (IADL). Cluster M7 had the strongest associations with motor traits, namely Bradykinesia score (mUPDRS), gait score (mUPDRS), and hand strength, compared to any other cluster, but also had higher frequencies of AD genetic risk factors (APOE/TOMM40). Meanwhile, participants from cluster M9 showed the strongest correlations with neurofibrillary tangles, hippocampal sclerosis, and gross cerebral infarctions compared to any other cluster. These clusters also differed in their molecular signatures. While cluster M7 was mostly influenced by Factor 8, cluster M9 was primarily associated with Factor 42 and, to a lesser extent, Factor 12. Conversely, M6 correlated with Factor 14 and to a smaller degree with Factors 1 and 8. Overall, multi-omic stratification reveals discrete AD subtypes with unique molecular and phenotypic profiles, which may help refine prognosis and guide the development of tailored therapeutic approaches.

## DISCUSSION

Here, we employed MOFA, an unsupervised method for identifying the principal sources of variation across multimodal data, to analyze multi-omics data from the ROS/MAP cohorts. The data included H3K9ac histone acetylation, mRNA expression from three brain regions (AC, PCG and DLPFC), proteomics, metabolomics and proportions of cell-type specific subpopulations. MOFA inferred a set of 50 factors that captured biological and technical sources of variability across multiple omics, with 9 of these factors showing a meaningful association with AD. Then, we performed GSEA on the selected factors to determine their molecular profiles and identified genes of interest. Finally, we clustered the ROS/MAP samples and identified several groups of people associated with varying AD traits and factors.

We identified multiple factors linked to pathophysiological mechanisms associated with AD. Several factors captured distinct immune system processes, including interleukin signaling, inflammatory responses, and cytokine production, all of which are known to be dysregulated in AD ^52–54^. In addition, MOFA detected other key mechanisms such as cytoskeleton organization, proteostasis, and synaptic activity, each of which has been previously implicated in AD pathology ^55–57^. Moreover, some factors showed strong associations with comorbid conditions commonly linked to AD, including depression and vascular pathologies such as arteriosclerosis and cerebral amyloid angiopathy, both of which are known to be risk factors to AD ^58,59^. Finally, our clustering approach was able to classify participants into distinct molecular subtypes reflecting the recognized molecular heterogeneity of AD ^60^. One key feature that distinguishes this analysis from prior work is the description of how these AD-implicated systems span multiple omics. This approach enables disease investigations to proceed with greater confidence, focusing on coordinated signals supported across data types, which have independent sources of biological and technical variation.

One of these multi-omic signals - Factor 8 - emerged as the signal most strongly associated with Alzheimer’s dementia and was linked to nearly 80% of the tested phenotypes. Top features included genes related to AD pathophysiology and cell subpopulations positively associated with AD. These findings may serve as a resource for candidate genes of interest for potential novel therapies. Importantly, the positively weighted gene *S100A4* has been implicated in multiple AD-related pathways, including neuroinflammation and oxidative stress. *In vitro*, *S100A4* knockdown in amyloid-β oligomer (AβO)-treated BV2 microglia improved cell viability, reduced lactate dehydrogenase release, and partially alleviated apoptosis. *S100A4* inhibition attenuated pro-inflammatory responses and promoted an anti-inflammatory state. Furthermore, *S100A4* knockdown mitigated oxidative stress, restoring mitochondrial function and decreasing reactive oxygen species levels ^61^. The *BAG3* (*BCL-2-associated athanogene 3*) gene exhibited high negative weights across multiple *views*, suggesting its implication in several pathways linked to factor 8, including immune response, Prot-M12 (related to the cytoskeleton), and protein folding. *BAG3* is a multi-functional protein involved in many biological processes, with evidence showing a protective role against AD by facilitating phospho-tau degradation ^62^ and autophagy of diseased mitochondria ^63,64^. *BAG3* also regulates mitochondrial function by supporting calcium (Ca^2+^) transport in and out of mitochondria ^65,66^, drives the cytoskeleton dynamics by regulating actin folding ^67–69^, inhibits caspase activation ^70,71^ and interacts with *HSF1* ^72^ and MAPK molecules (e.g. *MAP3K5*) ^73,74^. We hypothesize that factor 8 captures genes acting in the regulation of immune response at the level of H3K9ac acetylation; protein folding and *HSF1* activation in RNA; and cytoskeleton dynamics, MAPK metabolism, and mitochondrial matrix in proteomics, reflecting a coordinated multi-omic program that together may contribute to AD pathogenesis.

Our analysis revealed an intriguing correlation in Factors 2 and 3. While both demonstrated protective effects against AD pathology, their overall enrichments pointed to discordant immune signatures. However, in DLPFC RNA, we found concordant upregulation of genes related to synapse and neuronal-related terms, along with another set of genes pointing to opposite immune cell activation patterns. This observation initially challenges the conventional view of neuroinflammation in AD as uniformly detrimental. However, our finding suggests that the timing, context, and specific nature of immune activation may determine whether neuroinflammatory responses are protective or harmful. The positive weights for reactive-like glial subpopulations in Factor 2 and surveillance microglial cells in Factor 3 indicate that different glial activation states may confer protection through distinct but complementary mechanisms, potentially by promoting neuronal resilience or facilitating clearance of pathological proteins. These observations are, however, based on postmortem samples, and further studies in living human brains will be important for comparing these dynamics across disease stages.

Beyond these primary factors, our analysis identified additional significant contributors to AD heterogeneity. Factor 1, explaining the highest proportion of total variance, captured fundamental biological processes including mitochondrial function, ribosomal activity, and proteostasis pathways, suggesting that these core cellular mechanisms constitute a major axis of variation in the aging brain. Factor 14 showed the second strongest association with AD pathology and was also associated with PD pathology and major depressive disorder, highlighting potential shared mechanisms between these disorders. Factors 26 and 4 exhibited protective signatures, with Factor 26 showing the strongest protective effect through enrichment for RNA processing mechanisms and vascular-related pathways, while Factor 4 was uniquely negatively associated with Lewy body disease and PD pathology, suggesting broader neuroprotective mechanisms. Factor 4 also exhibited high positive weights for genes and metabolites known to be protective against AD. *FKBP5* and *VGF* are multifunctional genes with high absolute weights across multiple *views* and direct relevance to AD. The immunophilin FK-506-binding protein 51 (*FKBP51*) may contribute to AD development through its interaction with the heat shock protein *Hsp90*, which leads to tau phosphorylation, inhibits tau degradation, and results in neurotoxic tau accumulation ^75–77^. Conversely, the neurotrophin *VGF* (a nerve growth factor-inducible protein) and its derived peptides play a protective role in AD. For instance, the biopeptide *TLQP-62* positively influences synaptic plasticity and memory ^78^. Furthermore, increasing *VGF* expression in the 5xFAD mouse model has been shown to reduce β-amyloid levels in the brain and improve memory performance ^79,80^.

While our study provides valuable insights, it is important to acknowledge its limitations. MOFA, being a linear model, may overlook non-linear relationships between features within and across assays ^81,82^. Additionally, while our multi-omics approach provides unprecedented molecular resolution, it cannot fully capture the complex cellular interactions and spatial organization of the brain. There was a considerable disparity in the amount of information available for each omic type (e.g., 17,309 mRNA transcripts versus 667 metabolites), which may limit the ability to assess the relative importance of individual features. Moreover, our epigenetic data was restricted to H3K9ac acetylation data, constraining our ability to capture a more comprehensive view of epigenetic regulation, potentially overlooking key methylation-associated mechanisms involved in the molecular framework of AD. In addition, ROS/MAP cohorts are mainly composed of individuals of European ancestry, and additional data from other human populations are needed to generalize the results. The cross-sectional nature of post-mortem tissue analysis limits our ability to establish causal relationships and temporal dynamics. Finally, this study employs an exploratory approach, utilizing collected omics data to generate data-driven hypotheses. These hypotheses will require further investigation through complementary, targeted experimental approaches conducted with statistical rigor.

In summary, our integrative multi-omics analysis provides a comprehensive map of molecular mechanisms underlying AD heterogeneity, revealing both shared pathological processes and distinct molecular subtypes. Our findings have several important implications for AD research and therapeutic development. First, they highlight the capability of identifying well known signatures from AD using unbiased dimensionality reduction of multi-omics data. Second, these findings also expand beyond classical amyloid/tau pathology and can be linked to a series of aging related phenotypes. Third, by integrating multi-omics in a common framework we were able to connect apparently disjoint disease components usually observed in single-omics. Finally, by identifying AD associated factors and molecular subtypes we create a basis for patient stratification in clinical trials, potentially improving the success rate of AD drug development. Progress in the next generation of AD therapy will likely stem from “combination” approaches that reinforce proteostasis, buffer neurotransmission, and reduce chronic inflammation, tailored to patient-specific genetic and epigenetic profiles. The genes and features highlighted herein exemplify both the challenge and opportunity in pursuing this multi-factorial, precision medicine vision for AD.

## METHODS

### Study Participants

Participants came from two ongoing longitudinal clinical-pathological cohort studies of aging and Alzheimer’s disease conducted by the Rush Alzheimer’s Disease Center (RADC), the Religious Orders Study (ROS) and the Rush Memory and Aging Project (MAP). ROS was established in 1994 and includes nuns and priests from across the United States ^35,83,84^, while MAP began in 1997 and includes individuals from retirement communities, subsidized housing, and private dwellings in Illinois ^35,84,85^. Both cohorts, collectively referred to as ROS/MAP, admit older individuals without known dementia at the time of study enrollment. Participants agree to annual clinical assessments, cognitive tests, and blood collections, as well as organ donation (spinal cord, muscles, nerves and brain). Through March 2025, ROS/MAP enrolled 4,025 participants, of whom 2,127 came to autopsy. The present study is focused on 1,383 deceased individuals (**Table 1**) with respective omics data available as described below.

### Clinical and neuropathological evaluations

Here we briefly describe the harmonized, deeply phenotyped measures performed on ROS/MAP participants. These include clinical diagnoses of Alzheimer’s dementia and MCI; measures of global cognitive function, cognitive decline, and resilience; neuropathologic evaluations of AD pathology including NIA-Reagan classification, global AD pathology, β-amyloid, tangles, diffuse and neuritic plaques, TDP-43, Lewy bodies, PD pathology, and hippocampal sclerosis; vascular pathologies including arteriolosclerosis, atherosclerosis, CAA, and gross and microinfarcts; frailty classification; indices of global motor and parkinsonian function; disability measures including ADL, IADL, and mobility disability; and diagnosis of depressive symptoms and major depressive disorder (MDD). More detailed descriptions are provided in the **Supplementary Material – Supplementary Notes**.

### Cognitive function and Alzheimer’s dementia

Cognitive function was assessed annually for up to 23 years (Mean = 7.4, SD = 4.9) using a battery of 21 performance tests. Scores from 17 tests were standardized into z-scores using the baseline mean and standard deviation and then averaged to derive a composite measure of global cognition. Lower z-scores reflect lower performance relative to baseline cohort norms ^86^. For each individual, the annual rate of change in global cognitive function (cognitive decline) was estimated using a linear mixed effects model, controlling for age at baseline, sex, and years of education ^87^. To examine cognitive resilience, a similar model was used that further adjusted for neuropathologic measures including global AD pathology, β-amyloid, neurofibrillary tangles, Lewy body disease, TDP-43 pathology, infarcts, hippocampal sclerosis, and vascular pathologies ^88^. Clinical diagnosis of cognitive status was made using a modified three-stage procedure based on a neuropsychologist’s review of 19 test scores, followed by clinician review and diagnosis ^89–91^. Final classification analyzed Alzheimer’s dementia vs. no dementia (MCI + NCI) and with cognitive impairment (AD + MCI) vs. no cognitive impairment (NCI) ^92^.

### Neuropathologic outcomes

Neuropathologic evaluations were performed blinded to clinical data and included uniform assessments of Alzheimer’s disease and common co-pathologies. A modified version of the NIA-Reagan criteria was used to classify the likelihood of Alzheimer’s disease pathology on a 4-point scale from no to high likelihood based on the presence of both neurofibrillary tangles (Braak) and neuritic plaques (CERAD) ^93^. Quantitative measures of pathology were derived from silver-stained slides across multiple brain regions. Summary scores were computed for neuritic plaques, diffuse plaques, and neurofibrillary tangles ^94^. These were standardized within each region, averaged across regions, and square root-transformed to reduce skewness. A composite global AD pathology score was then calculated by averaging these three summary scores ^95,96^. Tangle density was assessed as the mean square root of tangle scores in eight regions, including the hippocampus and frontal cortex ^97^. β-amyloid burden was quantified using immunohistochemistry across eight regions, and summarized as the square root of the mean regional β-amyloid score ^98^. Additional pathologies assessed included TDP-43 staging (no TDP-43 pathology or in amygdala only, and TDP-43 pathology extending beyond amygdala) ^99^, Lewy body disease (binary classification based on presence in seven regions) ^100^, and hippocampal sclerosis (graded 0–5, with severe pathology defined as a score of 5 in CA1 or subiculum) ^101^. In addition, a binary variable representing Parkinson’s disease (PD) pathology was defined based on the presence of Lewy bodies and the severity of neuronal loss in the substantia nigra.

### Vascular pathologies

Cerebral infarctions were documented separately as gross and microinfarcts based on standardized neuropathologic examination ^102,103^. Arteriolosclerosis and cerebral atherosclerosis were graded semi-quantitatively (0–3) based on severity observed in histologic and gross pathology assessments, respectively ^104,105^. Cerebral amyloid angiopathy (CAA) was quantified in four cortical regions by scoring meningeal and parenchymal vessel involvement from 0 to 4, and averaging across regions. CAA severity was also categorized semi-quantitatively using established cutpoints ^106,107^.

### Motor and parkinsonian function

Motor function was evaluated annually using ten performance tests encompassing dexterity, gait, strength, and balance, and a derived slope representing motor decline was estimated using a linear mixed effects model, controlling for age at baseline, sex, and years of education. ^108^. Performance scores were standardized based on cohort baseline means and combined to generate composite measures of global motor function, gait, dexterity, and hand strength ^109^. Parkinsonian signs (bradykinesia, rigidity, tremor, gait disturbance) were assessed using modified motor items from the Unified Parkinson’s Disease Rating Scale. Scores were scaled from 0 to 100, with higher scores reflecting more severe impairment ^110^. Longitudinal trajectories of global parkinsonian score were derived from the average of the four domain scores modeled over time, adjusted for baseline age, sex, and years of education ^111,112^.

### Disability and frailty

Disability was assessed using three measures: basic activities of daily living (ADL), instrumental activities of daily living (IADL), and mobility disability. ADL and IADL scores reflected the number of activities with which participants reported needing assistance ^113,114^, while mobility disability was assessed with the Rosow-Breslau scale ^115^. Frailty measure reflects multisystem vulnerability, based on four components: grip strength, timed walk, body composition (BMI), and fatigue. The measure was constructed by converting each raw component score into a z-score using baseline mean and standard deviation values from all participants^116,117^.

### Psychological factors

Depressive symptoms were measured annually using a 10-item modified Center for Epidemiologic Studies Depression Scale (CES-D). Each participant’s mean score across visits was used to reflect their enduring disposition toward depressive symptoms ^118–120^. A clinical diagnosis of major depressive disorder was rendered by a physician based on structured interviews and DSM-III-R criteria. A dichotomous indicator of probable depression was derived by collapsing the diagnostic scale into highly probable vs. other scores ^121^.

### Genetics

Genotyping of DNA extracted from either brain tissue or peripheral blood mononuclear cells was conducted using high-throughput sequencing, targeting codon 112 (position 3937) and codon 158 (position 4075) of exon 4 of the APOE gene ^122^. A dichotomized score was created where samples were categorized into those with the APOEε4 allele and those without. TOMM40’523 genotypes are determined by rs10524523 (chr19:44,899,792-44,899,826, human genome reference assembly GRCh38/hg38), a homopolymer length polymorphism (poly-T), at intron 6 of the TOMM40 gene ^123,124^. Allele lengths are classified according to the number of poly-T repeats: short alleles (S) have fewer than 20 repeats, long alleles (L) have 20 to 29 repeats, and very long alleles (VL) have 30 or more repeats ^125^. A dichotomized score was created where samples were categorized into those with a least one long allele and those without.

### Omics data

#### H3K9ac histone acetylation (ChIP-seq)

H3K9ac histone acetylation data generation has been previously described ^126^. In essence, gray matter tissue from ROS/MAP subjects was dissected from the Dorsolateral Prefrontal Cortex (DLPFC), homogenized, incubated with H3K9ac antibody using the ChIP Dilution Buffer and purified with protein A Sepharose beads. The resulting DNA was used for Illumina library construction and then sequenced on the Illumina HiSeq (36 bp single-end reads). Low-quality samples were removed and H3K9ac domains were defined by calculating all genomic regions that were detected as a peak in at least 15% of the samples. Regions neighboring within 100 bp were merged, and very small regions of less than 100 bp were removed, resulting in 26,384 H3K9ac domains from 632 samples. The Data was normalized, log transformed and technical confounders removed.

#### Transcriptomics (RNA-Seq)

RNA-Seq data were previously generated and raw data can be downloaded from the AMP-AD platform (syn3388564). Briefly, RNA RNA were extracted from gray matter tissues from 3 regions: anterior caudate (AC), posterior cingulate gyrus (PCG) and DLPFC. Extraction was performed using the Qiagen miRNeasy kit and DNase treatment, with quality assessed by Bioanalyzer and quantity by Nanodrop. Three batches of RNA-Seq libraries were prepared from different brain regions using distinct protocols: Batch 1 (DLPFC) used the dUTP method with poly-A selection; Batch 2 (DLPFC, PCC, AC) used KAPA’s stranded kit with ribosomal depletion; Batch 3 (DLPFC, PCC) involved Chemagic extraction, RiboGold depletion, and a modified TruSeq protocol. Sequencing was performed using Illumina platforms: Batch 1 used HiSeq (101bp PE reads, 50–150M reads/sample), Batch 2 used NovaSeq6000 (2×100bp, 30M reads/sample), and Batch 3 used NovaSeq6000 (2×150bp, 40–50M reads/sample). A harmonized data processing was applied to all three regions as previously described ^127^. Briefly, after QC, quantification was performed using Kallisto (v0.46). Gene level counts were filtered to keep only genes with at least 10 counts in >50% of samples. Counts were adjusted by GC content and gene length using CQN (conditional quantile normalization) and converted to log2-CPM (counts per million) followed by quantile normalization using the limma voom function. Finally, a linear model was applied to remove major technical confounding factors matrics as reported by Picard and Kallisto. The final resulting matrices included 17,294 mRNAs for 721 AC, 656 PCG and 1,210 DLPFC samples.

#### Single-nuclei RNA sequencing (snRNA-Seq)

The protocol for the isolation of nuclei from frozen postmortem brain tissue, RNA sequencing and data processing was previously published ^128,129^. Briefly, 50 to 100mg of DLPFC gray matter from ROS/MAP subjects were homogenized, the cells nuclei filtered and sequenced by 10X Single Cell RNA-Seq Platform allowing the creation of 128 pooled libraries. Preprocessing of each library involved alignment, normalization, cell clustering, background noise removal, de-multiplexing, classification of nuclei for cell types, removal of low-quality nuclei and detection of doublets for removal. Following major cell-type classification, a sub-clustering analysis was performed resulting in 96 cell subpopulations measured for 424 donors ^129,130^. We considered the relative fractions of each subpopulation by donor in our downstream analysis.

#### TMT-MS-based quantitative proteomics

Methods employed data generations and processing has also been detailed in a prior publication ^131^. Briefly, DLPFC tissue samples were homogenized, digested and purified. Then the resulting peptides were labeled using the TMT 10-plex kit and fractionated by high pH. All fractions were resuspended and analyzed by liquid chromatography coupled to tandem Mass Spectrometry and the resulting spectra were searched against the UniProtKB human proteome database. After removing technical variance 8,425 proteins from 596 donors were used in our downstream analysis.

#### Metabolomics profiling

Metabolomics data generation and processing has been previously described ^132^. Summarily, tissues from DLPFC subjects were homogenized, the proteins were removed and the resulting extract was divided into four fractions: two for ultra-high performance liquid chromatography-tandem mass spectrometry (UPLC-MS/MS) with positive ionization, one for UPLC-MS/MS with negative ionization and one for UPLC-MS/MS polar platform with negative ionization. Peaks were identified, and quantified and technical confounders were excluded. The process resulted in 667 metabolites from 500 subjects that were used in our analysis.

### Multi-omics Factor analysis (MOFA)

MOFA is a computational framework for unsupervised multi-omics integration analysis capable of capturing both biological and technical sources of variability across multiple omics datasets. Intuitively, it can be viewed as a generalization of principal component analysis to multi-omics data. The method employs Bayesian inference and probabilistic models to derive a low-dimensional representation of multiple data modalities in terms of latent factors that capture major sources of variation within the data. It also provides the loadings of each individual feature for every factor, granting a mapping between the high and low-dimensional spaces. Furthermore, MOFA disentangles to what extent each factor is unique to a single data modality or is manifested in multiple modalities, thereby revealing shared axes of variation between the different omics layers. The model is able to efficiently handle missing values and is robust to scalability of large volumes of data ^133^.

### MOFA implementation

The analysis was conducted utilizing the ‘MOFA2’ package ^134^, version 1.10.0, in R. Initially, to conform the data to one of MOFA’s input data formats, we transformed each omic dataset into matrices, with the same number of samples (missing samples were added as NAs) arranged in the same sequence. To perform model training, we used MOFA’s default data, model and training settings, with the exception of two adjustments: the number of factors was set to 50 (default is 15) and the data was scaled to have the same unit variance (default is FALSE).

### Factor selection

In order to retain only the factors associated with AD, we conducted a regression analysis using the 50 factors generated by MOFA and the 38 aforementioned covariates, the tests were controlled for sex, age at death and years of education. For every covariate we employed different regression techniques tailored to each data type: for continuous variables we used the linear regression method implemented in R with the lm function from the ‘Base’ package, version 4.3.1 ^135^. For binomial covariates, we employed the Bayesian logistic regression approach using the bayesglm function from the ‘arm’ package, version 1.14.4 in R ^136^. For ordinal covariates, we utilized the cumulative link model implemented in R with the clm function from the ‘ordinal’ package, version 2023.12.4 ^137^. We selected the factors that exhibit statistically significant association (p < 0.05) with either NIA-Reagan score or with clinical cognition status. Nine factors were elected for downstream analysis: factors 1, 2, 3, 4, 8, 12, 14, 26, 48.

### Gene Set Enrichment Analysis (GSEA)

For GSEA, we used the ‘fgsea’ package version 1.26.0 ^138^ in R. GSEA was performed for each factor by running the ‘fgseaMultilevel’ function with default settings across all omics. Gene sets for Reactome ^139^ and Gene Ontology ^140,141^ were sourced through the ‘msigdbr’ package version 7.5.1 ^142^. For proteomics, gene background was restricted to all proteins measured in the TMT experiment. The metabolomics enrichment set was adapted from Batra et al. (2022), and the cell subpopulation type set was obtained from Green et al. (2024). Additionally, we incorporated transcriptomic co-expression modules (Mostafavi et al., 2018), proteomic co-expression modules (Johnson et al., 2022), an AD genome-wide association study gene set ^143^, the plaque-induced genes (PIGs) co-expression network ^144^, and two AD-associated microglia subpopulations: DAM ^145^ and HAM ^146^. We also used the function ‘reduceSimMatrix’ (threshold varying from 0.9 to 0.7) from the ‘rrvgo’ package version 1.12.2 ^147^ to reduce terms redundancy.

### Robustness and reproducibility

To evaluate the robustness and stability of the MOFA model, we conducted a series of computational experiments on subsets of the original data. We systematically varied the number of samples and the completeness of the data to assess their impact on the inferred factors. The data was split into training and testing sets using a stratified sampling approach based on the number of omics available for each sample. We generated datasets with 90%, 80%, 70%, 60%, and 50% of the original samples. Additionally, we created non-overlapping 50/50 splits of the data to assess the reproducibility of the factors in independent datasets. For each data subset, we trained a MOFA model with 10 latent factors with a fast convergence mode. To compare the factors inferred from different MOFA models, we matched factors between two models based on the Pearson correlation of their feature weights. The optimal matching of factors was determined using a greedy algorithm. For each pair of matched factors, we calculated the correlation of their weights and corrected for potential sign flips. The mean correlation across all matched factors was used as a measure of the overall similarity between the two MOFA models.

### Cluster analysis with Speakeasy

To categorize the ROS/MAP participants into AD subtypes, we used the 50 factors generated by MOFA as input for the Speakeasy algorithm ^51^. This method is well suited for the analysis, as it has been previously tested with the Lancichinetti-Fortunato-Radicchi (LFR) benchmarks, a widely used test for evaluating overlapping and non-overlapping clustering methods across different types of data ^148^. Speakeasy has been used before with RNA-Seq data ^149^ and proteomics data ^150^ and has been extensively compared against other clustering methods, such as normalized mutual information and weighted gene co-expression networks. Speakeasy does not require extensive parameterization; however, we used the default settings of 100 permutations, Pearson correlation, and a minimum of 30 individuals per cluster. Finally, we analyzed the results at three levels of clustering, with larger clusters at level 1 and more refined clusters containing fewer individuals at levels 2 and 3.

## DECLARATIONS

### Ethics approval and consent to participate

The ROS and MAP studies were approved by an Institutional Review Board (IRB) of Rush University Medical Center, Chicago, IL. All participants agreed to an annual clinical evaluation and signed an informed consent and an Anatomic Gift Act agreeing to post-mortem brain donation. All procedures performed in studies involving human participants were in accordance with the ethical standards of the Institutional Review Board of Rush University Medical Center and with the 1964 Helsinki Declaration and its later amendments or comparable ethical standards. Each participant signed an informed consent form to participate in the study.

### Consent for publication

All authors received and approved the manuscript for publication.

### Availability of data and materials

Raw data used in this manuscript are available through the AD Knowledge Portal (https://adknowledgeportal.org) with the following accession numbers: Epigenetics ChIP-Seq (H3K9ac) (syn4896408), RNA-Seq (DLPFC, AC, PCG) (syn3388564), Proteomics TMT (syn17015098), Metabolomics (syn26007830), and snRNA-Seq (syn31512863). The phenotype data can be requested at the RADC Resource Sharing Hub at www.radc.rush.edu. The relevant code generated by this study is available on the GitHub repository: https://github.com/RushAlz/ROSMAP_MOFA_public_release.

## Acknowledgements

We thank the participants of ROS/MAP cohorts for their essential contributions and gift to these projects, as well the investigators and staff at the Rush Alzheimer’s Disease Center. The data available in the AD Knowledge Portal would not be possible without the participation of research volunteers and the contribution of data by collaborating researchers. The results published here are in whole or in part based on data obtained from the AD Knowledge Portal (https://adknowledgeportal.org).

## Contributions

RAV designed the project. LPS performed the main analysis. KDP performed the clustering analysis. LPS and RAV drafted the manuscript. KDP, ST, CG, ASB, RTR and. DAB edited the manuscript and assisted in writing. PLD and VM generated the snRNA-Seq atlas. YW sequenced the bulk RNA-Seq data. CG, JAS, PLD, and DAB funded the project. All authors read and approved the final manuscript.

## REFERENCES

1. Selkoe, D. J. & Hardy, J. The amyloid hypothesis of Alzheimer’s disease at 25 years. EMBO Mol. Med. 8, 595–608 (2016).

2. Zhang, J. et al. Recent advances in Alzheimer’s disease: Mechanisms, clinical trials and new drug development strategies. Signal Transduct. Target. Ther. 9, 211 (2024).

3. Du, X., Wang, X. & Geng, M. Alzheimer’s disease hypothesis and related therapies. Translational Neurodegeneration 7, 1–7 (2018).

4. Murdock, M. H. & Tsai, L.-H. Insights into Alzheimer’s disease from single-cell genomic approaches. Nat Neurosci 26, 181–195 (2023).

5. Wong, M. W. et al. Dysregulation of lipids in Alzheimer’s disease and their role as potential biomarkers. Alzheimers Dement 13, 810–827 (2017).

6. Leng, F. & Edison, P. Neuroinflammation and microglial activation in Alzheimer disease: where do we go from here? Nat Rev Neurol 17, 157–172 (2021).

7. Yan, X., Hu, Y., Wang, B., Wang, S. & Zhang, X. Metabolic Dysregulation Contributes to the Progression of Alzheimer’s Disease. Front Neurosci 14, 530219 (2020).

8. Clark, C., Dayon, L., Masoodi, M., Bowman, G. L. & Popp, J. An integrative multi-omics approach reveals new central nervous system pathway alterations in Alzheimer’s disease. Alzheimers. Res. Ther. 13, 71 (2021).

9. St John-Williams, L., et al. Targeted metabolomics and medication classification data from participants in the ADNI1 cohort. Sci Data 4, 170140 (2017).

10. Wang, M. et al. The Mount Sinai cohort of large-scale genomic, transcriptomic and proteomic data in Alzheimer’s disease. Scientific Data 5, 1–16 (2018).

11. De Jager, P. L. et al. A multi-omic atlas of the human frontal cortex for aging and Alzheimer’s disease research. Sci Data 5, 180142 (2018).

12. Greenwood, A. K. et al. The AD Knowledge Portal: A Repository for Multi-Omic Data on Alzheimer’s Disease and Aging. Current Protocols in Human Genetics 108, e105 (2020).

13. De Jager, P. L. et al. Alzheimer’s disease: early alterations in brain DNA methylation at ANK1, BIN1, RHBDF2 and other loci. Nat Neurosci 17, 1156–1163 (2014).

14. Allen, M. et al. Gene expression, methylation and neuropathology correlations at progressive supranuclear palsy risk loci. Acta Neuropathol. 132, 197–211 (2016).

15. Carrasquillo, M. M. et al. A candidate regulatory variant at the TREM gene cluster associates with decreased Alzheimer’s disease risk and increased TREML1 and TREM2 brain gene expression. Alzheimers Dement 13, 663– 673 (2017).

16. Allen, M. et al. Conserved brain myelination networks are altered in Alzheimer’s and other neurodegenerative diseases. Alzheimers Dement 14, 352–366 (2018).

17. Allen, M. et al. Divergent brain gene expression patterns associate with distinct cell-specific tau neuropathology traits in progressive supranuclear palsy. Acta Neuropathol 136, 709–727 (2018).

18. Nho, K. et al. Genome-wide transcriptome analysis identifies novel dysregulated genes implicated in Alzheimer’s pathology. Alzheimers Dement 16, 1213–1223 (2020).

19. Ma, Y. et al. Atlas of RNA editing events affecting protein expression in aged and Alzheimer’s disease human brain tissue. Nat Commun 12, 7035 (2021).

20. McKenzie, A. T. et al. Multiscale network modeling of oligodendrocytes reveals molecular components of myelin dysregulation in Alzheimer’s disease. Mol Neurodegener 12, 82 (2017).

21. Beckmann, N. D. et al. Multiscale causal networks identify VGF as a key regulator of Alzheimer’s disease. Nat Commun 11, 3942 (2020).

22. Seyfried, N. T. et al. A Multi-network Approach Identifies Protein-Specific Co-expression in Asymptomatic and Symptomatic Alzheimer’s Disease. Cell Syst 4, 60–72.e4 (2017).

23. Mostafavi, S. et al. A molecular network of the aging human brain provides insights into the pathology and cognitive decline of Alzheimer’s disease. Nat Neurosci 21, 811–819 (2018).

24. Johnson, E. C. B. et al. Large-scale deep multi-layer analysis of Alzheimer’s disease brain reveals strong proteomic disease-related changes not observed at the RNA level. Nat Neurosci 25, 213–225 (2022).

25. Kodam, P., Sai Swaroop, R., Pradhan, S. S., Sivaramakrishnan, V. & Vadrevu, R. Integrated multi-omics analysis of Alzheimer’s disease shows molecular signatures associated with disease progression and potential therapeutic targets. Sci. Rep. 13, 3695 (2023).

26. Zhang, J. et al. Integrative multi-omics analysis reveals the critical role of the PBXIP1 gene in Alzheimer’s disease. Aging Cell 23, e14044 (2024).

27. Liu, N. et al. Identification of novel prognostic biomarkers by integrating multi-omics data in gastric cancer. BMC Cancer 21, 460 (2021).

28. Jin, X. et al. Prioritization of therapeutic targets for cancers using integrative multi-omics analysis. Hum Genomics 18, 42 (2024).

29. Park, J.-C. et al. Multi-Omics-Based Autophagy-Related Untypical Subtypes in Patients with Cerebral Amyloid Pathology. Adv. Sci. 9, e2201212 (2022).

30. Iturria-Medina, Y. et al. Unified epigenomic, transcriptomic, proteomic, and metabolomic taxonomy of Alzheimer’s disease progression and heterogeneity. Sci Adv 8, eabo6764 (2022).

31. Iturria-Medina, Y., et al. Translating the Post-Mortem Brain Multi-Omics Molecular Taxonomy of Alzheimer’s Dementia to Living Humans. bioRxiv (2025) doi:10.1101/2025.03.20.644323.

32. Argelaguet, R. et al. Multi-Omics Factor Analysis-a framework for unsupervised integration of multi-omics data sets. Mol. Syst. Biol. 14, e8124 (2018).

33. Argelaguet, R. et al. MOFA+: a statistical framework for comprehensive integration of multi-modal single-cell data. Genome Biol 21, 111 (2020).

34. Strefeler, A. et al. Molecular insights into sex-specific metabolic alterations in Alzheimer’s mouse brain using multi-omics approach. Alzheimers. Res. Ther. 15, 8 (2023).

35. Bennett, D. A. et al. Religious orders study and rush memory and aging project. J. Alzheimers. Dis. 64, S161–S189 (2018).

36. Green, G. S. et al. Cellular communities reveal trajectories of brain ageing and Alzheimer’s disease. Nature 633, 634–645 (2024).

37. Bionetworks, S. Agora AD Knowledge Portal. (Sage Bionetworks, 2023). doi:10.57718/AGORA-ADKNOWLEDGEPORTAL.

38. Green, G. S. et al. Cellular communities reveal trajectories of brain ageing and Alzheimer’s disease. Nature 633, 634–645 (2024).

39. Houten, S. M., Violante, S., Ventura, F. V. & Wanders, R. J. A. The Biochemistry and Physiology of Mitochondrial Fatty Acid β-Oxidation and Its Genetic Disorders. Annu Rev Physiol 78, 23–44 (2016).

40. Batra, R. et al. The landscape of metabolic brain alterations in Alzheimer’s disease. Alzheimers. Dement. (2022) doi:10.1002/alz.12714.

41. Aaes, T. L., Burgoa Cardás, J. & Ravichandran, K. S. Defining solute carrier transporter signatures of murine immune cell subsets. Front Immunol 14, 1276196 (2023).

42. Chen, R. & Chen, L. Solute carrier transporters: emerging central players in tumour immunotherapy. Trends Cell Biol 32, 186–201 (2022).

43. Iwasaki, M. et al. BAG3 regulates motility and adhesion of epithelial cancer cells. Cancer Res 67, 10252–10259 (2007).

44. Marzullo, L., Turco, M. C. & De Marco, M. The multiple activities of BAG3 protein: Mechanisms. Biochim Biophys Acta Gen Subj 1864, 129628 (2020).

45. Stürner, E. & Behl, C. The Role of the Multifunctional BAG3 Protein in Cellular Protein Quality Control and in Disease. Front Mol Neurosci 10, 177 (2017).

46. Johnson, E. C. B. et al. Large-scale deep multi-layer analysis of Alzheimer’s disease brain reveals strong proteomic disease-related changes not observed at the RNA level. Nat Neurosci 25, 213–225 (2022).

47. Green, G. S. et al. Cellular communities reveal trajectories of brain ageing and Alzheimer’s disease. Nature 633, 634–645 (2024).

48. Green, G. S. et al. Cellular communities reveal trajectories of brain ageing and Alzheimer’s disease. Nature 633, 634–645 (2024).

49. Johnson, E. C. B. et al. Large-scale deep multi-layer analysis of Alzheimer’s disease brain reveals strong proteomic disease-related changes not observed at the RNA level. Nat Neurosci 25, 213–225 (2022).

50. Farid, M. M., Yang, X., Kuboyama, T. & Tohda, C. Trigonelline recovers memory function in Alzheimer’s disease model mice: evidence of brain penetration and target molecule. Sci Rep 10, 16424 (2020).

51. Gaiteri, C. et al. Identifying robust communities and multi-community nodes by combining top-down and bottom-up approaches to clustering. Sci. Rep. 5, 16361 (2015).

52. Si, Z.-Z. et al. Targeting neuroinflammation in Alzheimer’s disease: from mechanisms to clinical applications. Neural Regen Res 18, 708–715 (2023).

53. Onyango, I. G., Jauregui, G. V., Čarná, M., Bennett, J. P., Jr & Stokin, G. B. Neuroinflammation in Alzheimer’s Disease. Biomedicines 9, (2021).

54. Rajesh, Y. & Kanneganti, T.-D. Innate Immune Cell Death in Neuroinflammation and Alzheimer’s Disease. Cells 11, (2022).

55. Bamburg, J. R. & Bloom, G. S. Cytoskeletal pathologies of Alzheimer disease. Cell Motil Cytoskeleton 66, 635–649 (2009).

56. Barmaki, H., Nourazarian, A. & Khaki-Khatibi, F. Proteostasis and neurodegeneration: a closer look at autophagy in Alzheimer’s disease. Front Aging Neurosci 15, 1281338 (2023).

57. Scheff, S. W., Price, D. A., Schmitt, F. A., DeKosky, S. T. & Mufson, E. J. Synaptic alterations in CA1 in mild Alzheimer disease and mild cognitive impairment. Neurology 68, 1501–1508 (2007).

58. Ownby, R. L., Crocco, E., Acevedo, A., John, V. & Loewenstein, D. Depression and risk for Alzheimer disease: systematic review, meta-analysis, and metaregression analysis. Arch Gen Psychiatry 63, 530–538 (2006).

59. Boyle, P. A. et al. Attributable risk of Alzheimer’s dementia attributed to age-related neuropathologies. Ann Neurol 85, 114–124 (2019).

60. Eteleeb, A. M. et al. Brain high-throughput multi-omics data reveal molecular heterogeneity in Alzheimer’s disease. PLoS Biol 22, e3002607 (2024).

61. Li, M.-M. et al. Programmed cell death signatures-driven microglial transformation in Alzheimer’s disease: single-cell transcriptomics and functional validation. Front Immunol 16, 1610717 (2025).

62. Lei, Z., Brizzee, C. & Johnson, G. V. W. BAG3 facilitates the clearance of endogenous tau in primary neurons. Neurobiol. Aging 36, 241–248 (2015).

63. Marzullo, L., Turco, M. C. & De Marco, M. The multiple activities of BAG3 protein: Mechanisms. Biochim Biophys Acta Gen Subj 1864, 129628 (2020).

64. Brenner, C. M. et al. BAG3: Nature’s Quintessential Multi-Functional Protein Functions as a Ubiquitous Intra-Cellular Glue. Cells 12, (2023).

65. Wang, J. et al. Bag3 Regulates Mitochondrial Function and the Inflammasome Through Canonical and Noncanonical Pathways in the Heart. JACC Basic Transl Sci 8, 820–839 (2023).

66. Brenner, C. M. et al. BAG3: Nature’s Quintessential Multi-Functional Protein Functions as a Ubiquitous Intra-Cellular Glue. Cells 12, (2023).

67. Fontanella, B. et al. The co-chaperone BAG3 interacts with the cytosolic chaperonin CCT: new hints for actin folding. Int J Biochem Cell Biol 42, 641–650 (2010).

68. Franceschelli, S. et al. BAG3 Protein Is Involved in Endothelial Cell Response to Phenethyl Isothiocyanate. Oxid Med Cell Longev 2018, 5967890 (2018).

69. Marzullo, L., Turco, M. C. & De Marco, M. The multiple activities of BAG3 protein: Mechanisms. Biochim Biophys Acta Gen Subj 1864, 129628 (2020).

70. Ying, Z.-M. et al. BAG3 promotes autophagy and suppresses NLRP3 inflammasome activation in Parkinson’s disease. Ann Transl Med 10, 1218 (2022).

71. Brenner, C. M. et al. BAG3: Nature’s Quintessential Multi-Functional Protein Functions as a Ubiquitous Intra-Cellular Glue. Cells 12, (2023).

72. Franceschelli, S. et al. Bag3 gene expression is regulated by heat shock factor 1. J Cell Physiol 215, 575–577 (2008).

73. Meriin, A. B. et al. Hsp70-Bag3 complex is a hub for proteotoxicity-induced signaling that controls protein aggregation. Proc Natl Acad Sci U S A 115, E7043–E7052 (2018).

74. Hiebel, C. et al. BAG3 Proteomic Signature under Proteostasis Stress. Cells 9, (2020).

75. Beckmann, N. D. et al. Multiscale causal networks identify VGF as a key regulator of Alzheimer’s disease. Nat Commun 11, 3942 (2020).

76. Jinwal, U. K. et al. The Hsp90 cochaperone, FKBP51, increases Tau stability and polymerizes microtubules. J Neurosci 30, 591–599 (2010).

77. Blair, L. J. et al. Accelerated neurodegeneration through chaperone-mediated oligomerization of tau. J Clin Invest 123, 4158–4169 (2013).

78. Bozdagi, O. et al. The neurotrophin-inducible gene Vgf regulates hippocampal function and behavior through a brain-derived neurotrophic factor-dependent mechanism. J Neurosci 28, 9857–9869 (2008).

79. Bozdagi, O. et al. The neurotrophin-inducible gene Vgf regulates hippocampal function and behavior through a brain-derived neurotrophic factor-dependent mechanism. J Neurosci 28, 9857–9869 (2008).

80. Beckmann, N. D. et al. Multiscale causal networks identify VGF as a key regulator of Alzheimer’s disease. Nat Commun 11, 3942 (2020).

81. Buettner, F. & Theis, F. J. A novel approach for resolving differences in single-cell gene expression patterns from zygote to blastocyst. Bioinformatics 28, i626–i632 (2012).

82. Argelaguet, R. et al. Multi-Omics Factor Analysis-a framework for unsupervised integration of multi-omics data sets. Mol. Syst. Biol. 14, e8124 (2018).

83. Bennett, D. A., Schneider, J. A., Bienias, J. L., Evans, D. A. & Wilson, R. S. Mild cognitive impairment is related to Alzheimer disease pathology and cerebral infarctions. Neurology 64, 834–841 (2005).

84. Bennett, D. A., Schneider, J. A., Arvanitakis, Z. & Wilson, R. S. Overview and findings from the religious orders study. Curr. Alzheimer Res. 9, 628–645 (2012).

85. Bennett, D. A. et al. Decision rules guiding the clinical diagnosis of Alzheimer’s disease in two community-based cohort studies compared to standard practice in a clinic-based cohort study. Neuroepidemiology 27, 169–176 (2006).

86. Wilson, R. S., Boyle, P. A., Yang, J., James, B. D. & Bennett, D. A. Early life instruction in foreign language and music and incidence of mild cognitive impairment. Neuropsychology 29, 292–302 (2015).

87. De Jager, P. L. et al. A genome-wide scan for common variants affecting the rate of age-related cognitive decline. Neurobiol. Aging 33, 1017.e1–15 (2012).

88. Boyle, P. A. et al. To what degree is late life cognitive decline driven by age-related neuropathologies? Brain 144, 2166–2175 (2021).

89. Bennett, D. A. et al. Natural history of mild cognitive impairment in older persons. Neurology 59, 198–205 (2002).

90. Bennett, D. A. et al. Decision rules guiding the clinical diagnosis of Alzheimer’s disease in two community-based cohort studies compared to standard practice in a clinic-based cohort study. Neuroepidemiology 27, 169–176 (2006).

91. Schneider, J. A., Arvanitakis, Z., Bang, W. & Bennett, D. A. Mixed brain pathologies account for most dementia cases in community-dwelling older persons. Neurology 69, 2197–2204 (2007).

92. Vialle, R. A. et al. Structural variants linked to Alzheimer’s disease and other common age-related clinical and neuropathologic traits. Genome Med 17, 20 (2025).

93. Bennett, D. A. et al. Neuropathology of older persons without cognitive impairment from two community-based studies. Neurology 66, 1837–1844 (2006).

94. Wilson, R. S., Arnold, S. E., Schneider, J. A., Tang, Y. & Bennett, D. A. The relationship between cerebral Alzheimer’s disease pathology and odour identification in old age. J. Neurol. Neurosurg. Psychiatry 78, 30–35 (2007).

95. Bennett, D. A. et al. Apolipoprotein E epsilon4 allele, AD pathology, and the clinical expression of Alzheimer’s disease. Neurology 60, 246–252 (2003).

96. Bennett, D. A., Schneider, J. A., Tang, Y., Arnold, S. E. & Wilson, R. S. The effect of social networks on the relation between Alzheimer’s disease pathology and level of cognitive function in old people: a longitudinal cohort study. Lancet Neurol. 5, 406–412 (2006).

97. Kapasi, A. et al. High-throughput digital quantification of Alzheimer disease pathology and associated infrastructure in large autopsy studies. J. Neuropathol. Exp. Neurol. 82, 976–986 (2023).

98. Wilson, R. S., Arnold, S. E., Schneider, J. A., Tang, Y. & Bennett, D. A. The relationship between cerebral Alzheimer’s disease pathology and odour identification in old age. J. Neurol. Neurosurg. Psychiatry 78, 30–35 (2007).

99. Nag, S. et al. TDP-43 pathology in anterior temporal pole cortex in aging and Alzheimer’s disease. Acta Neuropathol Commun 6, 33 (2018).

100. Schneider, J. A., Arvanitakis, Z., Bang, W. & Bennett, D. A. Mixed brain pathologies account for most dementia cases in community-dwelling older persons. Neurology 69, 2197–2204 (2007).

101. Nag, S. et al. Hippocampal sclerosis and TDP-43 pathology in aging and Alzheimer disease. Ann Neurol 77, 942– 952 (2015).

102. Schneider, J. A. et al. The apolipoprotein E epsilon4 allele increases the odds of chronic cerebral infarction [corrected] detected at autopsy in older persons. Stroke 36, 954–959 (2005).

103. Arvanitakis, Z., Leurgans, S. E., Barnes, L. L., Bennett, D. A. & Schneider, J. A. Microinfarct pathology, dementia, and cognitive systems. Stroke 42, 722–727 (2011).

104. Arvanitakis, Z. et al. The Relationship of Cerebral Vessel Pathology to Brain Microinfarcts. Brain Pathol. 27, 77–85 (2017).

105. Buchman, A. S., Leurgans, S. E., Nag, S., Bennett, D. A. & Schneider, J. A. Cerebrovascular disease pathology and parkinsonian signs in old age. Stroke 42, 3183–3189 (2011).

106. Love, S. et al. Development, appraisal, validation and implementation of a consensus protocol for the assessment of cerebral amyloid angiopathy in post-mortem brain tissue. Am. J. Neurodegener. Dis. 3, 19–32 (2014).

107. Boyle, P. A. et al. Cerebral amyloid angiopathy and cognitive outcomes in community-based older persons. Neurology 85, 1930–1936 (2015).

108. Buchman, A. S. et al. Spinal motor neurons and motor function in older adults. J. Neurol. 266, 174–182 (2019).

109. Buchman, A. S., Wilson, R. S., Leurgans, S. E., Bennett, D. A. & Barnes, L. L. Change in motor function and adverse health outcomes in older African-Americans. Exp Gerontol 70, 71–77 (2015).

110. Buchman, A. S. et al. Nigral pathology and parkinsonian signs in elders without Parkinson disease. Ann. Neurol. 71, 258–266 (2012).

111. Buchman, A. S. et al. Nigral pathology and parkinsonian signs in elders without Parkinson disease. Ann. Neurol. 71, 258–266 (2012).

112. Buchman, A. S. et al. Person-specific contributions of brain pathologies to progressive parkinsonism in older adults. J. Gerontol. A Biol. Sci. Med. Sci. 76, 615–621 (2021).

113. Katz, S. & Akpom, C. A. A measure of primary sociobiological functions. Int J Health Serv 6, 493–508 (1976).

114. Boyle, P. A., Buchman, A. S., Wilson, R. S., Bienias, J. L. & Bennett, D. A. Physical activity is associated with incident disability in community-based older persons. J Am Geriatr Soc 55, 195–201 (2007).

115. Buchman, A. S., Boyle, P. A., Leurgans, S. E., Evans, D. A. & Bennett, D. A. Pulmonary function, muscle strength, and incident mobility disability in elders. Proc Am Thorac Soc 6, 581–587 (2009).

116. Buchman, A. S., Boyle, P. A., Wilson, R. S., Tang, Y. & Bennett, D. A. Frailty is associated with incident Alzheimer’s disease and cognitive decline in the elderly. Psychosom. Med. 69, 483–489 (2007).

117. Buchman, A. S., Wilson, R. S., Bienias, J. L. & Bennett, D. A. Change in frailty and risk of death in older persons. Exp. Aging Res. 35, 61–82 (2009).

118. Kohout, F. J., Berkman, L. F., Evans, D. A. & Cornoni-Huntley, J. Two shorter forms of the CES-D (Center for Epidemiological Studies Depression) depression symptoms index. J. Aging Health 5, 179–193 (1993).

119. Wilson, R. S. et al. Depressive symptoms, cognitive decline, and risk of AD in older persons. Neurology 59, 364–370 (2002).

120. Wilson, R. S. et al. Clinical-pathologic study of depressive symptoms and cognitive decline in old age. Neurology 83, 702–709 (2014).

121. Bennett, D. A., Wilson, R. S., Schneider, J. A., Bienias, J. L. & Arnold, S. E. Cerebral infarctions and the relationship of depression symptoms to level of cognitive functioning in older persons. Am. J. Geriatr. Psychiatry 12, 211–219 (2004).

122. Yu, L. et al. ’523 variant and cognitive decline in older persons with ε3/3 genotype. Neurology 88, 661–668 (2017).

123. Roses, A. D. et al. A TOMM40 variable-length polymorphism predicts the age of late-onset Alzheimer’s disease. Pharmacogenomics J. 10, 375–384 (2010).

124. Yu, L. et al. ’523 variant and cognitive decline in older persons with ε3/3 genotype. Neurology 88, 661–668 (2017).

125. Yu, L. et al. ’523 variant and cognitive decline in older persons with ε3/3 genotype. Neurology 88, 661–668 (2017).

126. Klein, H.-U. et al. Epigenome-wide study uncovers large-scale changes in histone acetylation driven by tau pathology in aging and Alzheimer’s human brains. Nat. Neurosci. 22, 37–46 (2018).

127. Tasaki, S. et al. Inferring protein expression changes from mRNA in Alzheimer’s dementia using deep neural networks. Nat. Commun. 13, 655 (2022).

128. Green, G. S. et al. Cellular communities reveal trajectories of brain ageing and Alzheimer’s disease. Nature 633, 634–645 (2024).

129. Fujita, M. et al. Cell subtype-specific effects of genetic variation in the Alzheimer’s disease brain. Nat. Genet. 56, 605–614 (2024).

130. Green, G. S. et al. Cellular communities reveal trajectories of brain ageing and Alzheimer’s disease. Nature 633, 634–645 (2024).

131. Johnson, E. C. B. et al. Large-scale proteomic analysis of Alzheimer’s disease brain and cerebrospinal fluid reveals early changes in energy metabolism associated with microglia and astrocyte activation. Nat. Med. 26, 769–780 (2020).

132. Batra, R. et al. The landscape of metabolic brain alterations in Alzheimer’s disease. Alzheimers. Dement. (2022) doi:10.1002/alz.12714.

133. Argelaguet, R. et al. Multi-Omics Factor Analysis-a framework for unsupervised integration of multi-omics data sets. Mol. Syst. Biol. 14, e8124 (2018).

134. Argelaguet, R. et al. Multi-Omics Factor Analysis-a framework for unsupervised integration of multi-omics data sets. Mol. Syst. Biol. 14, e8124 (2018).

135. R Core Team. R: A Language and Environment for Statistical Computing. (Vienna, Austria, 2023).

136. Gelman, A. & Su, Y.-S. arm: Data Analysis Using Regression and Multilevel/Hierarchical Models. Preprint at https://CRAN.R-project.org/package=arm (2024).

137. Christensen, R. H. B. ordinal---Regression Models for Ordinal Data. Preprint at https://CRAN.R-project.org/package=ordinal (2023).

138. Korotkevich, G. et al. Fast gene set enrichment analysis. bioRxiv (2016) doi:10.1101/060012.

139. Milacic, M. et al. The Reactome Pathway Knowledgebase 2024. Nucleic Acids Res. 52, D672–D678 (2024).

140. Gene Ontology Consortium et al. The Gene Ontology knowledgebase in 2023. Genetics 224, (2023).

141. Ashburner, M. et al. Gene ontology: tool for the unification of biology. The Gene Ontology Consortium. Nat. Genet. 25, 25–29 (2000).

142. Dolgalev, I. msigdbr: MSigDB Gene Sets for Multiple Organisms in a Tidy Data Format. Preprint at https://CRAN.R-project.org/package=msigdbr (2022).

143. Bellenguez, C. et al. New insights into the genetic etiology of Alzheimer’s disease and related dementias. Nat Genet 54, 412–436 (2022).

144. Chen, W.-T. et al. Spatial Transcriptomics and In Situ Sequencing to Study Alzheimer’s Disease. Cell 182, 976–991.e19 (2020).

145. Keren-Shaul, H. et al. A Unique Microglia Type Associated with Restricting Development of Alzheimer’s Disease. Cell 169, 1276–1290.e17 (2017).

146. Srinivasan, K. et al. Alzheimer’s Patient Microglia Exhibit Enhanced Aging and Unique Transcriptional Activation. Cell Rep 31, 107843 (2020).

147. Sayols, S. rrvgo: a Bioconductor package for interpreting lists of Gene Ontology terms. MicroPubl Biol 2023, (2023).

148. Lancichinetti, A., Fortunato, S. & Radicchi, F. Benchmark graphs for testing community detection algorithms. Phys. Rev. E Stat. Nonlin. Soft Matter Phys. 78, 046110 (2008).

149. Mostafavi, S. et al. A molecular network of the aging human brain provides insights into the pathology and cognitive decline of Alzheimer’s disease. Nat Neurosci 21, 811–819 (2018).

150. Ng, B. et al. Integration across biophysical scales identifies molecular and cellular correlates of person-to-person variability in human brain connectivity. Nat. Neurosci. 27, 2240–2252 (2024).

