## Supplementary Material for "Integration of aged brain multi-omics reveals cross-system mechanisms underlying Alzheimer’s disease heterogeneity"

### Table Of Contents

**Supplementary Notes** 4

**Detailed characterization of AD-related factors** 4

Factor 1 4
Factor 2 4
Factor 3 5
Factor 4 5
Factor 12 6
Factor 14 6
Factor 26 7
Factor 42 8

**Detailed description of ADRD phenotypes analyzed** 9

NIA-Reagan criteria (modified) 9
Tangle density 9
Neurofibrillary tangle burden 9
β-amyloid level 9
Neuritic plaque burden 9
Diffuse plaque burden 9
Global AD pathology burden 10
TDP-43 stage 10
Lewy body disease 10
Parkinson's disease pathology 10
Hippocampal sclerosis 10

**Alzheimer’s dementia and cognitive function** 10

Clinical cognition status (AD dementia and Mild cognitive impairment) 11
Global cognition function 11
Cognitive decline 11
Cognitive resilience 11

**Parkinsonism and motor function** 11

Frailty 11
Motor function 11
Bradykinesia 12
Rigidity 12
Gait 12
Tremor 12
Motor gait 12
Motor dexterity 13
Motor hand strength 13
Global Parkinsonian score 13

**Vascular pathologies** 13

Arteriosclerosis 13
Cerebral atherosclerosis 13
Cerebral amyloid angiopathy (CAA) 14
Cerebral infarctions (gross) 14

### Supplementary Notes

#### Detailed characterization of AD-related factors

##### Factor 1

Factor 1 was positively associated with AD dementia (adj. P = 1.84 x $10^{-7}$), NIA-Reagan score (adj. P = 1.18 x $10^{-4}$), the APOEε4 allele (adj. P = 1.44 x $10^{-2}$), and most pathological traits (except for Lewy body disease and PD pathology). This factor was also negatively associated with global cognitive function (adj. P = 1.74 x $10^{-7}$) and cognitive decline (adj. P = 2.04 x $10^{-6}$) (**Figure 1d**). RNA (DLPFC) was the view that contributed most to Factor 1 variance explained (4.10%), whereas H3K9ac contributed the least (1.29%) (**Figure 1c**). Nevertheless, relevant terms were identified across most views. Transcriptomics in the AC and DLPFC regions were positively enriched with proteostasis and heat shock response terms, such as chaperone-mediated protein folding (adj. P = 4.05 x $10^{-3}$) and HSF1 activation (adj. P = 4.05 x $10^{-3}$). These views also showed positive enrichment for End.5 markers (adj. P = 4.56 x $10^{-5}$), a subpopulation of endothelial cells that expresses heat and oxidative stress response, tau binding, and necroptosis, as well as AD risk genes [1]. Proteomics was positively enriched for a variety of terms, including chaperone-mediated protein folding (adj. P = 5.06 x $10^{-5}$), complement activation (adj. P = 8.64 x $10^{-10}$), and the co-expression protein modules Prot-M20 (related to RNA splicing, adj. P = 1.55 x $10^{-39}$) and Prot-M7 (MAPK metabolism-related, adj. P = 1.18 x $10^{-30}$). Proteomics also showed several negatively enriched terms, such as mitochondrial gene expression (adj. P = 1.10 x $10^{-10}$), ribosomal subunit (adj. P = 3.15 x $10^{-27}$), and Prot-M12 (related to the cytoskeleton, adj. P = 2.30 x $10^{-22}$) (**Supplementary Figure S9a**).

##### Factor 2

Factor 2 was negatively associated with AD dementia (adj. P = 1.44 x $10^{-3}$), NIA-Reagan score (adj. P = 7.51 x $10^{-3}$), six out of ten pathological traits, and was the only factor negatively associated with gross cerebral infarctions (adj. P = 2.22 x $10^{-3}$). It was also positively associated with global cognitive function (adj. P = 2.52 x $10^{-3}$) and cognitive decline (adj. P = 8.50 x $10^{-4}$) (**Figure 1d**). The view that contributed the most to Factor 2's total variance was cell types (6.41%), while the variance explained by the other views ranged from 3.22% (PCG mRNA) to 0.26% (H3K9ac) (**Figure 1c**). Still, all views presented relevant terms. All views were positively enriched for immune-related terms, such as immune response (adj. P = 1.25 x $10^{-45}$), response to cytokine (adj. P = 2.15 x $10^{-19}$), and regulation of phagocytosis (adj. P = 5.07 x $10^{-7}$). H3K9ac and transcriptomics were also positively enriched for terms related to interleukin signaling, such as signaling by interleukins (adj. P = 1.10 x $10^{-17}$) and interleukin-4 and interleukin-13 signaling (adj. P = 9.64 x $10^{-10}$). Most views were negatively enriched for RNA module M109 (adj. P = 1.32 x $10^{-5}$), a co-expression gene module strongly related to cognitive decline, β-amyloid pathology, and clinical diagnosis of AD dementia. For cell types, most views were enriched with different subtypes of microglia: Mic.7 (reacting, adj. P = 7.80 x $10^{-9}$) and Mic.10 (enhanced-redox, adj. P = 2.11 x $10^{-10}$) were enriched in transcriptomics and proteomics, while Mic.14 (interferon response, adj. P = 6.05 x $10^{-5}$) was enriched in all views except RNA (DLPFC) (**Supplementary Figure S10a**).

#### Factor 3

Factor 3 showed a negative association with Alzheimer's dementia (adj. P = 2.89 x $10^{-4}$), NIA-Reagan score (adj. P = 1.26 x $10^{-2}$), and most of the pathological traits. It was the only factor to show a negative association with the TOMM40'523 genotype (adj. P = 2.36 x $10^{-2}$). Factor 3 was positively enriched for global cognitive function (adj. P = 4.23 x $10^{-4}$), cognitive decline (adj. P = 3.05 x $10^{-4}$), and cognitive resilience (adj. P = 2.36 x $10^{-2}$) (**Figure 1d**). The view that contributed the most to Factor 3's total variance explained was RNA (DLPFC) (10.91%); the variance explained by the other views ranged from 2.24% (PCG mRNA) to 0.06% (H3K9ac) (**Figure 1c**). Despite the low contribution of H3K9ac, it still presented relevant terms, namely innate immune system (adj. P = 2.14 x $10^{-3}$), interleukin-4 and interleukin-13 signaling (adj. P = 2.14 x $10^{-3}$), and cytokine signaling in immune system (adj. P = 3.59 x $10^{-3}$), among others. mRNA from DLPFC and PCG (the top contributors), along with proteomics, were negatively enriched for several immune-related terms, such as inflammatory response (adj. P = 1.76 x $10^{-40}$), cytokine production (adj. P = 2.36 x $10^{-30}$), and immune effector process (adj. P = 1.50 x $10^{-38}$). These views were also negatively enriched for PIGs (adj. P = 1.08 x $10^{-6}$), a co-expression network of plaque-induced genes [2], and for Min.15 (high expression of inflammation and stress signatures, adj. P = 2.23 x $10^{-12}$). H3K9ac, transcriptomics, and proteomics were negatively enriched for monocytes (adj. P = 1.83 x $10^{-37}$) and macrophages (adj. P = 2.40 x $10^{-14}$) (**Supplementary Figure S11a**). Metabolomics was enriched for the bioenergetic pathway (adj. P = 2.07 x $10^{-2}$) (**Supplementary Table S1**).

#### Factor 4

Factor 4 was negatively associated with AD dementia (adj. P = 2.34 x $10^{-6}$), NIA-Reagan score (adj. P = 5.78 x $10^{-4}$), and positively associated with global cognitive function (adj. P = 1.74 x $10^{-7}$). This factor was also associated with all motor and disability traits except for the tremor score. Additionally, it was the only factor, among the AD-related factors, negatively associated with Lewy body disease (adj. P = 1.71 x $10^{-2}$), PD pathology (adj. P = 8.27 x $10^{-5}$), mobility disability (adj. P = 1.78 x $10^{-3}$), and depressive symptoms (mCES-D) (adj. P = 9.09 x $10^{-3}$) (**Figure 1d**). Metabolomics was the view that contributed the most to Factor 4 variance explained (4.49%), whereas H3K9ac contributed the least (0.37%) (**Figure 1c**). Nevertheless, relevant terms were identified across all views. H3K9ac was positively enriched for immune-related processes, such as interferon-gamma signaling (adj. P = 3.96 x $10^{-2}$), cellular respiration activity, including respiratory electron transport (adj. P = 8.33 x $10^{-3}$), and the co-expression protein module associated with oligodendrocyte myelination (adj. P = 7.79 x $10^{-5}$). H3K9ac also showed positive enrichment for Mic.14 (adj. P = 7.79 x $10^{-5}$), a microglial subtype characterized by an interferon response [1]. Transcriptomics in the AC region was positively enriched for terms related to the GPCR superfamily, namely GPCR ligand binding (adj. P = 2.91 x $10^{-3}$) and class A1 Rhodopsin-like receptors (adj. P = 3.41 x $10^{-2}$). Other relevant terms associated with the AC region included cell-cell signaling (adj. P = 5.38 x $10^{-4}$) and ion transport (adj. P = 6.93 x $10^{-4}$). RNA DLPFC and PCG shared terms primarily associated with immune system processes, such as leukocyte-mediated immunity (adj. P = 1.70 x $10^{-6}$) and inflammatory response (adj. P = 4.68 x $10^{-5}$), as well as with the GPCR superfamily, including GPCR ligand binding (adj. P = 1.74 x $10^{-3}$) and signaling by GPCR (adj. P = 1.38 x $10^{-3}$). Both views were also positively enriched for Mic.14 (adj. P = 8.7 x $10^{-5}$) and negatively enriched for Ast.8 (adj. P = 1.37 x $10^{-3}$), an astrocyte subtype characterized by the expression of heat stress and DNA damage, calcium, and sterol metabolism genes [1]. Proteomics was positively enriched for terms related to the ribosome, including structural constituent of the ribosome (adj. P = 1.38 x $10^{-11}$) and the cytosolic large ribosomal subunit (adj. P = 3.86 x $10^{-7}$). Other relevant terms included cytoplasmic translation (adj. P = 7.53 x $10^{-13}$) and co-expression protein modules related to the cytoskeleton (adj. P = 1.54 x $10^{-10}$) and MAPK metabolism (adj. P = 1.92 x $10^{-6}$) (**Supplementary Figure S12a**). Finally, metabolomics was enriched for lysophospholipid (adj. P = 3.31 x $10^{-3}$) and long-chain polyunsaturated fatty acid (n-3 and n-6) (adj. P = 1.20 x $10^{-2}$) (**Supplementary Table S1**).

The features with the highest absolute weights (**Supplementary Figure S12b-c**) for Factor 4 included several genes related to AD. Among the features with high positive weights, many are considered protective against AD. For instance, the gene *CX3CR1* encodes a microglial chemokine receptor that modulates microglial activation. *CX3CR1* deficiency exacerbates AD-related neuronal and behavioral impairments in a mouse model, indicating that it is a protective gene against AD-related cognitive deficits [3]. As for metabolites, trigonelline has shown a protective function in mice by ameliorating axonal and dendritic atrophy in cortical neurons treated with Aβ [4]. Nicotinamide riboside also demonstrated beneficial effects in an 8-week trial, where patients supplemented with nicotinamide riboside exhibited lower pTau217 concentrations, whereas those in the placebo group showed an increase [5]. Among the features with high negative weights, several contribute to AD progression. For example, *FKBP5* (FK506 binding protein 5) prevents tau degradation in mice by forming a chaperone complex with HSP90 that blocks tau clearance [5,6]. Similarly, *IGFBP3* (insulin-like growth factor-binding protein 3) encodes IGFBP-3, a protein released by astrocytes that induces tau phosphorylation in neurons [7] (**Supplementary Figure S12b**).

##### Factor 12

Factor 12 was negatively associated with AD dementia (adj. P = 2.88 x $10^{-2}$), NIA-Reagan score (adj. P = 2.22 x $10^{-3}$), and six pathological traits. It showed no association with the other categories (**Figure 1d**). Among all views, H3K9ac contributed by far the most to the explained variance (5.34%), while the second-biggest contributor (mRNA PCG) explained only 0.21% (**Figure 1c**). H3K9ac was enriched for a variety of terms, including the co-expression RNA modules RNA-M12 (adj. P = 9.83 x $10^{-7}$) and RNA-M114 (adj. P = 3.28 x $10^{-16}$), both associated with β-amyloid burden; RNA-M7, associated with PHF-tau tangle density (adj. P = 6.43 x $10^{-12}$); postsynapse (adj. P = 9.48 x $10^{-8}$); and others (**Supplementary Figure S13a**).

##### Factor 14

Factor 14 showed a positive association with Alzheimer's Dementia (adj. P = 1.33 x $10^{-12}$) and NIA-Reagan score (adj. P = 3.34 x $10^{-7}$), and a negative association with Global cognitive function (adj. P = 1.13 x $10^{-17}$) and Cognitive decline (adj. P = 1.67 x $10^{-15}$). It was positively associated with all pathological traits tested, except for hippocampal sclerosis, and had the strongest association with the presence of Lewy Bodies (adj. P = 2.17 x $10^{-4}$). It showed the only positive association with PD pathology (adj. P = 4.87 x $10^{-6}$) and major depressive disorder (adj. P = 2.78 x $10^{-2}$) among the selected factors (**Figure 1d**). The views that contributed the most to Factor's 14 total variance were proteomics (2.83%) and metabolomics (1.19%), while the variance explained by the other views ranged from 0.34% (AC mRNA) to 0.007% (cell type) (**Figure 1c**). Proteomics enriched with a diverse array of terms, some of the positive terms were the protein module Prot-M7 (involved with MAPK metabolism) (adj. P = 1.94 x $10^{-29}$) and Prot-M11 (related to cell interaction with the extracellular matrix) (adj. P = 3.72 x $10^{-26}$). Proteomics was also negatively enriched for mitochondrial gene expression (adj. P = 1.62 x $10^{-17}$), and modules Prot-M5 (related to post-synaptic density) (adj. P = 2.42 x $10^{-16}$) and Prot-M8 (engaged in protein transport) (adj. P = 1.68 x $10^{-25}$). Metabolomics enriched with energy metabolism, such as, glycolysis, gluconeogenesis, and pyruvate metabolism (adj. P = 1.25 x $10^{-2}$), Nicotinate and Nicotinamide Metabolism (adj. P = 1.65 x $10^{-2}$) and Purine Metabolism, Adenine containing (adj. P = 6.90 x $10^{-3}$). Metabolomics also enriched for Glutathione Metabolism (adj. P = 1.25 x $10^{-2}$), possibly related to antioxidant defense, and Histidine Metabolism (adj. P = 2.76 x $10^{-3}$), related to protein synthesis (**Supplementary Table S2**). Despite the low overall contributions of the remaining views, their top features also presented biologically relevant terms. For instance, H3K9ac explained 0.05% of its total variance but was enriched mostly with synapse related terms and RNA processing and splicing terms, such as metabolism of RNA (adj. P = 3.11 x $10^{-6}$), synapse (adj. P = 2.83 x $10^{-5}$) and co-expression protein modules such as Prot-M13 (related to RNA splicing) (adj. P = 2.48 x $10^{-3}$) and Prot-M1/M4 (related to synapses). mRNA from AC, PCG, and DLPFC explained 0.34%, 0.27,% and 0.33% of their data variance, respectively. AC region enriched mostly with terms related to cell signaling activity, such as cell-cell signaling (adj. P = 1.22 x $10^{-4}$) and PCG enriched with immune related terms, such as interleukin 4 and interleukin 13 signaling (adj. P = 4.23 x $10^{-2}$), and neuronal signaling pathways, namely NGF stimulated transcription (adj. P = 2.47 x $10^{-2}$). PCG also enriched with markers for the microglial subpopulation Mic.15 (adj. P = 6.97 x $10^{-4}$), which highly express inflammation and stress signatures. (**Supplementary Figure S14a**).

The features with highest absolute weights (**Supplementary Figure S14b-c**) for Factor 14 showed genes related to Alzheimer's disease, such as *Notch1*, which modulates the activity of the Amyloid-β precursor protein (APP) by downregulating the APP intracellular domain [8] and *SST*, which regulates the metabolism of amyloid β peptide through modulating proteolytic degradation [9]. Differences in cell proportions showed positive weights for two important subtypes of lipid-associated microglia: MIC.12 (associated with neocortical Aβ burden and tau burden) and MIC.13 (associated with neocortical Aβ burden, tau burden, and rapid rate of cognitive decline), and also OPC.3 [1]. Metabolites captured Uridine 5′-diphosphoglucose (UDP-Glucose), which is an agonist of the purinergic $P2Y_{14}$ receptor ($P2Y_{14}R$). Interactions between UDP-G and microglial $P2Y_{14}Rs$ seem to be involved in the modulation of neuroinflammation via cAMP-NLRP3, MAPKs, and STAT1 pathways [10] (**Supplementary Figure S14b**).

#### Factor 26

Factor 26 had the strongest negative association with Alzheimer's Dementia (adj. P = 5.42 x $10^{-9}$) and NIA-Reagan score (adj. P = 1.86 x $10^{-4}$), and the strongest positive association with Global cognitive function (adj. P = 2.16 x $10^{-12}$) and cognitive decline (adj. P = 1.19 x $10^{-10}$). It was associated negatively with most pathological traits, such as beta-amyloid (adj. P = 1.39 x $10^{-2}$), tangles density (adj. P =4.50 x $10^{-6}$), diffuse plaques (adj. P = 2.36 x $10^{-2}$), neuritic plaques (adj. P = 7.02 x $10^{-4}$) and others. It was also the only factor negatively associated with cerebral atherosclerosis (adj. P = 1.25 x $10^{-2}$) (**Figure 1d**). The variance explained for Factor 26 ranged from 0.86% (PCG mRNA) to 0.17% (cell type), with many biologically relevant terms identified in each view (**Figure 1c**). H3K9ac explained 0.18% of its total variance and enriched mainly with terms related to neuronal development, such as synapse (adj. P = 2.64 x $10^{-5}$), neurogenesis (adj. P = 1.40 x $10^{-11}$), and neuron projection (adj. P = 1.34 x $10^{-5}$). mRNA from AC, PCG and DLPFC explained 0.51%, 0.86%, and 0.39% of their data variance, respectively. AC region enriched mostly with terms related to cytokine signaling, including cytokine signaling in immune system (adj. P = 8.70 x $10^{-3}$) and vascular terms, like vasculature development (adj. P = 2.16 x $10^{-5}$). RNA DLPFC enriched with vasculature development (adj. P = 3.07 x $10^{-3}$), organic acid transport (adj. P = 6.56 x $10^{-3}$), among others. RNA PCG enriched with response to lipid (adj. P = 1.70 x $10^{-6}$), cytokine signaling in immune system (adj. P = 3.57 x $10^{-3}$), vasculature development (adj. P = 2.48 x $10^{-13}$), and more. Proteomics explained 0.68% of its data total variance and enriched with terms related to mRNA processes, such as mRNA metabolic process (adj. P = 5.95 x $10^{-6}$) and mRNA binding (adj. P = 3.85 x $10^{-6}$), it also enriched with protein folding (adj. P = 1.04 x $10^{-9}$) and M2-mitochondria (adj. P = 1.89 x $10^{-20}$) (**Supplementary Figure S15a**). Metabolomics accounted for 0.42% of its total data variance and enriched for energy metabolism (bioenergetic pathway, adj. P = 1.22 x $10^{-3}$) and lipid metabolism (Sphingomyelins, adj. P = 1.22 x $10^{-3}$) (**Supplementary Table S2**).

The features with the highest absolute weights (**Supplementary Figure S15b-c**) for factor 26 showed protective genes and biomarkers related to AD. For instance, *GFAP* is an astrocytic cytoskeletal protein and serves as an early marker for AD [11,12]. *VEGFA* reduces the recruitment of hyperreactive neutrophils into the brain by mediating endothelial Cdk5 activity in decreasing *CXCL1* secretion in the AD brain, therefore reducing neuroinflammation [13]. For cell type, MIC.6 is a reactive-like subpopulation of microglia, and it is associated with a constant level of A$\beta$, low/no neocortical tau, and a slower rate of cognitive decline [1]. As for the metabolites, Spermidate (N-Carboxyethyl-g-aminobutyric acid) is a derivative of gamma-aminobutyric acid (GABA) and has a positive association with Aβ levels and NIA Reagan score [13,14] (**Supplementary Figure S13b**).

##### Factor 42

Factor 42 was positively associated with Alzheimer's dementia (adj. P = 8.93 x $10^{-3}$), NIA-Reagan score (adj. P = 1.10 x $10^{-4}$), and most of the pathological traits. It was also associated with instrumental activities of daily living (adj. P = 9.09 x $10^{-3}$) (**Figure 1d**). Among the AD-related factors, it is the factor with the lowest total variance explained; nonetheless, some views presented relevant terms (**Figure 1c**). For instance, H3K9ac explained 0.05% of its total variance and was positively enriched for terms related to neural/synaptic components, such as synapse (adj. P = 6.48 x $10^{-13}$), neuron projection (adj. P = 2.61 x $10^{-8}$), and somatodendritic compartment (adj. P = 1.48 x $10^{-5}$). H3K9ac was also positively enriched for terms related to signal transduction, namely small GTPase-mediated signal transduction (adj. P = 2.61 x $10^{-8}$) and molecular transducer activity (adj. P = 4.54 x $10^{-5}$), and for terms related to cell regulation, including regulation of transport (adj. P = 7.15 x $10^{-6}$) and regulation of cell population proliferation (adj. P = 1.32 x $10^{-3}$). mRNA from the AC region explained 0.34% of its data variance and was positively enriched for vascular-related terms, including Peri.2 (pericyte subpopulation aligned with extracellular matrix, adj. P = 4.70

x $10^{-17}$) [1], arteriole (adj. P = 7.38 x $10^{-15}$), and endothelial cell apoptotic process (adj. P = 8.33 x $10^{-3}$); it was also negatively enriched for mitochondrial-related terms, in particular mitochondrial matrix (adj. P = 2.04 x $10^{-5}$) and the module of co-expression Prot-M2 (mitochondria-related) (adj. P = 4.46 x $10^{-3}$). Proteomics explained 0.14% of its total variance and was positively enriched for terms related to neural/synaptic structures, such as Prot-M22 (adj. P = 9.95 x $10^{-11}$) and Prot-M5 (adj. P = 4.32 x $10^{-2}$), both related to postsynaptic density; synapse (adj. P = 2.42 x $10^{-3}$); and somatodendritic compartment (adj. P = 5.18 x $10^{-3}$). It was also positively enriched for terms related to vasculature, such as positive regulation of vasoconstriction (adj. P = 1.59 x $10^{-3}$) and endothelial cell apoptotic process (adj. P = 4.20 x $10^{-3}$) (**Supplementary Figure S16a**). Finally, metabolites were enriched for leucine, isoleucine, and valine metabolism (adj. P = 2.40 x $10^{-3}$) and the bioenergetic pathway (adj. P = 3.29 x $10^{-6}$) (**Supplementary Table S2**).

#### Detailed description of ADRD phenotypes analyzed

##### NIA-Reagan criteria (modified)

A modified version of the NIA-Reagan diagnosis of AD criteria, where the neuropathological evaluation is done without knowledge of clinical information, including a diagnosis of dementia [15,16]. It varies from 1 (no likelihood of AD) to 4 (high likelihood of AD).

##### Tangle density

It is based on historical and/or digital data and is the mean of the square root transformation of the tangle score in 8 regions: Angular gyrus (AG), Anterior cingulate (ACC), Calcarine cortex (CC), Entorhinal cortex (EC), Hippocampus (HC), Inferior temporal (IT), Midfrontal gyrus (MFG) and Superior frontal (SFG) [17].

##### Neurofibrillary tangle burden

Determined by microscopic examination of silver-stained slides from midfrontal córtex (MFC), midtemporal córtex (MTC), inferior parietal córtex (IPC), EC, and HC. Each region is scaled by dividing the corresponding standard deviation and then averaged to obtain a summary measure for neurofibrillary tangle burden, which is then square-rooted [18].

##### β-amyloid level

The square root of the mean Aβ score of 8 regions (HC, EC, MFC, IT, AG, CC, ACC, and SFC). Aβ is identified by molecularly specific immunohistochemistry and quantified by image analysis [18].

##### Neuritic plaque burden

Determined by microscopic examination of silver-stained slides from 5 regions (EC, HC, MTC, IPC, MFC). Each region is scaled by dividing the corresponding standard deviation, then averaged and square-rooted to obtain a summary measure for neuritic plaque burden [18].

##### Diffuse plaque burden

Determined by microscopic examination of silver-stained slides from 5 regions (EC, HC, MTC, IPC, MFC). Each region is scaled by dividing the corresponding standard deviation, then averaged and square-rooted to obtain a summary measure for diffuse plaque burden [18].

##### Global AD pathology burden

A quantitative summary of AD pathology derived from neuritic plaques, diffuse plaques, and neurofibrillary tangles, as determined by microscopic examination of silver-stained slides from MFC, MTC, IPC, EC, and HC. Each regional count is scaled by dividing by the corresponding standard deviation and averaged to obtain summary measures. The 3 summary measures are then averaged to obtain the measure of global AD pathology [19,20].

##### TDP-43 stage

The presence of TAR DNA-binding protein 43 (TDP-43) cytoplasmic inclusions in neurons and glia by immunohistochemistry from 8 brain regions: amygdala, EC, HC CA1, HC dentate gyrus, anterior temporal pole cortex, MTC, Orbital frontal cortex, and MFC. Two stages of TDP-43 distribution are recognized according to its presence and distribution in the regions (No TDP-43 pathology or in amygdala only, and TDP-43 pathology extending beyond amygdala) [21].

##### Lewy body disease

Binary variable in which the presence of Lewy bodies is identified by α-synuclein immunohistochemistry in 7 regions: substantia nigra, entorhinal cortex, anterior cingulate cortex, midfrontal cortex, superior or middle temporal cortex, inferior parietal cortex, and amygdala. The McKeith criteria were modified to assess the presence of LB in different categories: not present, nigral-predominant, limbic-type, and neocortical-type. A dichotomized version of this variable is used, referring to Lewy bodies present or absent [22].

##### Parkinson's disease pathology

Parkinson’s disease (PD) pathology was defined as a binary variable based on the presence of Lewy bodies and the severity of neuronal loss in the substantia nigra. Neuronal loss in the substantia nigra was graded on a 4-point scale (0 = none/rare/scattered, 1 = mild, 2 = moderate, 3 = severe). PD pathology was defined as present (PD pathology = 1) if Lewy bodies were detected in at least one region and neuronal loss was moderate or severe. All other combinations were considered absent PD pathology (PD pathology = 0).

##### Hippocampal sclerosis

Hippocampal sclerosis is assessed unilaterally in a coronal section of the mid-hippocampus at the level of the lateral geniculate body. It is classified as absent or present based on severe neuronal loss and gliosis in CA1 and/or the subiculum. Neuronal loss and gliosis in each region are graded on a scale from 0 to 5, with higher values indicating greater severity (0 = none, 1 = mild, 2 = mild to moderate, 3 = moderate, 4 = moderate to severe, 5 = severe). Hippocampal sclerosis is identified when a grade of 5 (severe) is assigned [23].

##### Alzheimer’s dementia and cognitive function

#### Clinical cognition status (AD dementia and Mild cognitive impairment)

A modified version of a three-stage process that classifies participants' cognition status into 6 categories: No cognitive impairment (NCI), Mild cognitive impairment (MCI), MCI and another condition contributing to cognitive status, Alzheimer’s dementia, Alzheimer’s dementia and other condition contributing to cognitive status, and other dementia. This process involves a battery of 19 cognitive tests, a clinical judgment by a neuropsychologist, and diagnostic classification by a clinician [24,25]. The Clinical cognition status variable condenses the participants' status in only 3 categories instead of the original 6: NCI, MCI, and Alzheimer’s dementia. In our analysis, two binary statuses were used: Alzheimer’s dementia vs no dementia (MCI + NCI) and with cognitive impairment (AD + MCI) vs no cognitive impairment (NCI).

#### Global cognition function

It is a composite measure of global cognitive function comprising a battery of 19 cognitive performance tests. The test scores were converted to z-scores and averaged to yield a global cognitive function summary. Mean and standard deviation at baseline were used to compute the z-scores. A negative z-score means that the participant has an overall score that is lower than the average of the cohort at baseline [26].

#### Cognitive decline

It is the estimated person-specific rate of change in the global cognition function over time, controlled for demographics (age at baseline, sex, and years of education) [27].

#### Cognitive resilience

It is the estimated person-specific rate of change in the global cognition function over time, controlling for demographics and pathology (global AD pathology burden, β-amyloid level, PHF tau tangles, Lewy body disease, TDP-43 stage, gross chronic cerebral infarctions, chronic microinfarctions, hippocampal sclerosis, cerebral amyloid angiopathy, cerebral atherosclerosis, arteriolosclerosis [27].

#### Parkinsonism and motor function

#### Frailty

Frailty measure reflects multisystem vulnerability, based on four components: grip strength, timed walk, body composition (BMI), and fatigue. The measure was constructed by converting each raw component score into a z-score using baseline mean and standard deviation values from all participants. Grip strength and timed walk were obtained from the motor exam, with sex-specific standardization applied; grip strength was reversed so that lower strength (greater frailty) corresponded to higher z-scores, while timed walk was not reversed. BMI and fatigue were standardized across all participants regardless of sex. Fatigue was derived from two CES-D items assessing effort and motivation, scored 0–2 based on affirmative responses. The final frailty score was computed as the mean of the four z-scores, with higher values indicating greater frailty [28,29].

#### Motor function

A composite measure of global motor function was calculated using 10 metrics: purdue pegboard test (no. of pegs), finger-tapping test (taps/10 seconds), time and number of steps to cover a distance of 8 feet (1/seconds), the time and number of steps for 360-degree turn, leg and toe stand, grip strength, and pinch strength. The performance for each motor measure is converted to a score using the mean from all participants at baseline and averaging all the motor tests together [30]. For each individual, the annual rate of change in motor decline was estimated using a linear mixed effects model, controlling for age at baseline, sex, and years of education.

##### Bradykinesia

It is a measure of arm and leg agility determined by using a modified version of the motor portion of the United Parkinson's Disease Rating Scale (mUPDRS), where a trained nurse clinician scores 8 items: Right and left finger taps, right and left fist clench, right and left pronation-supination and right and left heel tap. The result is the sum of the ratings for the individual items, divided by the maximum possible score for the domain, then multiplied by 100. It varies from 0 (normal) to 5 (Can barely perform tasks) [31].

##### Rigidity

It is scored based on the passive movement of extremities when participants are in a relaxed sitting position. It is determined using a modified version of the motor portion of the mUPDRS and tests the neck and limb rigidity. The result is the sum of the ratings for the individual items, divided by the maximum possible score for the domain, then multiplied by 100. It varies from 0 (Absent) to 4 (Severe) [31].

##### Gait

The gait score is calculated using a modified version of the motor portion of the mUPDRS, and it is based on 6 items: Turning, posture, postural stability, arising from a chair, shuffling gait, and body bradykinesia/hypokinesia. Each item has its own score scale, and the gait score is the sum of the ratings for the individual items, divided by the maximum possible score for the domain, then multiplied by 100. The result varies from 0 to 10,0 with higher scores representing greater gait disturbance [31].

##### Tremor

Tremor score is determined using a modified version of the motor portion of the mUPDRS, and it is based on 7 items: left and right arm resting tremor, left and right leg resting tremor, left and right-hand postural tremor, and Chin/jaw resting tremor. Each item has its own score scale, and the tremor score is the sum of the ratings for the individual items, divided by the maximum possible score for the domain, then multiplied by 100. The result varies from 0 to 100, with higher scores reflecting more tremor [31].

##### Motor gait

Motor gait is a composite measure of gait created using 2 tests: walking and turning 360 degrees. Participants walk 8 feet and turn 360 degrees twice each, with time and steps recorded. These values are reciprocated, so larger values reflect faster performance and fewer steps. The two values from each trial are averaged to calculate the performance scores: walking time, walking steps, turning time, and turning steps. The 4 performance scores are each divided by the mean of the parent study at baseline. The 4 values are then averaged to yield a composite measure of motor gait [32].

##### Motor dexterity

Motor dexterity is a composite measure of 2 tests: the Purdue pegboard test and the finger-tapping test. It is measured bilaterally using the Purdue pegboard and an electronic tapper (Western Psychological Services, Los Angeles, CA). Participants perform each test twice per hand, and the average number of pegs inserted across four trials represents the performance score. The average of the 4 trials for the finger-tapping test represents a performance score, in number of finger taps. The performance scores are each divided by the mean of the parent study at baseline. The values are then averaged to yield a composite measure of motor dexterity [32].

##### Motor hand strength

Motor hand strength is a composite measure of 2 tests: grip and pinch strength. Grip and pinch strength are assessed bilaterally using Jamar hydraulic hand and pinch dynamometers (Lafayette Instruments, Lafayette, IN). Participants perform each test twice per hand, and the average of the four trials provides the performance scores for grip and pinch strength, measured in pounds of pressure. The performance scores are each divided by the sex-specific mean of the parent study at baseline. The values are then averaged to yield a composite measure of hand strength [32].

##### Global Parkinsonian score

The global Parkinsonian summary score is a continuous measure reflecting the severity of Parkinsonian signs, calculated as the average of four domains: tremor, rigidity, gait, and bradykinesia. Scores range from 0 to 100, with higher values indicating greater severity of Parkinsonism [31]. The longitudinal trajectory of this score was modeled using a linear mixed effects approach, with the square root-transformed global Parkinsonian score as the outcome. The person-specific random slope represents the individual rate of change in Parkinsonian symptoms over time, adjusted for baseline age, sex, and years of education. Time in this model is defined as years since the baseline interview [33].

##### Vascular pathologies

##### Arteriosclerosis

Arteriosclerosis was measured according to histological changes in the small vessels of the anterior basal ganglia. The alterations included intimal deterioration, smooth muscle degeneration, and fibrohyalinotic thickening of arterioles with consequent narrowing of the vascular lumen. Due to the lack of standard guidelines to grade the severity of arteriolosclerosis, it was evaluated with a semiquantitative grading system from 0 (none) to 3 (severe) [34].

##### Cerebral atherosclerosis

Large vessel cerebral atherosclerosis was evaluated by visual inspection after fixation, at the Circle of Willis at the base of the brain, and included evaluation of the vertebral, basilar, posterior cerebral, middle cerebral, and anterior cerebral arteries and their proximal branches. The severity was graded based on the extent of involvement of each artery, the number of arteries affected, and the degree of occlusion. The grading ranges from 0 (None or Possible) to 3 (severe) [35].

#### Cerebral amyloid angiopathy (CAA)

CAA levels were quantified across 4 neocortical regions (midfrontal, midtemporal, parietal, and calcarine cortices) utilizing an adaptation of a previously established protocol [36]: Paraffin-embedded sections were immunostained for β-amyloid, and for each region, meningeal and parenchymal vessels were assessed for β-amyloid deposition and scored from 0 (no deposition) to 4 (circumferential deposition over 75% of the total region). The CAA score for each area was determined by selecting the maximum score between meningeal and parenchymal CAA scores. These scores were then averaged across regions and summarized as a continuous measure of CAA pathology. For a semiquantitative summary, CAA scores were categorized from 0 (none) to 3 (severe) based on cutoffs established by a neuropathologist [36,37].

#### Cerebral infarctions (gross)

Gross cerebral infarctions were measured by the presence of one or more gross chronic cerebral infarctions, located anywhere in the brain, detected visually on fixed slabs and confirmed through neuropathologic evaluations [38,39].

#### Cerebral infarctions (micro)

The presence of one or more chronic microinfarcts was searched in at least nine regions (MFC, MTC, EC, HC, IPC, ACC, anterior basal ganglia, thalamus, and midbrain) of one hemisphere, confirmed through neuropathologic evaluations [39].

#### Disabilities

#### Basic activities of daily living (ADL)

It is measured with the Katz Activities of Daily Living Scale, which measures six basic physical abilities: walking across a small room, bathing, dressing, eating, getting from bed to chair, and toileting. Participants report whether they need help with ADLs, with responses dichotomized as 0 = no help and 1 = needs help or unable to perform. The composite score (0–6) sums the number of items requiring assistance, with higher scores indicating greater disability [40,41].

#### Instrumental activities of daily living (IADL)

It is a composite measure of disability using a sum of 8 items adapted from the Duke Older Americans Resources and Services project. The scale measures household management and self-care functions: telephone use, meal preparation, light housekeeping, heavy housekeeping, handling medications, handling finances, traveling within the community and shopping. Participants report their need for assistance with IADLs, with responses dichotomized as 0 = no help and 1 = needs help or cannot perform. The composite score (0–8) sums the items requiring assistance, with higher scores reflecting greater disability [41].

#### Mobility disability

The Rosow-Breslau scale measures the ability to do 3 activities: doing heavy work around the house, walking up and down stairs, and walking half a mile without help. Participants report their need for assistance with activities, with responses dichotomized as 0 = no help and 1 = needs help or cannot perform. The composite score (0–3) sums the items requiring assistance, with higher scores indicating greater disability [42].

#### Depressive disorder

#### Clinical diagnosis of depressive disorder

Clinical diagnosis of major depressive disorder rendered by a physician. The diagnosis is based on criteria of the Diagnostic and Statistical Manual of Mental Disorders, 3rd Edition, Revised (DSM-III-R), clinical interview with the participant, and review of responses to a series of questions adapted from the Diagnostic Interview Schedule. The score ranges from 1(highly probable) to 4 (not present) [43] and was adapted to a dichotomized version (1 as highly probable, and 0 as the other scores).

#### Modified CES-D

A modified, 10-item version of the Center for Epidemiologic Studies Depression scale (CES-D) was administered to assess depressive symptoms. The participants are asked whether or not they experienced each of ten symptoms, much of the time in the past week. The score is the total number of symptoms experienced divided by the number of items answered, then multiplied by 10; this way, the score is rescaled to account for missing data [44–46].

#### Genetics

#### APOE genotype

Genotyping of DNA extracted from either brain tissue or peripheral blood mononuclear cells was conducted using high-throughput sequencing, targeting codon 112 (position 3937) and codon 158 (position 4075) of exon 4 of the APOE gene [47]. A dichotomized score was created where samples were categorized into those with the APOEε4 allele and those without.

#### TOMM40 genotype (translocase of outer mitochondrial membrane, 40kD)

TOMM40’523 genotypes are determined by rs10524523 (chr19:44,899,792-44,899,826, human genome reference assembly GRCh38/hg38), a homopolymer length polymorphism (poly-T), at intron 6 of the TOMM40 gene [47,48]. Allele lengths are classified according to the number of poly-T repeats: short alleles (S) have fewer than 20 repeats, long alleles (L) have 20 to 29 repeats, and very long alleles (VL) have 30 or more repeats [47]. A dichotomized score was created where samples were categorized into those with at least one long allele and those without.

#### Demographics

##### Age at death

Age of death is calculated by subtracting date of birth from date of death and dividing the difference by days per year (365.25).

##### Years of education

The years of education variable is based on the number of years of regular school reported at baseline cognitive testing [49].

##### Sex

Sex is self-reported.

### Supplementary Figures

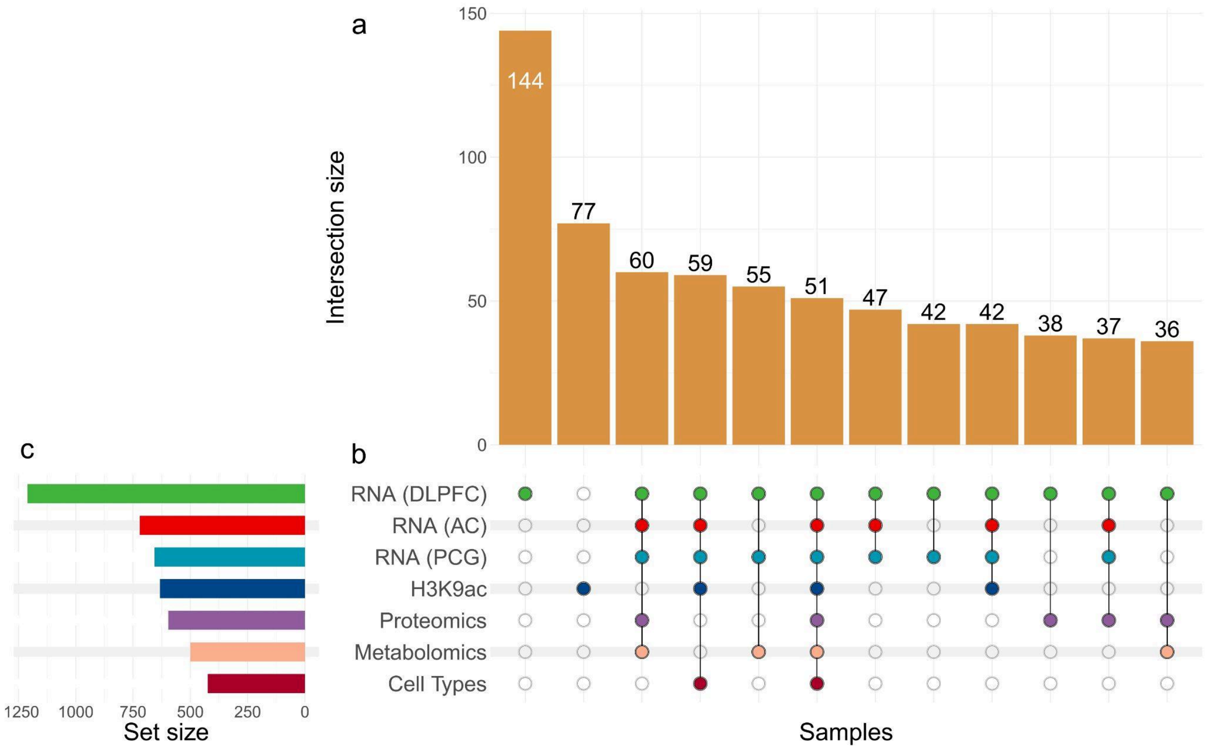

**Supplementary Figure S1. Overlap between omics datasets. (a)** intersection sizes, with the height of each bar corresponding to the number of samples shared among views. The number above each bar indicates the count of intersecting samples. **(b)** datasets included in each intersection, where filled circles represent participation in that intersection. **(c)** total set size for each dataset. For visualization, only intersections with n higher than 35 are shown.

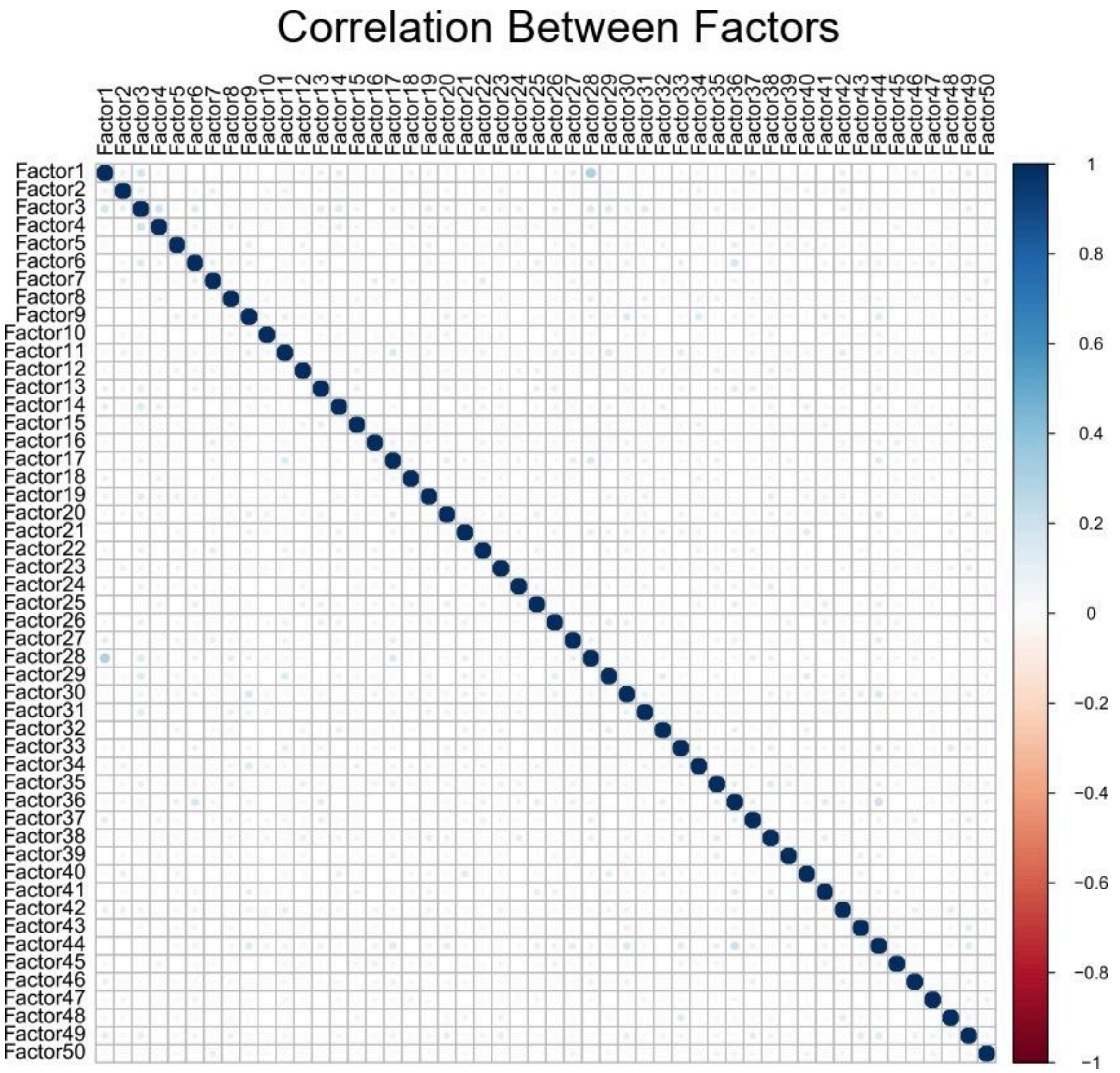

**Supplementary Figure S2. Correlation matrix between factors.** Color scale shows the correlation strength and direction from -1 (red) to 1 (blue) and the circle size represents the correlation strength as well. Pearson correlation was used to measure the relationship between the factors.

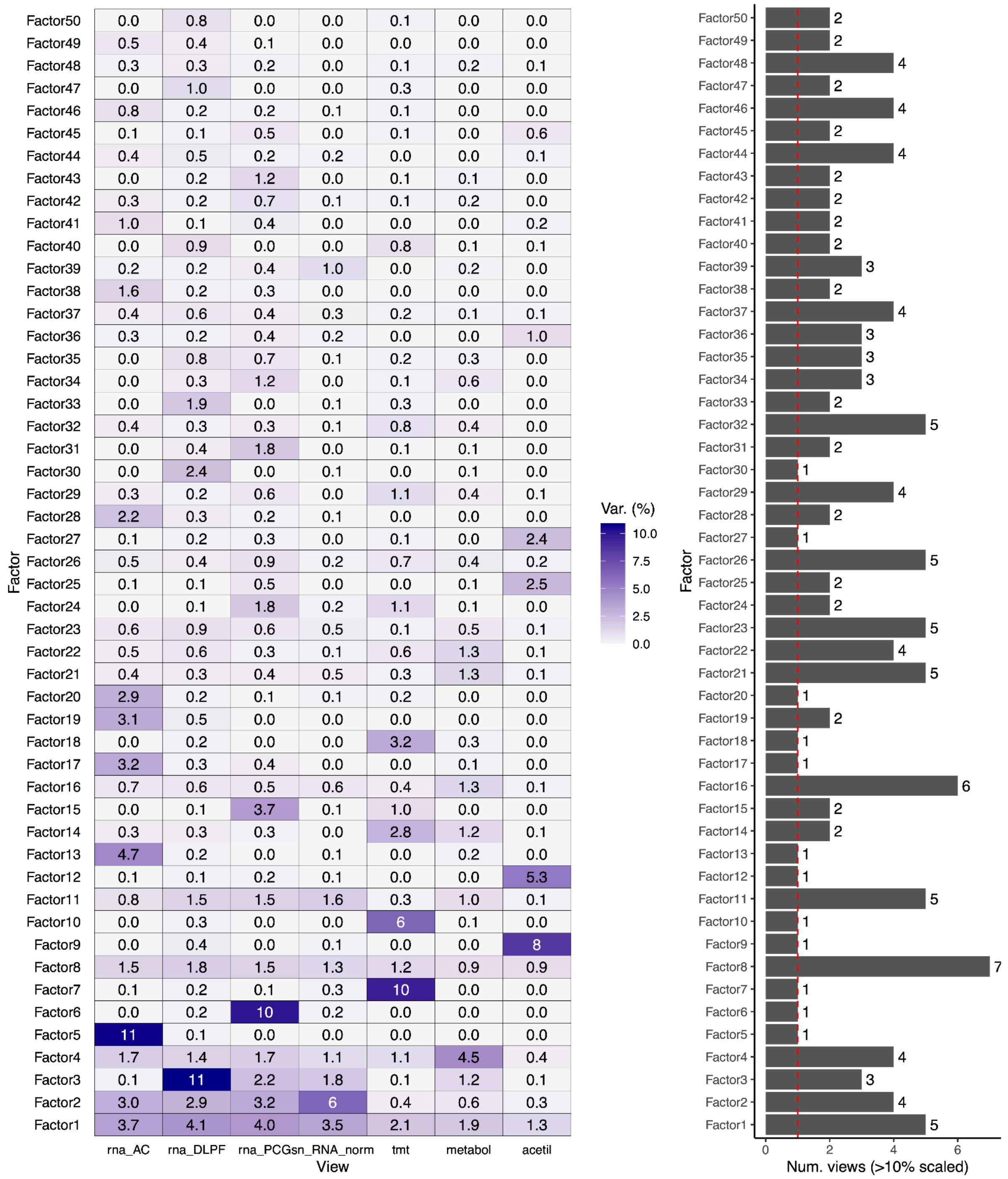

**Supplementary Figure S3. Variance explained by each view/factor. (a)** Heatmap showing the variance explained (shown as percentages) of all 50 MOFA factors and all seven views. **(b)** Number of views explaining more than 0.1 of the scaled variance by factor.

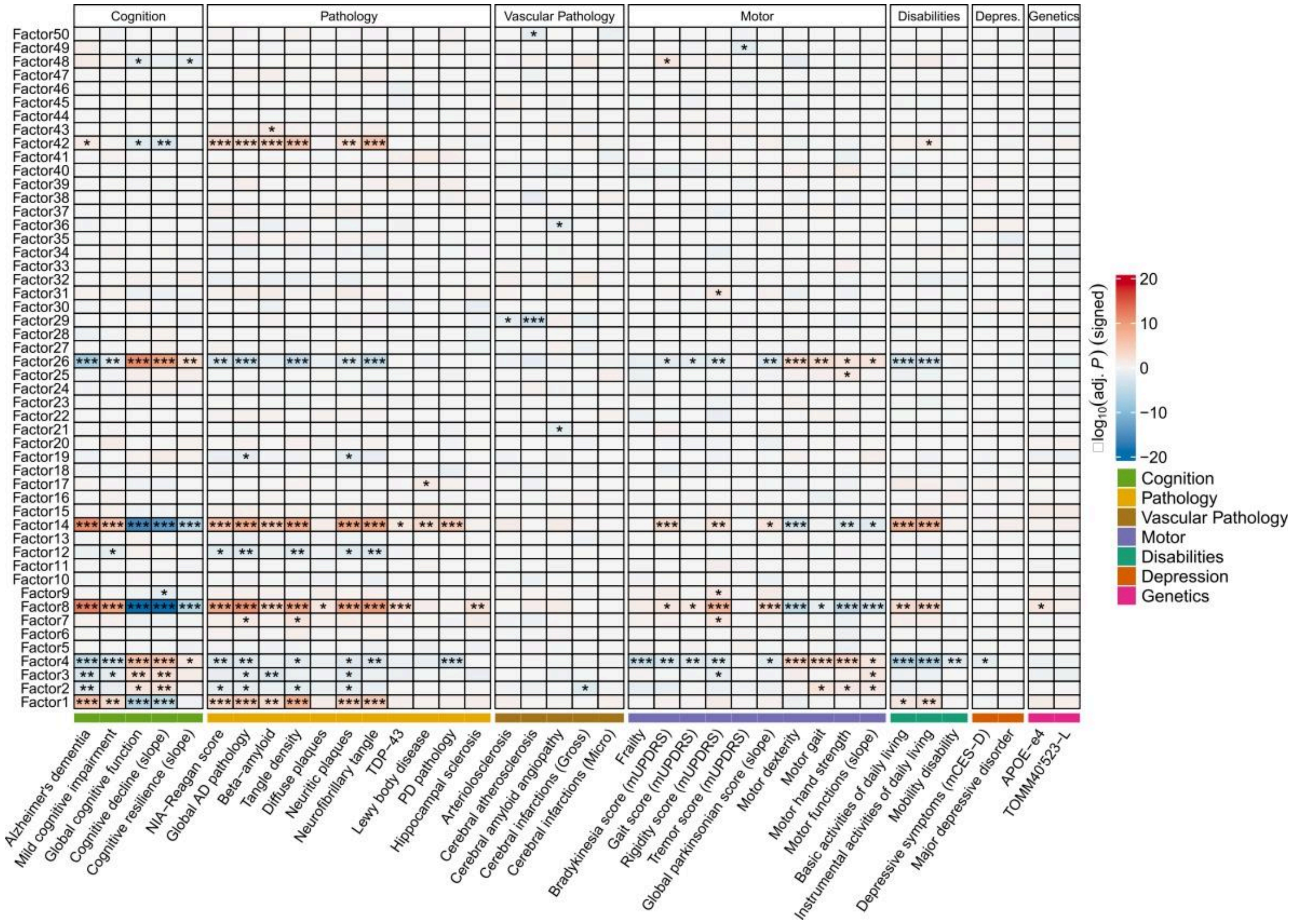

**Supplementary Figure S4. Relationship of all factors with covariates via regression analysis**. Results were controlled for sex, age, and years of education, with *P*-values corrected using the False Discovery Rate (FDR) method to address multiple comparisons. (*) = $-\log_{10}$ (adj. *P*) > 1.30103; (**) = $-\log_{10}$ (adj. *P*) > 2; (***) = $-\log_{10}$ (adj. *P*) > 3. The color scale shows the association effect size indicating direction and strength from negative (blue) to positive (red).

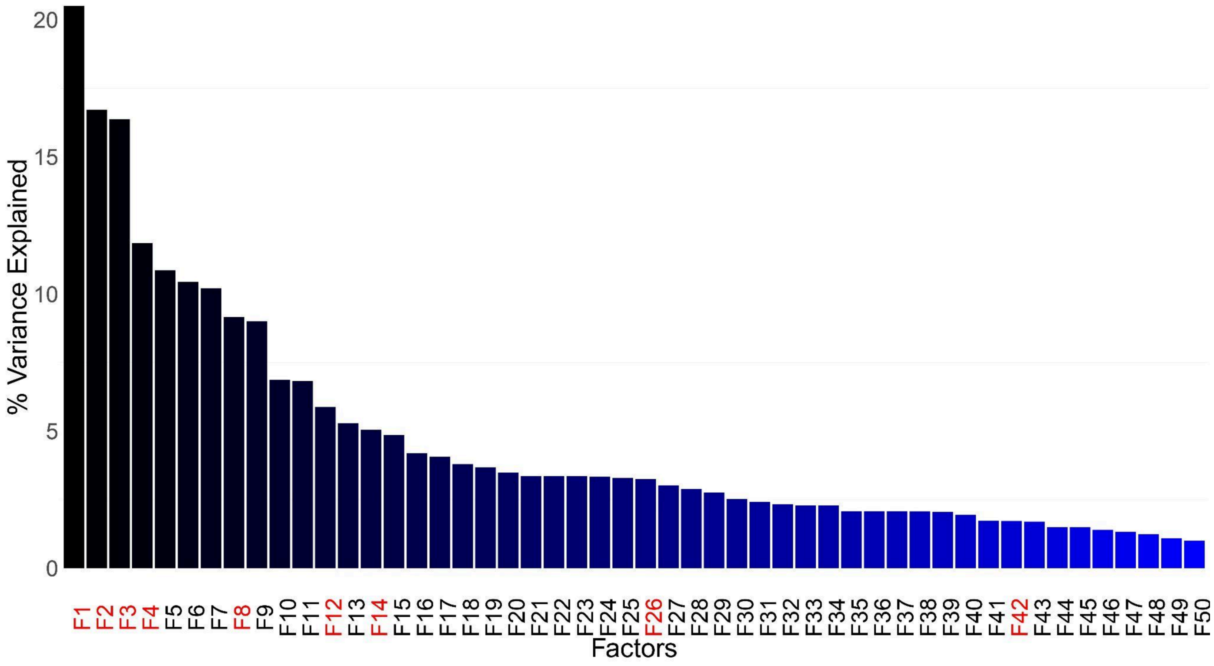

**Supplementary Figure S5. Percentage of variance explained for all factors**. Each bar height corresponds to the percentage of variance explained for each factor. Factors are represented in the X-axis, with the AD related factors highlighted in red.

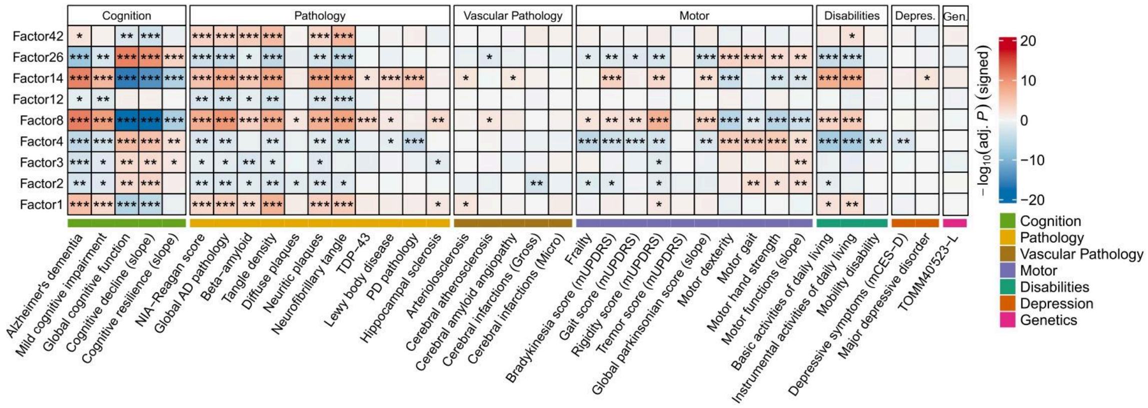

**Supplementary Figure S6. Relationship of factors with covariates via regression analysis**. Results were controlled for sex, age, years of education, and the presence of the allele APOE-ε4, with *P*-values corrected using the False Discovery Rate (FDR) method to address multiple comparisons. (*) = $-\log_{10}$ (adj. *P*) > 1.30103; (**) = $-\log_{10}$ (adj. *P*) > 2; (***) = $-\log_{10}$ (adj. *P*) > 3. The color scale shows the association effect size, indicating direction and strength from negative (blue) to positive (red).

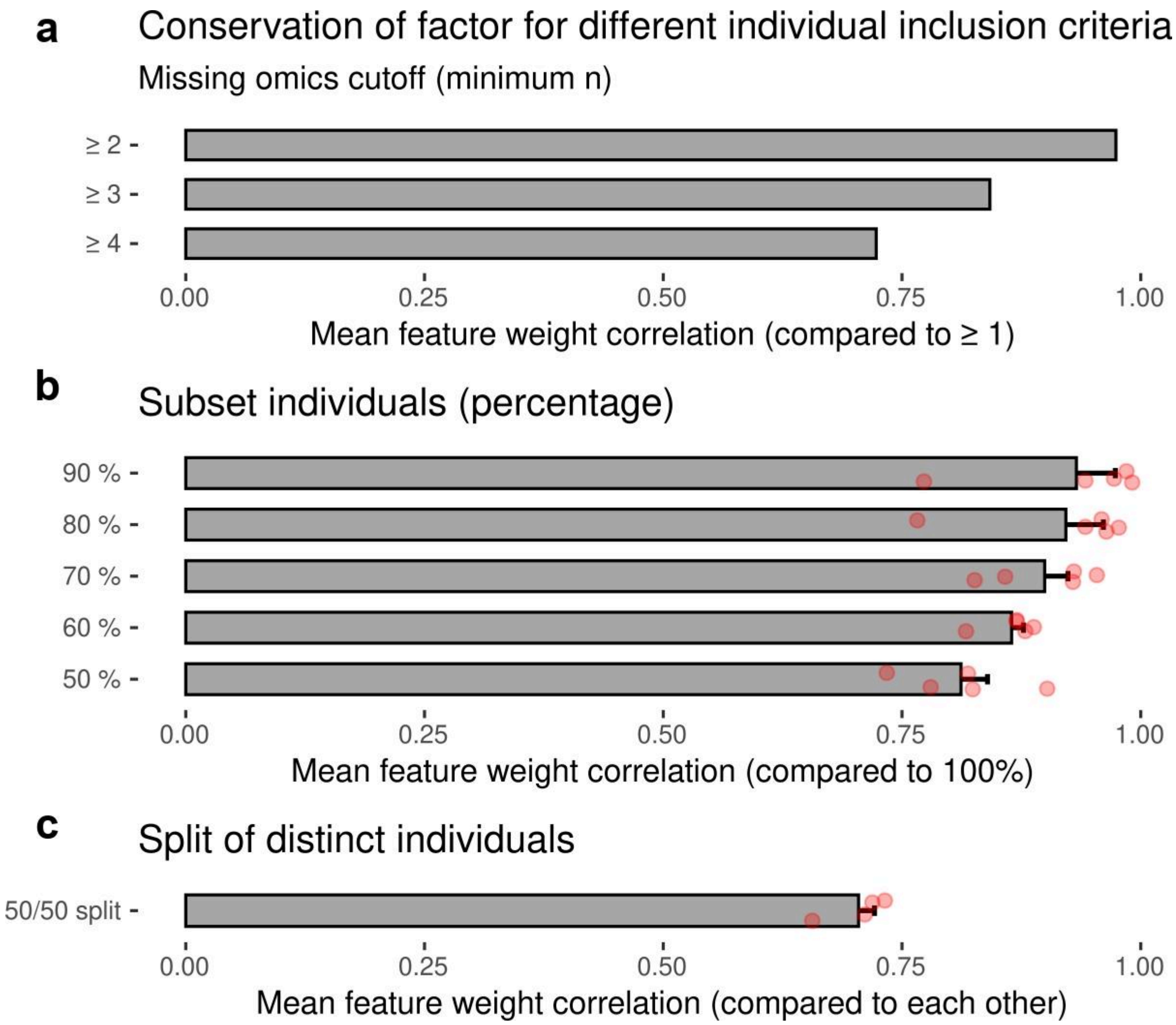

**Supplementary Figure S7. Robustness of MOFA feature weights across subsets of samples**. **(a)** Mean correlation of feature weights between the complete MOFA (trained on all samples) and alternative MOFA models trained on subsets of samples. Subset models included only individuals with at least *n* omics layers, thereby excluding samples with fewer data types. **(b)** Mean correlation of feature weights between the complete MOFA and MOFA models trained on random subsets of decreasing sample size. **(c)** Mean correlation of feature weights between two independent MOFA models trained on non-overlapping halves of the dataset. In all panels, bars indicate the mean correlation across factors.

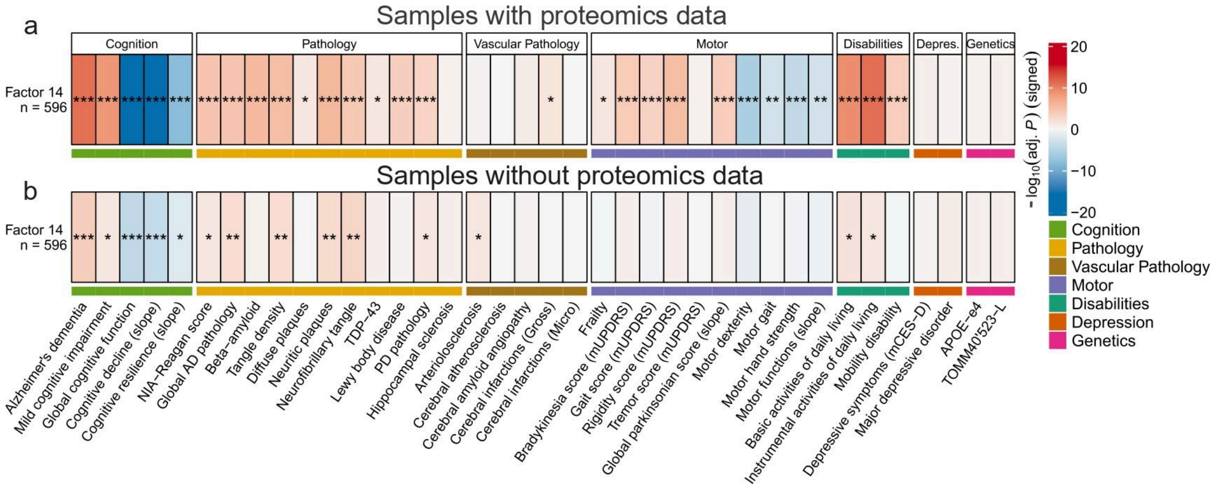

**Supplementary Figure S8. Relationship of Factor 14 with covariates in samples with and without proteomics data. (a)** Regression analyses restricted to samples with proteomics data. **(b)** Regression analyses restricted to samples without proteomics data. Results were controlled for sex, age, and years of education, with *P*-values corrected using the False Discovery Rate (FDR) method to address multiple comparisons. (*) = $-\log_{10}$ (adj. *P*) > 1.30103; (**) = $-\log_{10}$ (adj. *P*) > 2; (***) = $-\log_{10}$ (adj. *P*) > 3. The color scale shows the association effect size indicating direction and strength from negative (blue) to positive (red).

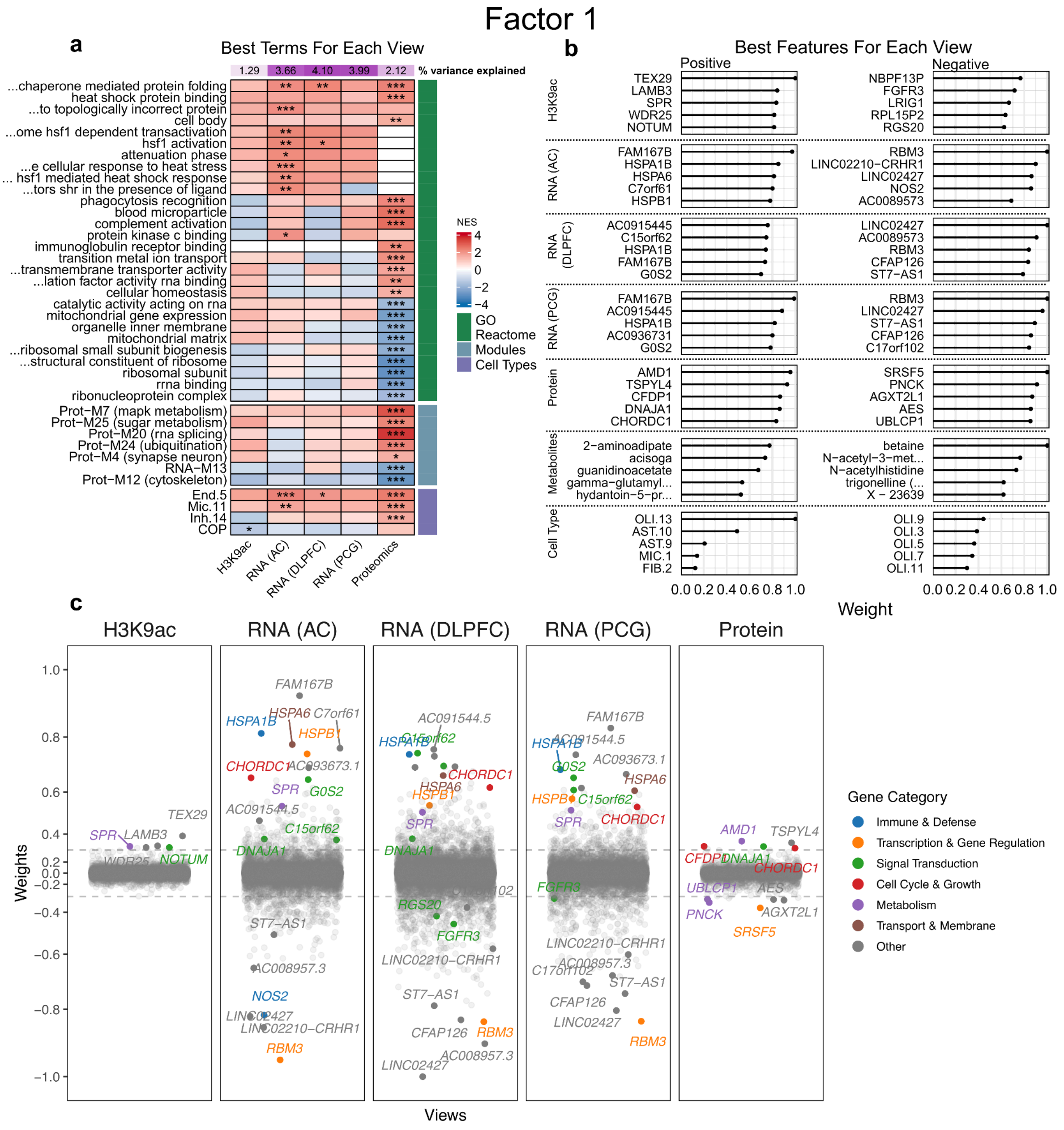

**Supplementary Figure S9. Characterization of factor 1 functional enrichment and feature contributions. (a)** Gene Set Enrichment Analysis of factor 1 showing the most significant terms for each view. *P*-values were corrected using *Benjamini-Hochberg* (BH) method to address multiple comparisons. Terms were selected based on the lowest *P* adjusted values for each view. 'GO' and 'Reactome' comprise terms from Gene Ontology and Reactome databases, respectively. 'Modules' include relevant co-expression protein modules from [50] and transcriptomic co-expression modules from [51]. The color scale shows the normalized enrichment score (NES) direction and strength from negative (blue) to positive (red). (*) = $-\log_{10}$(adj. $P < 0.05$); (**) = $-\log_{10}$(adj. $P < 0.01$); (***) = $-\log_{10}$(adj. $P < 0.001$). **(b)** The absolute value (scaled) of the five highest positive and negative features weights for each view. Positive weights indicate that the feature has higher levels in the cells with positive factor values, and vice-versa. Metabolites shown as 'X –' have not been characterized. **(c)** Jitter plot showing genes with weights in each *view* scaled across all *views* (cell types and metabolomics are not shown). Genes from selected pathways and with an absolute weight greater than 0.3 are highlighted (colored by category).

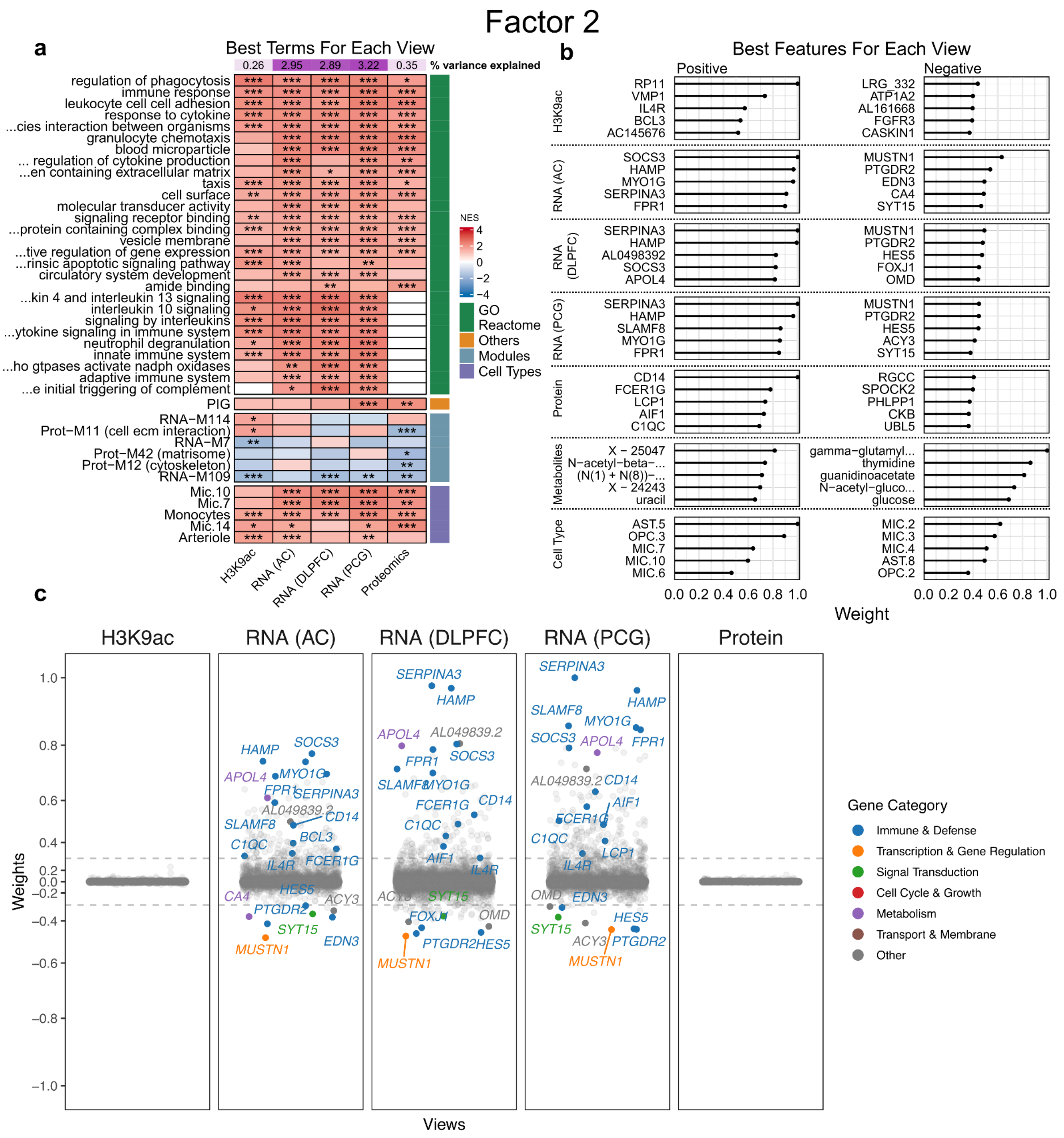

**Supplementary Figure S10. Characterization of factor 2 functional enrichment and feature contributions**. **(a)** Gene Set Enrichment Analysis of factor 2 showing the most significant terms for each view. *P*-values were corrected using *Benjamini-Hochberg* (BH) method to address multiple comparisons. Terms were selected based on the lowest *P* adjusted values for each view. 'GO' and 'Reactome' comprise terms from Gene Ontology and Reactome databases, respectively**.** 'Others' encompass a collection of pertinent AD-related modules. 'Modules' include relevant co-expression protein modules from [50] and transcriptomic co-expression modules from [51]. 'Cell Types' encompass subpopulations of cell lineages determined by [1]. The color scale shows the normalized enrichment score (NES) direction and strength from negative (blue) to positive (red). (*) = $-\log_{10}$ (adj. $P < 0.05$); (**) = $-\log_{10}$ (adj. $P < 0.01$); (***) = $-\log_{10}$ (adj. $P < 0.001$). **(b)** The absolute value (scaled) of the five highest positive and negative features weights for each view. Positive weights indicate that the feature has higher levels in the cells with positive factor values, and vice-versa. Metabolites shown as 'X –' have not been characterized. **(c)** Jitter plot showing genes with weights in each *view* scaled across all *views* (cell types and metabolomics are not shown). Genes from selected pathways and with an absolute weight greater than 0.3 are highlighted (colored by category).

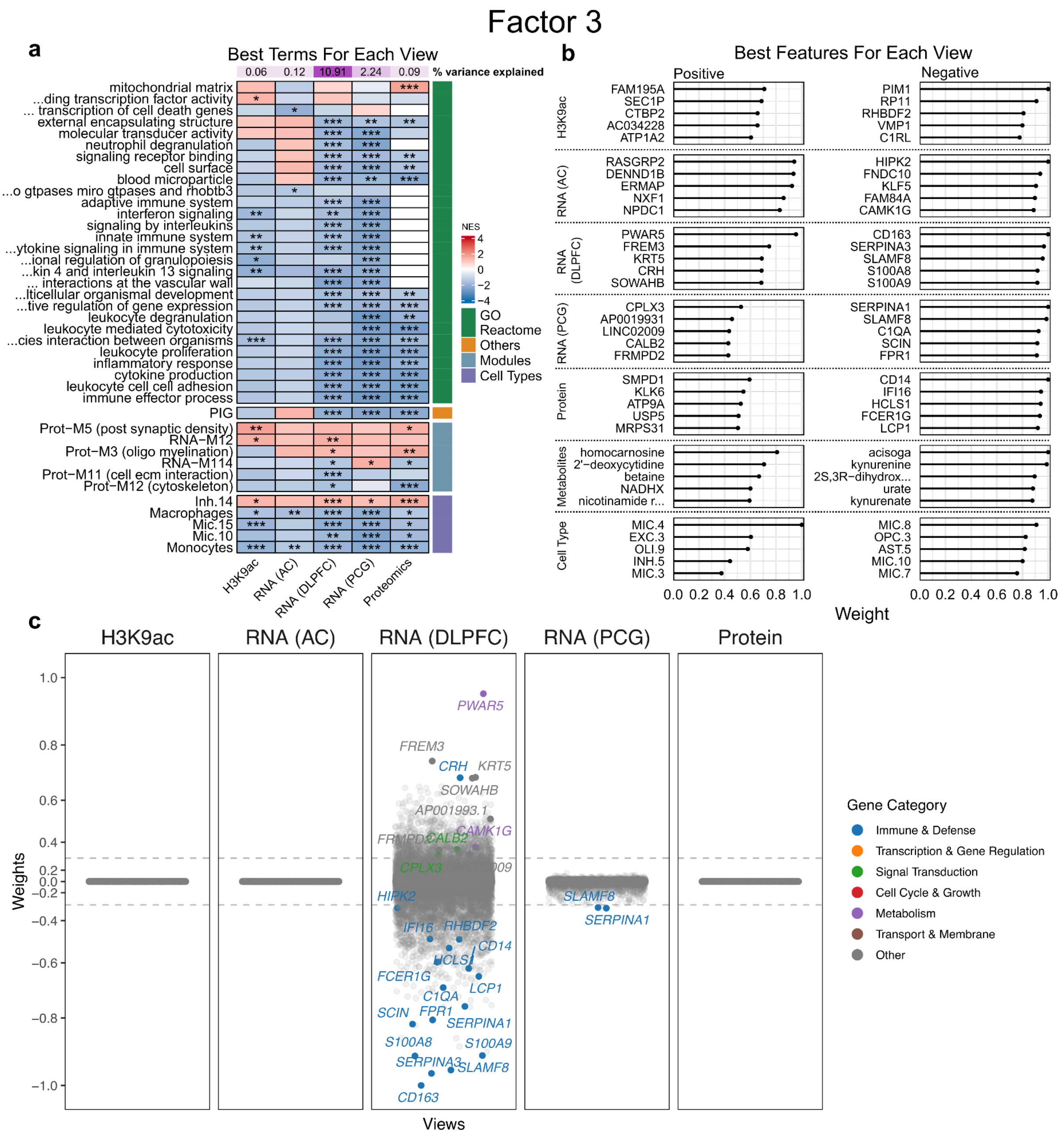

**Supplementary Figure S11. Characterization of factor 3 functional enrichment and feature contributions**. **(a)** Gene Set Enrichment Analysis of factor 3 showing the most significant terms for each view. *P*-values were corrected using *Benjamini-Hochberg* (BH) method to address multiple comparisons. Terms were selected based on the lowest *P* adjusted values for each view. 'GO' and 'Reactome' comprise terms from Gene Ontology and Reactome databases, respectively**.** 'Others' encompass a collection of pertinent AD-related modules. 'Modules' include relevant co-expression protein modules from [50] and transcriptomic co-expression modules from [51]. 'Cell Types' encompass subpopulations of cell lineages determined by [1]. The color scale shows the normalized enrichment score (NES) direction and strength from negative (blue) to positive (red). (*) = $-\log_{10}$ (adj. $P < 0.05$); (**) = $-\log_{10}$ (adj. $P < 0.01$); (***) = $-\log_{10}$ (adj. $P < 0.001$). **(b)** The absolute value (scaled) of the five highest positive and negative features weights for each view. Positive weights indicate that the feature has higher levels in the cells with positive factor values, and vice-versa. Metabolites shown as 'X –' have not been characterized. **(c)** Jitter plot showing genes with weights in each *view* scaled across all *views* (cell types and metabolomics are not shown). Genes from selected pathways and with an absolute weight greater than 0.3 are highlighted (colored by category).

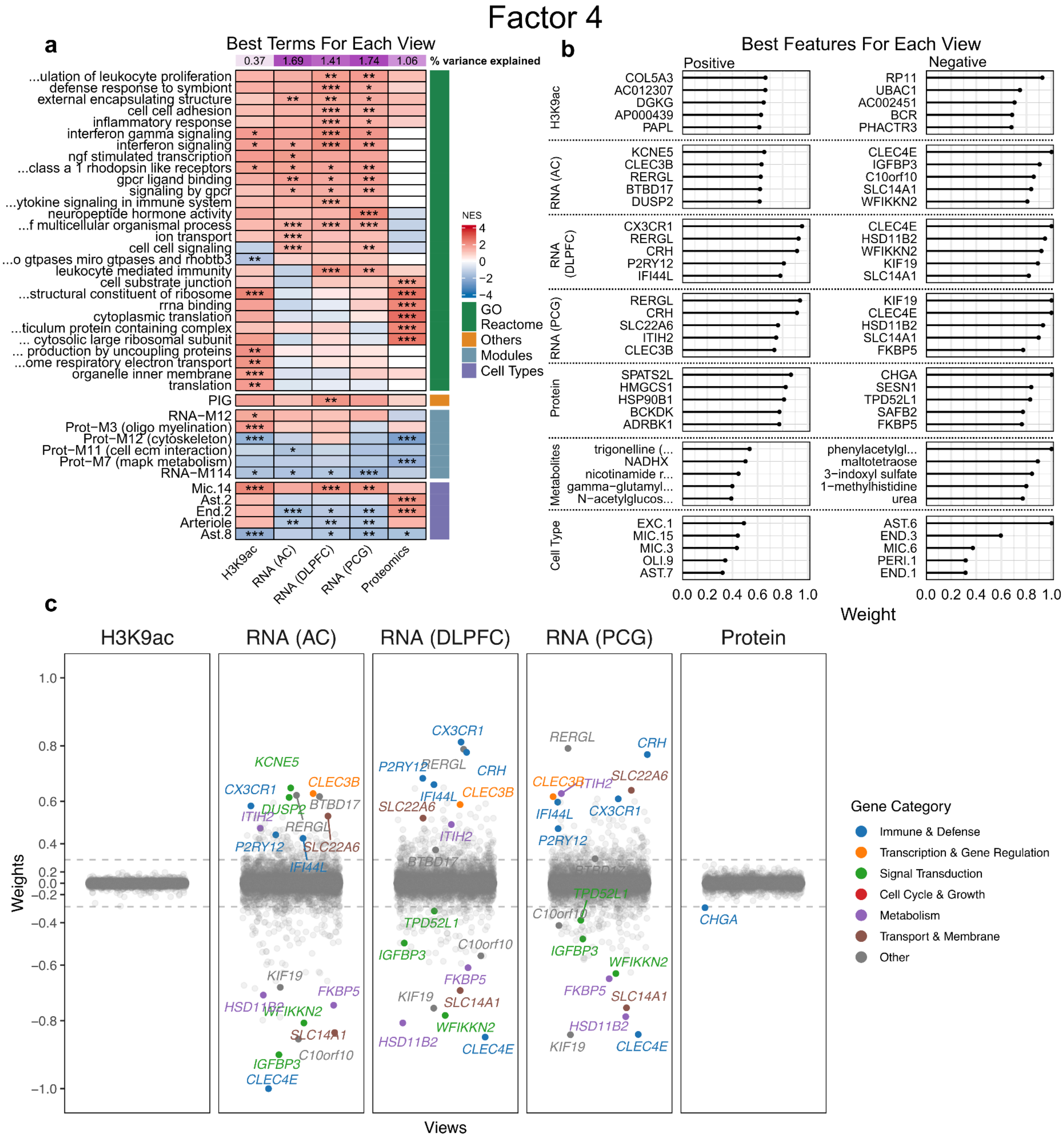

**Supplementary Figure S12. Characterization of factor 4 functional enrichment and feature contributions**. **(a)** Gene Set Enrichment Analysis of factor 4 showing the most significant terms for each view. *P*-values were corrected using *Benjamini-Hochberg* (BH) method to address multiple comparisons. Terms were selected based on the lowest *P* adjusted values for each view. 'GO' and 'Reactome' comprise terms from Gene Ontology and Reactome databases, respectively**.** 'Others' encompass a collection of pertinent AD-related modules. 'Modules' include relevant co-expression protein modules from [50] and transcriptomic co-expression modules from [51]. 'Cell Types' encompass subpopulations of cell lineages determined by [1]. The color scale shows the normalized enrichment score (NES) direction and strength from negative (blue) to positive (red). (*) = -log$_{10}$ (adj. $P < 0.05$); (**) = -log$_{10}$ (adj. $P < 0.01$); (***) = -log$_{10}$ (adj. $P < 0.001$). **(b)** The absolute value (scaled) of the five highest positive and negative features weights for each view. Positive weights indicate that the feature has higher levels in the cells with positive factor values, and vice-versa. Metabolites shown as 'X –' have not been characterized. **(c)** Jitter plot showing genes with weights in each *view* scaled across all *views* (cell types and metabolomics are not shown). Genes from selected pathways and with an absolute weight greater than 0.3 are highlighted (colored by category).

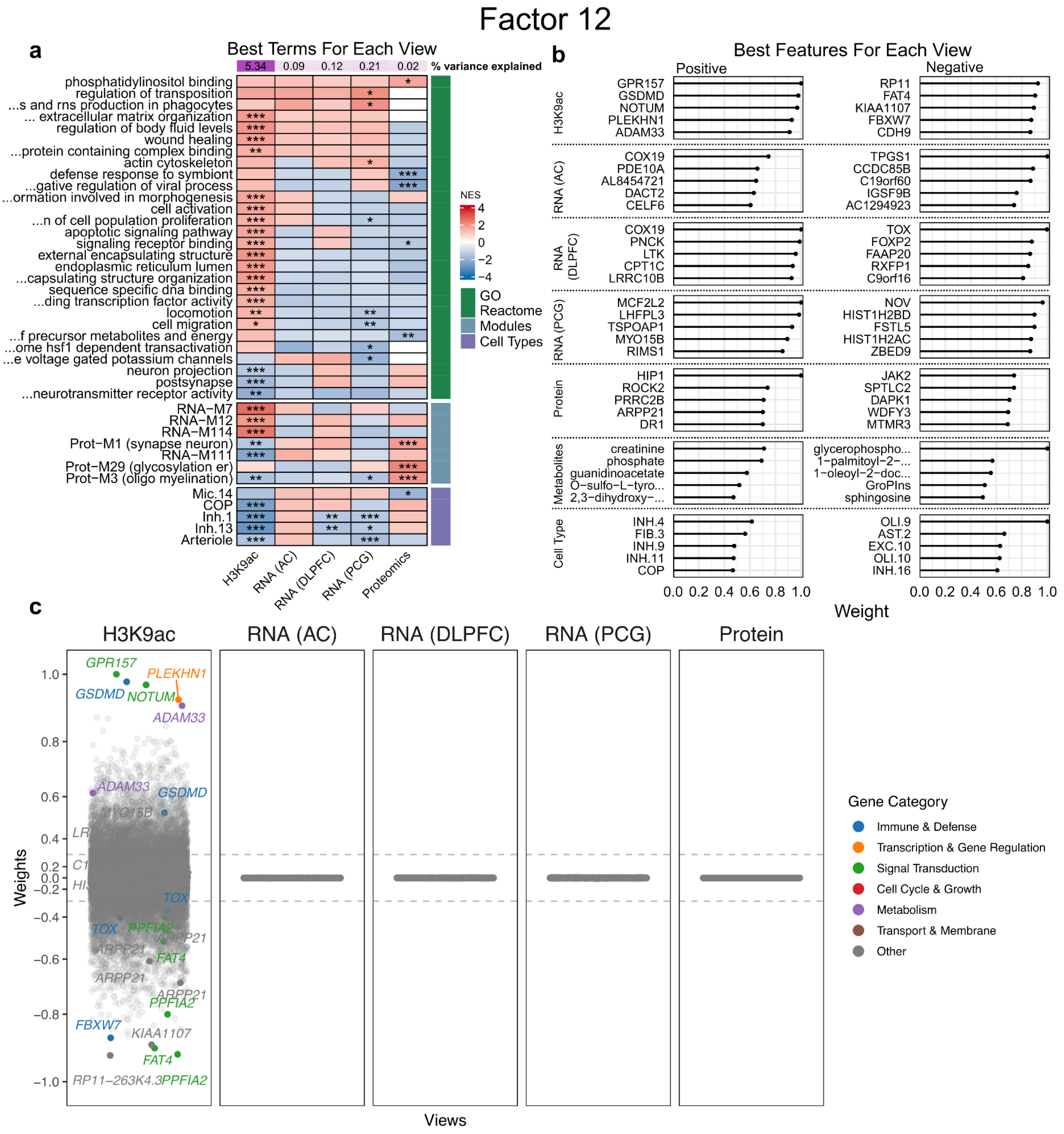

**Supplementary Figure S13. Characterization of factor 12 functional enrichment and feature contributions.** **(a)** Gene Set Enrichment Analysis of factor 12 showing the most significant terms for each view. *P*-values were corrected using *Benjamini-Hochberg* (BH) method to address multiple comparisons. Terms were selected based on the lowest *P* adjusted values for each view. 'GO' and 'Reactome' comprise terms from Gene Ontology and Reactome databases, respectively. 'Modules' include relevant co-expression protein modules from [50] and transcriptomic co-expression modules from [51]. 'Cell Types' encompass subpopulations of cell lineages determined by [1]. The color scale shows the normalized enrichment score (NES) direction and strength from negative (blue) to positive (red). (*) = -$\log_{10}$ (adj. $P$ < 0.05); (**) = -$\log_{10}$ (adj. $P$ < 0.01); (***) = -$\log_{10}$ (adj. $P$ < 0.001). **(b)** The absolute value (scaled) of the five highest positive and negative features weights for each view. Positive weights indicate that the feature has higher levels in the cells with positive factor values, and vice-versa. Metabolites shown as 'X –' have not been characterized. **(c)** Jitter plot showing genes with weights in each *view* scaled across all *views* (cell types and metabolomics are not shown). Genes from selected pathways and with an absolute weight greater than 0.3 are highlighted (colored by category).

Factor 14

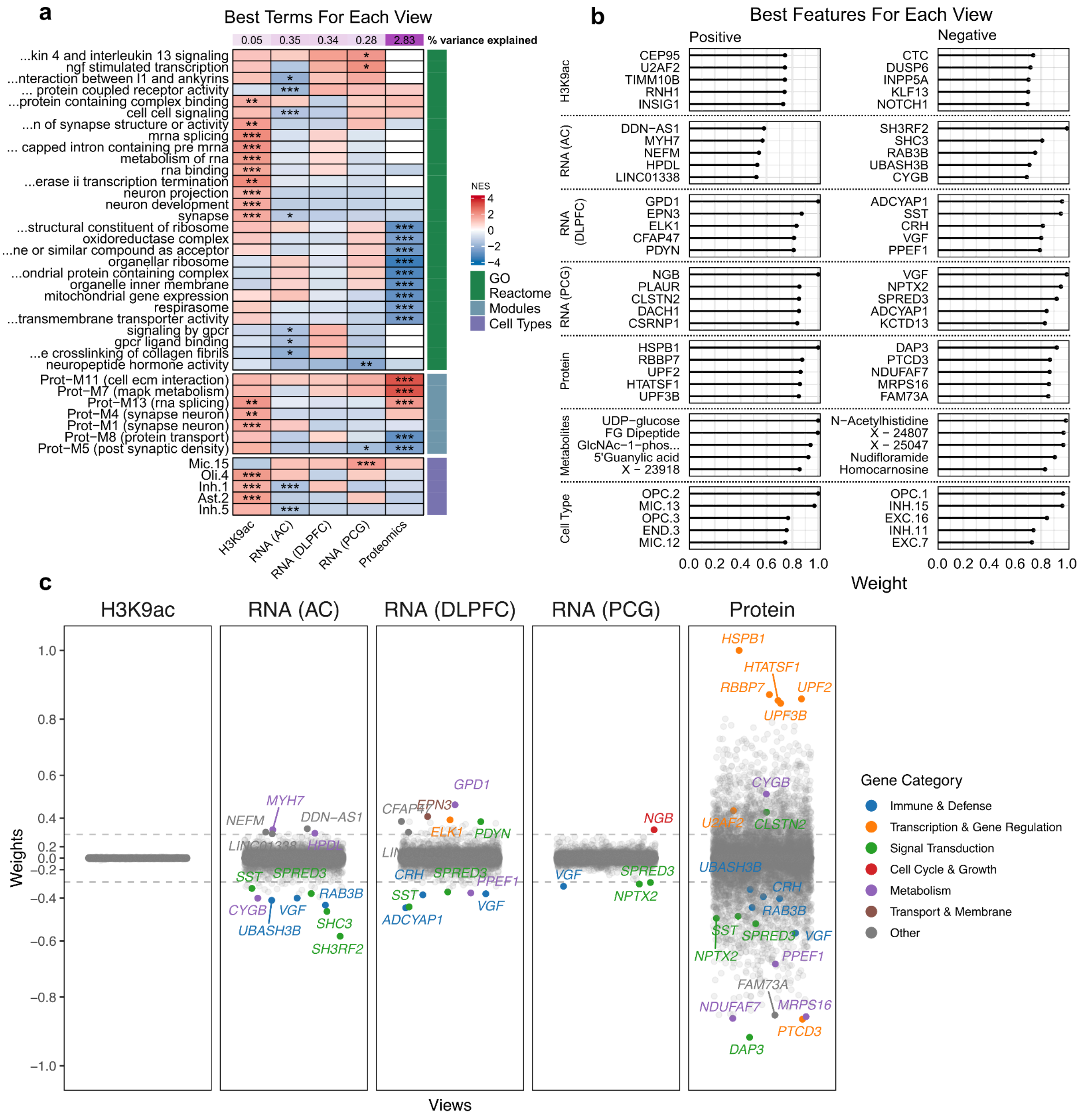

**Supplementary Figure S14. Characterization of factor 14 functional enrichment and feature contributions**. **(a)** Gene Set Enrichment Analysis of factor 14 showing the most significant terms for each view. *P*-values were corrected using *Benjamini-Hochberg* (BH) method to address multiple comparisons. Terms were selected based on the lowest *P* adjusted values for each view. 'GO' and 'Reactome' comprise terms from Gene Ontology and Reactome databases, respectively. 'Modules' include relevant co-expression protein modules from [50]. 'Cell Types' encompass subpopulations of cell lineages determined by [1]. The color scale shows the normalized enrichment score (NES) direction and strength from negative (blue) to positive (red). (*) = -$\log_{10}$ (adj. *P* < 0.05); (**) = -$\log_{10}$ (adj. *P* < 0.01); (***) = -$\log_{10}$ (adj. *P* < 0.001). **(b)** The absolute value (scaled) of the five highest positive and negative features weights for each view. Positive weights indicate that the feature has higher levels in the cells with positive factor values, and vice-versa. Metabolites shown as 'X –' have not been characterized. **(c)** Jitter plot showing genes with weights in each *view* scaled across all *views* (cell types and metabolomics are not shown). Genes from selected pathways and with an absolute weight greater than 0.3 are highlighted (colored by category).

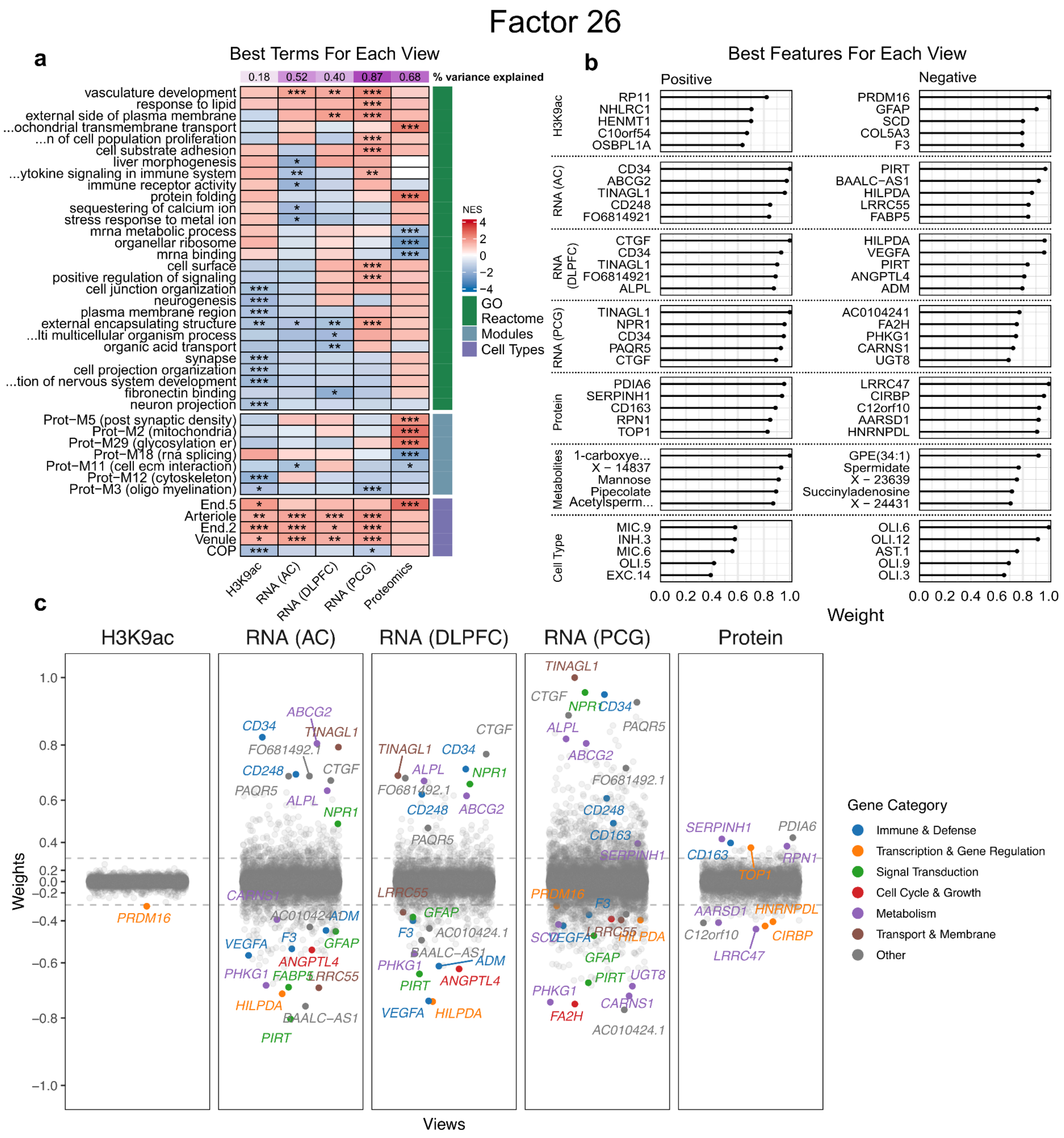

**Supplementary Figure S15. Characterization of factor 26 functional enrichment and feature contributions**. **(a)** Gene Set Enrichment Analysis of factor 26 showing the most significant terms for each view. *P*-values were corrected using *Benjamini-Hochberg* (BH) method to address multiple comparisons. Terms were selected based on the lowest *P* adjusted values for each view. 'GO' and 'Reactome' comprise terms from Gene Ontology and Reactome databases, respectively. 'Modules' include relevant co-expression protein modules from [50]. 'Cell Types' encompass subpopulations of cell lineages determined by [1]. The color scale shows the normalized enrichment score (NES) direction and strength from negative (blue) to positive (red). (*) = -$\log_{10}$ (adj. $P < 0.05$); (**) = -$\log_{10}$ (adj. $P < 0.01$); (***) = -$\log_{10}$ (adj. $P < 0.001$). **(b)** The absolute value (scaled) of the five highest positive and negative features weights for each view. Positive weights indicate that the feature has higher levels in the cells with positive factor values, and vice-versa. Metabolites shown as 'X –' have not been characterized. **(c)** Jitter plot showing genes with weights in each *view* scaled across all *views* (cell types and metabolomics are not shown). Genes from selected pathways and with an absolute weight greater than 0.3 are highlighted (colored by category).

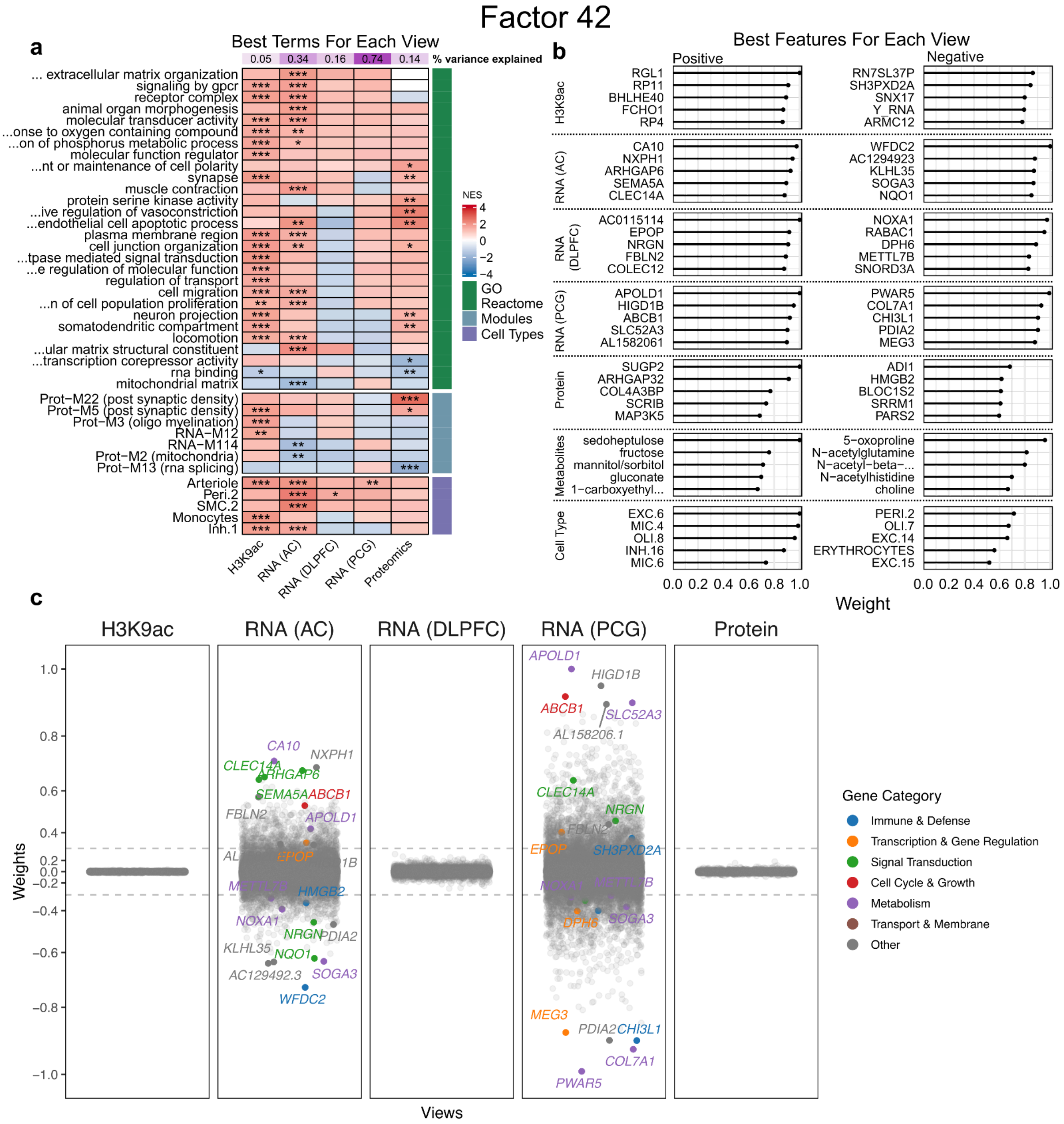

**Supplementary Figure S16. Characterization of factor 42 functional enrichment and feature contributions**. **(a)** Gene Set Enrichment Analysis of factor 42 showing the most significant terms for each view. *P*-values were corrected using *Benjamini-Hochberg* (BH) method to address multiple comparisons. Terms were selected based on the lowest *P* adjusted values for each view. 'GO' and 'Reactome' comprise terms from Gene Ontology and Reactome databases, respectively. 'Modules' include relevant co-expression protein modules from [50] and transcriptomic co-expression modules from [51]. 'Cell Types' encompass subpopulations of cell lineages determined by [1]. The color scale shows the normalized enrichment score (NES) direction and strength from negative (blue) to positive (red). (*) = $-\log_{10}$ (adj. $P < 0.05$); (**) = $-\log_{10}$ (adj. $P < 0.01$); (***) = $-\log_{10}$ (adj. $P < 0.001$). **(b)** The absolute value (scaled) of the five highest positive and negative features weights for each view. Positive weights indicate that the feature has higher levels in the cells with positive factor values, and vice-versa. Metabolites shown as 'X –' have not been characterized. **(c)** Jitter plot showing genes with weights in each *view* scaled across all *views* (cell types and metabolomics are not shown). Genes from selected pathways and with an absolute weight greater than 0.3 are highlighted (colored by category).

#### REFERENCES

1. Green, G. S. *et al.* Cellular communities reveal trajectories of brain ageing and Alzheimer's disease. *Nature* **633**, 634–645 (2024).
2. Chen, W.-T. *et al.* Spatial Transcriptomics and In Situ Sequencing to Study Alzheimer's Disease. *Cell* **182**, 976–991.e19 (2020).
3. Cho, S.-H. *et al.* CX3CR1 protein signaling modulates microglial activation and protects against plaque-independent cognitive deficits in a mouse model of Alzheimer disease. *J Biol Chem* **286**, 32713–32722 (2011).
4. Farid, M. M., Yang, X., Kuboyama, T. & Tohda, C. Trigonelline recovers memory function in Alzheimer's disease model mice: evidence of brain penetration and target molecule. *Sci Rep* **10**, 16424 (2020).
5. Wu, C.-Y. *et al.* Cognitive and Alzheimer's disease biomarker effects of oral nicotinamide riboside (NR) supplementation in older adults with subjective cognitive decline and mild cognitive impairment. *Alzheimers Dement (N Y)* **11**, e70023 (2025).
6. Blair, L. J. *et al.* Accelerated neurodegeneration through chaperone-mediated oligomerization of tau. *J Clin Invest* **123**, 4158–4169 (2013).
7. Watanabe, K. *et al.* The participation of insulin-like growth factor-binding protein 3 released by astrocytes in the pathology of Alzheimer's disease. *Mol Brain* **8**, 82 (2015).
8. Kim, S.-Y., Kim, M.-Y., Mo, J.-S. & Park, H.-S. Notch1 intracellular domain suppresses APP intracellular domain-Tip60-Fe65 complex mediated signaling through physical interaction. *Biochim. Biophys. Acta* **1773**, 736–746 (2007).
9. Saito, T. *et al.* Somatostatin regulates brain amyloid beta peptide Abeta42 through modulation of proteolytic degradation. *Nat. Med.* **11**, 434–439 (2005).
10. Zhang, J.-Z. *et al.* UDP-glucose sensing P2YR: A novel target for inflammation. *Neuropharmacology* **238**, 109655 (2023).
11. Beyer, L. *et al.* Amyloid-beta misfolding and GFAP predict risk of clinical Alzheimer's disease diagnosis within 17 years. *Alzheimers Dement* **19**, 1020–1028 (2023).
12. Kim, K. Y., Shin, K. Y. & Chang, K.-A. GFAP as a Potential Biomarker for Alzheimer's Disease: A Systematic Review and Meta-Analysis. *Cells* **12**, (2023).
13. Qi, F. *et al.* VEGF-A in serum protects against memory impairment in APP/PS1 transgenic mice by blocking neutrophil infiltration. *Mol Psychiatry* **28**, 4374–4389 (2023).
14. Batra, R. *et al.* The landscape of metabolic brain alterations in Alzheimer's disease. *Alzheimers. Dement.* (2022) doi:10.1002/alz.12714.
15. Bennett, D. A. *et al.* Neuropathology of older persons without cognitive impairment from two community-based studies. *Neurology* **66**, 1837–1844 (2006).
16. Consensus recommendations for the postmortem diagnosis of Alzheimer's disease. *Neurobiol. Aging* **18**, S1–S2 (1997).
17. Kapasi, A. *et al.* High-throughput digital quantification of Alzheimer disease pathology and associated infrastructure in large autopsy studies. *J. Neuropathol. Exp. Neurol.* **82**, 976–986 (2023).
18. Wilson, R. S., Arnold, S. E., Schneider, J. A., Tang, Y. & Bennett, D. A. The relationship between cerebral Alzheimer's disease pathology and odour identification in old age. *J. Neurol. Neurosurg. Psychiatry* **78**, 30–35 (2007).
19. Bennett, D. A. *et al.* Apolipoprotein E epsilon4 allele, AD pathology, and the clinical expression of Alzheimer's disease. *Neurology* **60**, 246–252 (2003).
20. Bennett, D. A., Schneider, J. A., Tang, Y., Arnold, S. E. & Wilson, R. S. The effect of social networks on the relation between Alzheimer's disease pathology and level of

cognitive function in old people: a longitudinal cohort study. *Lancet Neurol.* **5**, 406–412 (2006).
21. Nag, S. *et al.* TDP-43 pathology in anterior temporal pole cortex in aging and Alzheimer's disease. *Acta Neuropathol Commun* **6**, 33 (2018).
22. Schneider, J. A., Arvanitakis, Z., Bang, W. & Bennett, D. A. Mixed brain pathologies account for most dementia cases in community-dwelling older persons. *Neurology* **69**, 2197–2204 (2007).
23. Nag, S. *et al.* Hippocampal sclerosis and TDP-43 pathology in aging and Alzheimer disease. *Ann Neurol* **77**, 942–952 (2015).
24. Bennett, D. A. *et al.* Natural history of mild cognitive impairment in older persons. *Neurology* **59**, 198–205 (2002).
25. Bennett, D. A. *et al.* Decision rules guiding the clinical diagnosis of Alzheimer's disease in two community-based cohort studies compared to standard practice in a clinic-based cohort study. *Neuroepidemiology* **27**, 169–176 (2006).
26. Wilson, R. S. *et al.* Temporal course and pathologic basis of unawareness of memory loss in dementia. *Neurology* **85**, 984–991 (2015).
27. De Jager, P. L. *et al.* A genome-wide scan for common variants affecting the rate of age-related cognitive decline. *Neurobiol. Aging* **33**, 1017.e1–15 (2012).
28. Buchman, A. S., Boyle, P. A., Wilson, R. S., Tang, Y. & Bennett, D. A. Frailty is associated with incident Alzheimer's disease and cognitive decline in the elderly. *Psychosom. Med.* **69**, 483–489 (2007).
29. Buchman, A. S., Wilson, R. S., Bienias, J. L. & Bennett, D. A. Change in frailty and risk of death in older persons. *Exp. Aging Res.* **35**, 61–82 (2009).
30. Buchman, A. S. *et al.* Spinal motor neurons and motor function in older adults. *J. Neurol.* **266**, 174–182 (2019).
31. Buchman, A. S. *et al.* Nigral pathology and parkinsonian signs in elders without Parkinson disease. *Ann. Neurol.* **71**, 258–266 (2012).
32. Buchman, A. S., Wilson, R. S., Leurgans, S. E., Bennett, D. A. & Barnes, L. L. Change in motor function and adverse health outcomes in older African-Americans. *Exp Gerontol* **70**, 71–77 (2015).
33. Buchman, A. S. *et al.* Person-specific contributions of brain pathologies to progressive parkinsonism in older adults. *J. Gerontol. A Biol. Sci. Med. Sci.* **76**, 615–621 (2021).
34. Buchman, A. S., Leurgans, S. E., Nag, S., Bennett, D. A. & Schneider, J. A. Cerebrovascular disease pathology and parkinsonian signs in old age. *Stroke* **42**, 3183–3189 (2011).
35. Arvanitakis, Z. *et al.* The Relationship of Cerebral Vessel Pathology to Brain Microinfarcts. *Brain Pathol.* **27**, 77–85 (2017).
36. Love, S. *et al.* Development, appraisal, validation and implementation of a consensus protocol for the assessment of cerebral amyloid angiopathy in post-mortem brain tissue. *Am. J. Neurodegener. Dis.* **3**, 19–32 (2014).
37. Boyle, P. A. *et al.* Cerebral amyloid angiopathy and cognitive outcomes in community-based older persons. *Neurology* **85**, 1930–1936 (2015).
38. Schneider, J. A. *et al.* The apolipoprotein E epsilon4 allele increases the odds of chronic cerebral infarction [corrected] detected at autopsy in older persons. *Stroke* **36**, 954–959 (2005).
39. Arvanitakis, Z., Leurgans, S. E., Barnes, L. L., Bennett, D. A. & Schneider, J. A. Microinfarct pathology, dementia, and cognitive systems. *Stroke* **42**, 722–727 (2011).
40. Katz, S. & Akpom, C. A. A measure of primary sociobiological functions. *Int J Health*

*Serv* **6**, 493–508 (1976).
41. Boyle, P. A., Buchman, A. S., Wilson, R. S., Bienias, J. L. & Bennett, D. A. Physical activity is associated with incident disability in community-based older persons. *J Am Geriatr Soc* **55**, 195–201 (2007).
42. Buchman, A. S., Boyle, P. A., Leurgans, S. E., Evans, D. A. & Bennett, D. A. Pulmonary function, muscle strength, and incident mobility disability in elders. *Proc Am Thorac Soc* **6**, 581–587 (2009).
43. Bennett, D. A., Wilson, R. S., Schneider, J. A., Bienias, J. L. & Arnold, S. E. Cerebral infarctions and the relationship of depression symptoms to level of cognitive functioning in older persons. *Am. J. Geriatr. Psychiatry* **12**, 211–219 (2004).
44. Kohout, F. J., Berkman, L. F., Evans, D. A. & Cornoni-Huntley, J. Two shorter forms of the CES-D (Center for Epidemiological Studies Depression) depression symptoms index. *J. Aging Health* **5**, 179–193 (1993).
45. Wilson, R. S. *et al.* Depressive symptoms, cognitive decline, and risk of AD in older persons. *Neurology* **59**, 364–370 (2002).
46. Wilson, R. S. *et al.* Clinical-pathologic study of depressive symptoms and cognitive decline in old age. *Neurology* **83**, 702–709 (2014).
47. Yu, L. *et al.* ’523 variant and cognitive decline in older persons with ε3/3 genotype. *Neurology* **88**, 661–668 (2017).
48. Roses, A. D. *et al.* A TOMM40 variable-length polymorphism predicts the age of late-onset Alzheimer’s disease. *Pharmacogenomics J.* **10**, 375–384 (2010).
49. Bennett, D. A., Schneider, J. A., Wilson, R. S., Bienias, J. L. & Arnold, S. E. Education modifies the association of amyloid but not tangles with cognitive function. *Neurology* **65**, 953–955 (2005).
50. Johnson, E. C. B. *et al.* Large-scale deep multi-layer analysis of Alzheimer’s disease brain reveals strong proteomic disease-related changes not observed at the RNA level. *Nat Neurosci* **25**, 213–225 (2022).
51. Mostafavi, S. *et al.* A molecular network of the aging human brain provides insights into the pathology and cognitive decline of Alzheimer’s disease. *Nat Neurosci* **21**, 811–819 (2018).
